# HIV-Induced Sialoglycans on Infected Cells Promote Immune Evasion from Myeloid Cell-Mediated Killing

**DOI:** 10.1101/2025.02.21.639489

**Authors:** Shalini Singh, S M Shamsul Islam, Rui Liu, Opeyemi S. Adeniji, Leila B. Giron, Pratima Saini, Ali Danesh, Paul W Denton, Brad Jones, Han Xiao, Mohamed Abdel-Mohsen

## Abstract

Sialic acid-containing glycans (sialoglycans) on pathological cells interact with Siglecs, glyco-immune checkpoint receptors expressed on myeloid cells such as monocytes and neutrophils. This interaction suppresses the cytotoxic functions of these immune cells. We show that HIV infection reprograms the glycosylation machinery of infected cells to increase the expression of specific sialoglycan ligands for Siglecs-3, -7, and -9. These ligands engage Siglecs on myeloid cells, impairing their ability to target HIV-infected cells. Selective disruption of these interactions using 10-1074-Sia, an HIV-specific antibody conjugated to sialidase—an enzyme that removes sialic acids—significantly enhances monocyte- and neutrophil-mediated killing of HIV-infected cells in autologous assays. Treatment with 10-1074-Sia in humanized mice infected with HIV reduces viral load and decreases inflammation. These findings reveal a novel immune evasion mechanism exploited by HIV to evade myeloid cell immune surveillance and highlight the potential of targeting sialoglycan-Siglec interactions to improve immune clearance of HIV-infected cells.

## INTRODUCTION

HIV-infected cells employ various strategies to evade immune surveillance ^1, 2, 3, 4, 5^. Understanding the mechanisms of this immune evasion is crucial, as many therapeutic approaches aiming to eradicate HIV-infected cells depend on the immune system’s capacity to target and eliminate these cells ^6, 7, 8, 9^. Therefore, elucidating how HIV enhances its resistance to immune detection is essential for improving the efficacy of current eradication strategies and developing novel immunotherapeutic approaches to enhance immune-mediated clearance of infected cells.

A recently identified mechanism of immune evasion involves the alteration of surface glycans on pathological cells, including cancer cells, enabling them to escape innate immune surveillance by engaging glyco-immune checkpoints on immune cells ^10, 11, 12^. Classical immune checkpoints function through protein-protein interactions; for example, PD-1 on CD8^+^ T cells binds to PD-L1 on pathological cells, transmitting an inhibitory signal that suppresses immune responses. However, another class of immune checkpoints, known as glyco-immune checkpoints, recognizes glycans instead of proteins to mediate their inhibitory signals to immune functions ^13^. One prominent example of glyco-immune checkpoints is the Siglecs (sialic acid-binding immunoglobulin-like lectins), a family of receptors expressed on various innate immune cells, particularly those of myeloid origin, such as monocytes, macrophages, and neutrophils, as well as on natural killer (NK) cells ^14, 15^.

When pathological cells express elevated levels of sialic acid-containing ligands (sialoglycans), these ligands engage Siglec receptors on immune cells, leading to the activation of immunoreceptor tyrosine-based inhibitory motifs (ITIMs). This engagement triggers downstream signaling through the recruitment of phosphatases such as SHP-1 and SHP-2, which dephosphorylate signaling molecules critical for immune cell activation, including PI3K, Akt, MAPK, NF-κB. As a result, immune cell activity is suppressed, enabling pathological cells to evade innate immune surveillance ^15, 16, 17, 18^. In addition to cancer cells, recent studies have shown that some viral infections, such as Hepatitis B virus infection, may utilize the same mechanism to evade immune surveillance ^19^. The recognition of aberrant glycosylation patterns in cancer and other pathological cells has recently spurred significant interest in developing strategies to disrupt these glycan-mediated interactions, thereby enhancing immune-mediated clearance of these pathological cells ^10, 11, 19, 20, 21, 22, 23, 24, 25, 26, 27, 28, 29, 30^.

In this study, we investigate whether HIV infection alters the cell-surface glycosylation of infected cells to increase the expression of specific Siglec ligands and evade immune surveillance by innate immune cells, particularly myeloid cells. We show that HIV reprograms the glycosylation machinery of its host cells, leading to the upregulation of sialoglycan ligands for specific Siglecs, including Siglec-3, Siglec-7, and Siglec-9, but not Siglec-10. This glycan remodeling facilitates the evasion of these infected cells from myeloid cell-mediated cytotoxicity. After identifying this novel mechanism of immune evasion employed by HIV-infected cells, we further explored the potential of disrupting the sialic acid-Siglec interactions to enhance the susceptibility of HIV-infected cells to immune-mediated clearance, both *in vitro* and *in vivo*.

## RESULTS

### HIV infection induces the expression of sialoglycan ligands for Siglec-3, -7, and -9, but not Siglec-10, on the surface of primary CD4^+^ T cells

We first investigated whether HIV infection of primary CD4^+^ T cells induces the expression of sialoglycan ligands for specific Siglecs known to impact immune recognition, including Siglec-3, -7, -9, and -10 ^10, 19, 29, 30, 31, 32^. To this end, CD4^+^ T cells isolated from HIV-negative individuals were activated using αCD3/αCD28 beads and simultaneously infected with HIV_TYBE_ in the presence or absence of benzyl 2-acetamido-2-deoxy-α-D-galactopyranoside (BADG), an inhibitor of *O*-glycosylation. HIV infection of activated CD4^+^ T cells resulted in high levels of HIV p24^+^ cells, and BADG treatment did not affect the rate of this infection (**Fig. 1a**). To measure the levels of cell-surface Siglec ligands, we used recombinant Siglec-Fc chimera proteins and a BV421-conjugated secondary antibody, analyzed via flow cytometry. HIV_TYBE_ infection significantly induced the expression of Siglec-3 ligands on the surface of activated CD4^+^ T cells, an effect inhibited by BADG (**Fig. 1b**). Similar results were observed for the cell-surface expression of Siglec-7 (**Fig. 1c**) and Siglec-9 ligands (**Fig. 1d**). However, HIV infection did not affect the expression of Siglec-10 ligands (**Fig. 1e**), while BADG treatment reduced the expression of these ligands, as expected.

**Figure 1.**
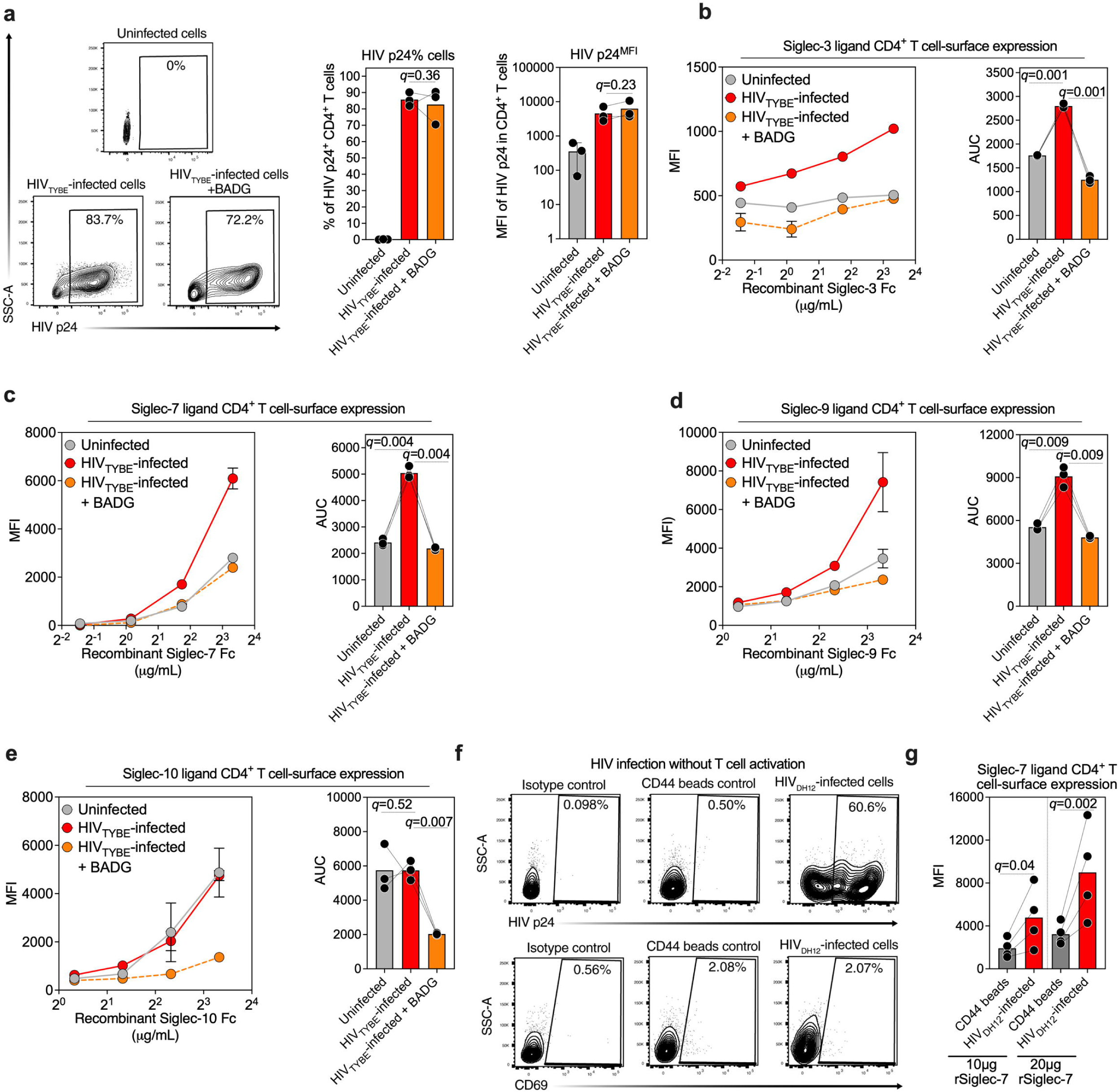
HIV infection induces the cell-surface expression of sialoglycan ligands for Siglec-3, -7, and -9, but not Siglec-10, on primary CD4^+^ T cells. **(a)** Primary CD4^+^ T cells isolated from HIV-negative donors were activated with anti-CD3/CD28 Dynabeads and concurrently infected with a primary HIV isolate (HIV_TYBE_) in the presence or absence of the glycosylation inhibitor BADG (2.5 mM). Intracellular p24 expression was assessed by flow cytometry 72 hours post-infection. Statistical analysis was performed using paired one-way ANOVA, corrected by the two-stage step-up procedure of Benjamini, Krieger, and Yekutieli. (**b-e)** HIV-uninfected as well as HIV_TYBE_-infected primary CD4+ T cells cultured in the presence or absence of BADG were stained with increasing concentrations of recombinant Siglec-3 Fc **(b**), Siglec-7 Fc **(c),** Siglec-9 Fc (d), and Siglec-10 Fc (e) 72 hours post-infection. The left panels show changes in Siglec ligand expression, while the right panels depict the area under the curve (AUC) for recombinant Siglec Fc binding. Error bars represent the mean ± standard deviation. Statistical analysis was performed using paired one-way ANOVA, corrected by the two-stage step-up procedure of Benjamini, Krieger, and Yekutieli. (**f**) Non-activated primary CD4^+^ T cells from HIV-negaive donors were infected with HIV_DH12_ using CD44 microbeads and stained for intracellular p24 expression and CD69 surface expression as a marker of T cell activation. **(g)** Non-activated CD4^+^ T cells exposed to CD44 microbeads or exposed and infected with HIV_DH12_ were stained with 10 or 20 µg of recombinant Siglec-7 Fc 48 hours post-infection. Statistical analyses were performed using paired one-way ANOVA, corrected by the two-stage step-up procedure of Benjamini, Krieger, and Yekutieli. N = 3–4.

To isolate the effects of T cell activation from the HIV-mediated induction of sialoglycans on the surface of infected cells, and to examine whether different viral strains elicit similar effects, we utilized CD44 microbeads to achieve high levels of HIV infection by HIV_DH12_ strain in non-activated T cells. Primary non-activated CD4^+^ T cells from several HIV-negative donors were infected with HIV_DH12_ in the presence of CD44 microbeads ^33^. Infection resulted in a high frequency of HIV p24^+^ cells (**Fig. 1f**) without increasing T cell activation, as measured by CD69 expression (**Fig. 1f**). Consistent with our findings using activated T cells and HIV_TYBE_, infection of non-activated T cells with HIV_DH12_ also induced Siglec-7 ligand expression compared to cells treated with CD44 microbeads alone (**Fig. 1g)**. These results suggest that HIV infection induces the expression of sialoglycan ligands for specific Siglecs, including Siglec-3, -7, and -9, on CD4^+^ T cells. This induction of Siglec ligands is independent of its activation state and may facilitate immune evasion by infected cells, leveraging Siglec-mediated immune suppression.

### HIV infection alters the glycosylation machinery of CD4^+^ T cells

We next investigated whether HIV infection alters the glycosylation machinery of CD4^+^ T cells, potentially explaining the induction of sialoglycan ligands for Siglecs on the surface of infected cells. To minimize the confounding effects of global T cell activation on glycosylation pathways, we infected non-activated CD4^+^ T cells from three HIV-negative donors with HIV_DH12_ in the presence of CD44 microbeads, which facilitate T cell infection without requiring activation (**Fig. 1f**). After 16 hours, cells were washed to remove unbound virus, cultured for an additional 24 hours, and subjected to global transcriptomic analysis using RNA-seq (**Fig. 2a**).

**Figure 2.**
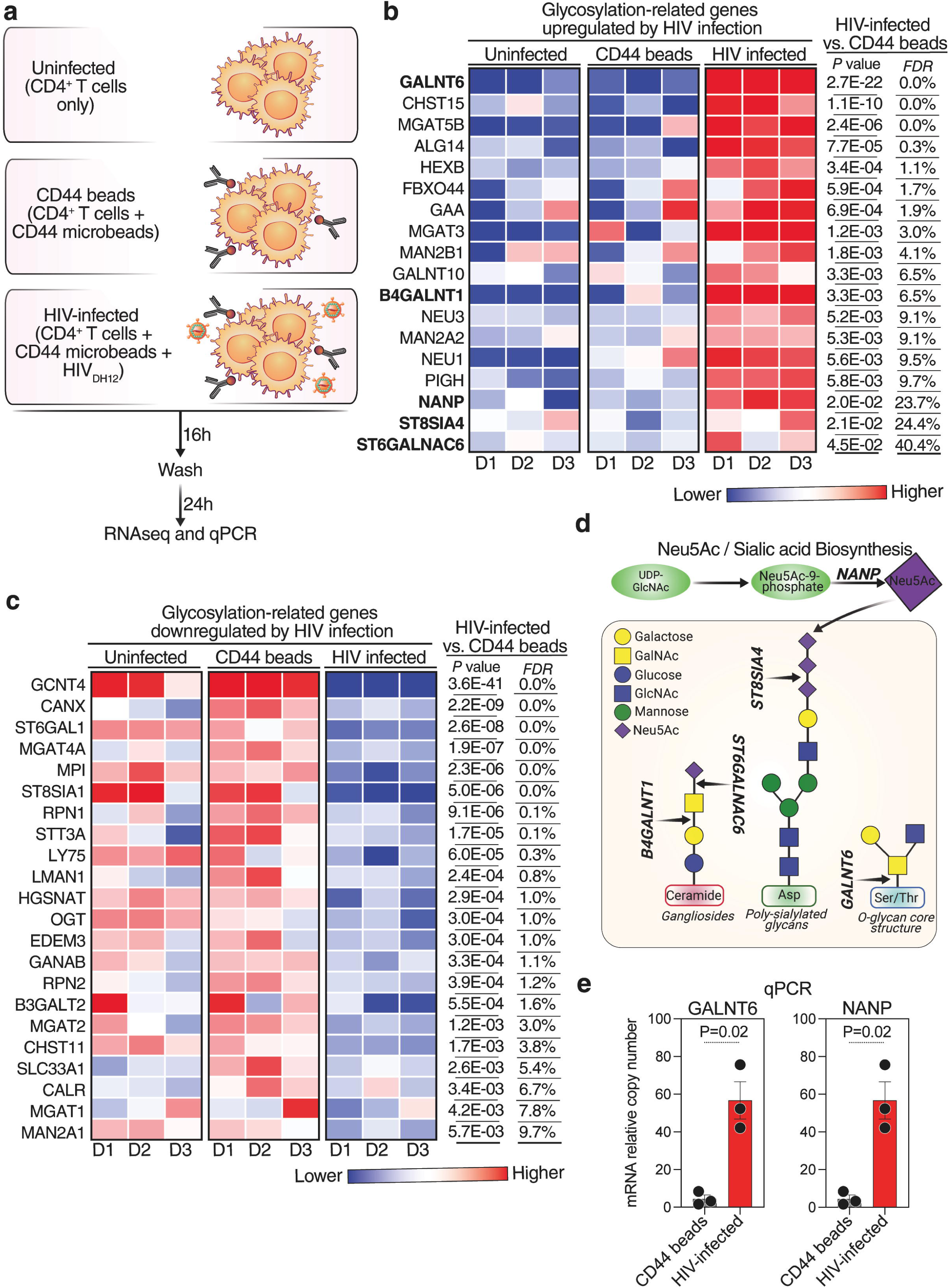
HIV infection alters the glycosylation machinery of primary CD4^+^ T cells. **(a)** Schematic representation of the workflow used to identify changes in the glycosylation machinery post-HIV infection. Non-activated primary CD4^+^ T cells from three HIV-negative donors were incubated with CD44 microbeads or a mixture of CD44 microbeads and HIVDH12 for 16 hours, washed, and cultured for another 24 hours before being processed for RNA sequencing. **(b-c)** Heatmaps showing glycosylation-related genes upregulated (b) or downregulated (c) in response to HIV infection. Red indicates higher gene expression, while blue indicates lower gene expression. Paired t-tests with p-values corrected using the Benjamini-Hochberg procedure were applied to determine the false discovery rate (FDR) and control for multiple testing. Glycosylation-related genes involved in Siglec ligand upregulation are shown in bold. **(d)** Schematic representation illustrating the role of key genes in sialic acid biosynthesis (*NANP*), core glycan biosynthesis (*GALNT6*), glycan elongation (*B4GALNT1*), and increased sialylation (sialyltransferases—*ST6GALNAC6*, *ST8SIA4*). **(e)** Bar plot representing the relative mRNA expression of GALNT6 and NANP in HIV-infected CD4^+^ T cells compared to CD44 bead-treated control cells. Error bars represent the mean ± standard error of the mean. Statistical analysis was performed using paired t-tests. N = 3.

For transcriptomic analyses, we focused on the expression of 560 genes involved in the glycosylation machinery of cells, including those involved in the addition, removal, or binding of glycans (**Supplementary Table 1**). RNA-seq analysis revealed that HIV infection significantly altered the expression of several glycosylation-related genes (**Fig. 2b-c**). Notably, several genes involved in sialic acid biosynthesis and glycan modification were upregulated, including GALNT6, NANP, B4GALNT1, ST8SIA4, and ST6GALNAC6, which are essential for the biosynthesis of sialoglycan ligands recognized by multiple Siglecs. GALNT6 initiates *O*-linked glycosylation by transferring GalNAc to serine or threonine residues. NANP regulates sialic acid precursor availability. B4GALNT1 extends glycan structures, enabling sialylation, and ST8SIA4 and ST6GALNAC6 catalyze the addition of sialic acids in α-2,8 and α-2,6 linkages, respectively, to produce functional Siglec ligands (**Fig. 2d**) ^34, 35, 36, 37^. This upregulation likely enhances specific glycosylation pathways, contributing to increased Siglec ligand expression on HIV-infected CD4^+^ T cells (**Fig. 2d**).

To validate the RNA-seq findings, we employed qPCR to examine the expression of GALNT6 and NANP, key genes involved in *O*-glycosylation initiation and sialic acid biosynthesis, respectively ^34, 35, 36, 37^. Consistent with the RNA-seq results, qPCR confirmed a significant upregulation of these genes in HIV-infected cells compared to cells treated with CD44 microbeads alone (**Fig. 2e**). These findings suggest that HIV infection reprograms the glycosylation machinery of CD4^+^ T cells, including the upregulation of pathways critical for the biosynthesis of Siglec ligands, potentially facilitating immune evasion.

### Siglec-3, -7, and -9 are highly expressed on monocytes and neutrophils, and disrupting sialic acid-Siglec interactions enhances the cytotoxic activity of these immune cells against HIV-infected cells in vitro

Given that HIV infection induces the expression of sialoglycan ligands for Siglec-3, -7, and -9, we hypothesized that these ligands facilitate immune evasion by HIV-infected cells. To test this, we developed a conjugate of sialidase (Sia; an enzyme that removes sialoglycans) and an HIV-specific broadly neutralizing antibody (bNAb; 10-1074). This conjugate, 10-1074-Sia, directs sialidase specifically to the surface of HIV-infected cells via antibody targeting, removing sialoglycan ligands and disrupting Siglec interactions. This disruption alleviates a negative immune suppression mechanism and enhances the cytotoxic capacity of immune cells expressing Siglecs against HIV-infected cells with high levels of sialoglycans (**Fig. 3a**).

**Figure 3.**
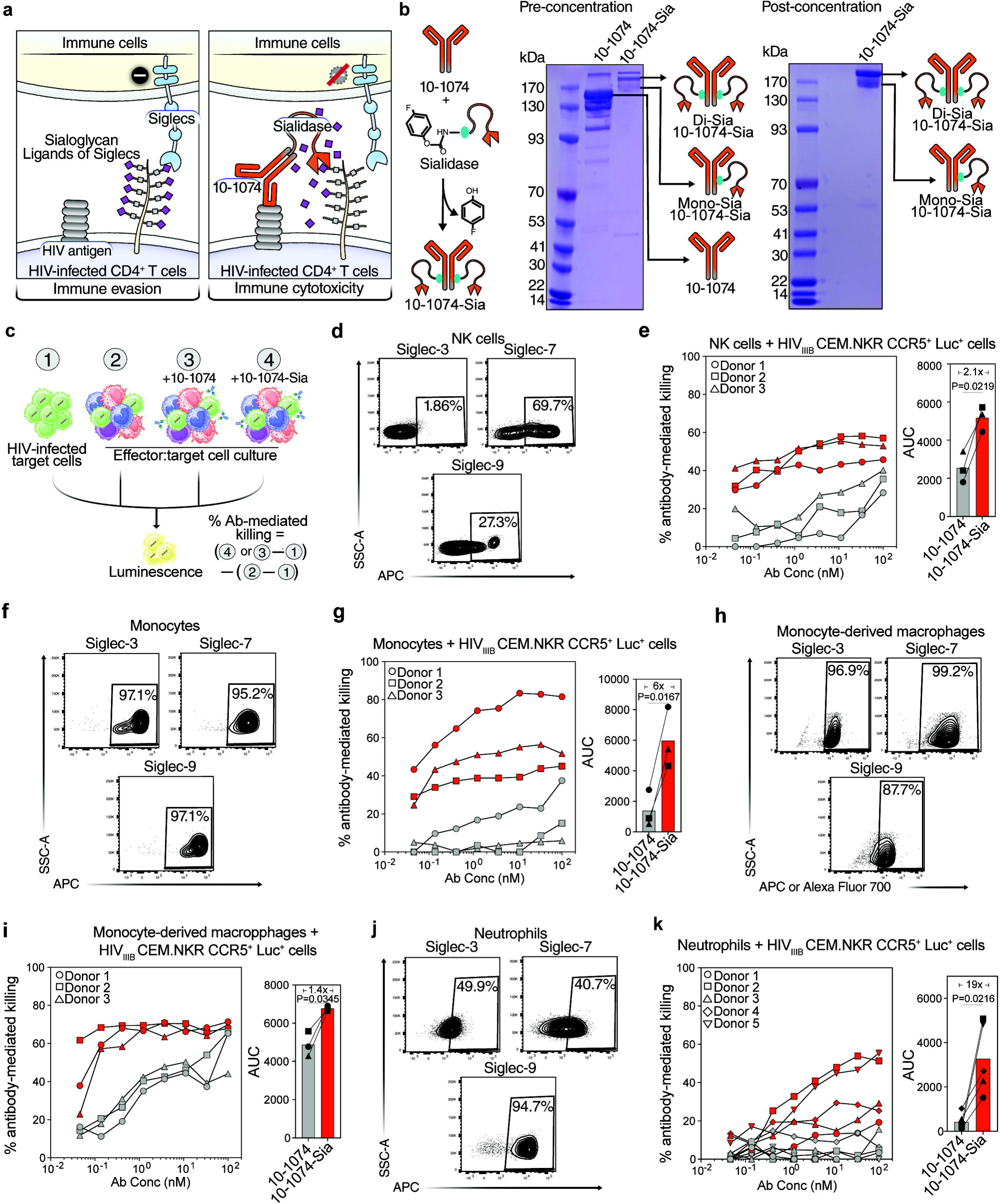
Siglec-3, -7, and -9 are highly expressed on monocytes and neutrophils, and disrupting sialic acid-Siglec interactions enhances the cytotoxic activity of these immune cells against HIV-infected cells *in vitro*. **(a)** Left panel: Increased expression of sialylated Siglec ligands on the surface of HIV-infected CD4^+^ T cells protects them from immune cell-mediated cytotoxicity. Right panel: Disruption of the sialoglycan-Siglec interaction using sialidase conjugated to 10-1074 (10-1074-Sia) enhances the immune cell-mediated clearance of HIV-infected cells. **(b)** Schematic representation of the strategy used for the conjugation of sialidase (Sia) to the HIV bNAb 10-1074, along with two SDS-PAGE gels showing the successful construction (left) and the purity (right) of the conjugate. **(c)** Workflow and equation used to evaluate antibody-mediated killing by multiple immune cell sub-populations. CEM.NKR CCR5^+^ Luc^+^ cells were infected with HIV_IIIB_ for 72 hours, followed by co-culture with whole PBMCs, NK cells, monocytes, monocyte-derived macrophages (MDMs), or neutrophils for 16 hours in the presence of increasing concentrations of 10-1074 or 10-1074-Sia. Cytotoxicity was assessed by measuring the decrease in relative luminescence units (RLU) compared to infected target cells alone. **(d, f, h, and j)** Representative plots showing the expression of Siglec-3, Siglec-7, and Siglec-9 on NK cells, monocytes, MDMs, and neutrophils, respectively. **(e, g, I, and k)** Antibody-mediated cytotoxicity of NK cells, monocytes, MDMs, and neutrophils, respectively against HIV_IIIB_-infected CEM.NKR CCR5^+^ Luc^+^ target cells was measured in the presence of increasing concentrations of 10-1074 or 10-1074-Sia by subtracting direct cytotoxicity from the total cytotoxicity (left panels), and AUCs were computed to assess the overall effect (right panels). Effector-to-target (E:T) ratios of 10:1, 10:1, 2.5:1, and 10:1 were used for NK cells, monocytes, MDMs, and neutrophils-based assays, respectively. Statistical analysis for AUC was performed using paired t-tests. N=3-5.

The 10-1074-Sia conjugate was synthesized using the proximity-induced antibody labeling (pClick) technology ^38^ (**Fig. 3b**). pClick allows site-specific conjugation of antibodies while minimizing the disruption of antigen and Fc receptor binding ^30^. To construct the conjugate, an antibody-binding protein (FB) with low risk of immunogenicity was genetically fused to the *N*-terminus of Sia via a flexible GGGS linker. Using genetic code expansion, the non-canonical amino acid 4-fluorophenyl carbamate lysine (FPheK) was introduced at position 25 of FB to enable site-specific crosslinking. FB specifically binds to the CH2-CH3 junction of the antibody 10-1074, positioning FPheK for proximity-induced covalent crosslinking with a nearby lysine residue and facilitating the formation of the 10-1074-Sia conjugate (**Fig. 3b**).

To assess the impact of disrupting Siglec-ligand interactions using 10-1074-Sia to enhance the cytotoxic capacity of immune cells against HIV-infected cells, we employed a cytotoxicity assay illustrated in **Fig. 3c**. HIV-infected CEM.NKR CCR5^+^ Luc^+^ cells expressing luciferase as a marker of infection were co-cultured with effector immune cells in the presence or absence of 10-1074 or 10-1074-Sia. Antibody-mediated cytotoxicity was calculated by measuring the difference between total cytotoxicity and direct cytotoxicity (without the antibody). 10-1074-Sia significantly enhanced the cytotoxic capacity of peripheral blood mononuclear cells (PBMCs) to eliminate HIV-infected cells compared to 10-1074 alone, as evidenced by both total killing (**Supplementary Fig. 1a**) and antibody-mediated killing (after subtracting direct killing; **Supplementary Fig. 1b**).

To determine which immune cell types are involved in Siglec-mediated immune evasion of HIV-infected cells, we first analyzed the expression of Siglec-3, -7, and -9 on six different immune cell types: CD8^+^ T cells, γδ T cells, NK cells, monocytes, macrophages, and neutrophils. Peripheral CD8^+^ T cells and γδ T cells did not express any of these Siglecs (**Supplementary Fig. 2a-b**), indicating that they are unlikely to contribute to Siglec-mediated immune evasion. NK cells expressed moderate levels of Siglec-7 and Siglec-9 but not Siglec-3 (**Fig. 3d**). Consistently, 10-1074-Sia moderately enhanced the ability of isolated NK cells to target HIV-infected cells compared to 10-1074 alone, as shown by both total killing (**Supplementary Fig. 3a**) and antibody-mediated killing (after subtracting direct killing; **Fig. 3e**). These findings are consistent with our previous data showing that Siglec interactions contribute to the ability of HIV-infected cells to evade NK immunosurveillance ^24^.

However, beyond NK cells, myeloid cells, including monocytes, monocyte-derived macrophages (MDMs), and neutrophils, expressed the highest levels of Siglec-3, -7, and -9 (**Fig. 3f-k**). Disrupting Siglec interactions using 10-1074-Sia significantly enhanced the cytotoxicity of these cells against HIV-infected cells, as indicated by both total killing (**Supplementary Fig. 3b-d**) and antibody-mediated killing (after subtracting direct killing; **Fig. 3f-k**). These effects were most pronounced in isolated monocyte cultures, with a 6-fold increase, in cytotoxicity and neutrophil cultures, with a 19-fold increase in cytotoxicity compared to 10-1074 alone (**Fig. 3g, 3k**). These findings suggest that HIV-infected cells evade immune responses from several myeloid cells by inducing specific sialoglycan ligands. Disrupting these interactions overcomes this immune evasion, enhancing the cytotoxic ability of these immune cells against HIV-infected cells.

### 10-1074-Sia enhances the cytotoxicity of myeloid cells selectively against HIV-infected cells

The 10-1074-Sia conjugate used in the cytotoxic assays in Fig. 3 is designed to direct sialidase to the cell surface of HIV-infected cells expressing viral antigens, enabling the removal of sialic acid from these cells. This disrupts interaction between the sialoglycans on infected cells and Siglecs on immune cells, enhancing the susceptibility of the infected cells to immune-mediated clearance without affecting HIV-uninfected cells. To test this, we co-cultured uninfected CEM.NKR CCR5^+^ Luc^+^ cells (stained with Incucyte rapid red dye) and HIV_IIIB_-infected CEM.NKR CCR5^+^ Luc^+^ cells (stained with Incucyte rapid green dye) to allow for visual characterization of both cell types using live imaging. This cell mixture was then co-cultured with primary monocytes isolated from four HIV-negative donors in the presence or absence of an isotype control, 10-1074, or 10-1074-Sia. Live cell imaging data, shown in **Fig. 4a-b**, along with cumulative data from the four donors (**Fig. 4c**), show that the 10-1074-Sia conjugate enhances the ability of monocytes to eliminate HIV-infected cells while minimally impacting HIV-uninfected cells. These findings indicate that bNAb-sialidase conjugates can selectively induce monocyte-mediated cytotoxicity against HIV-infected cells, offering an advantage over the bNAb alone.

**Figure 4.**
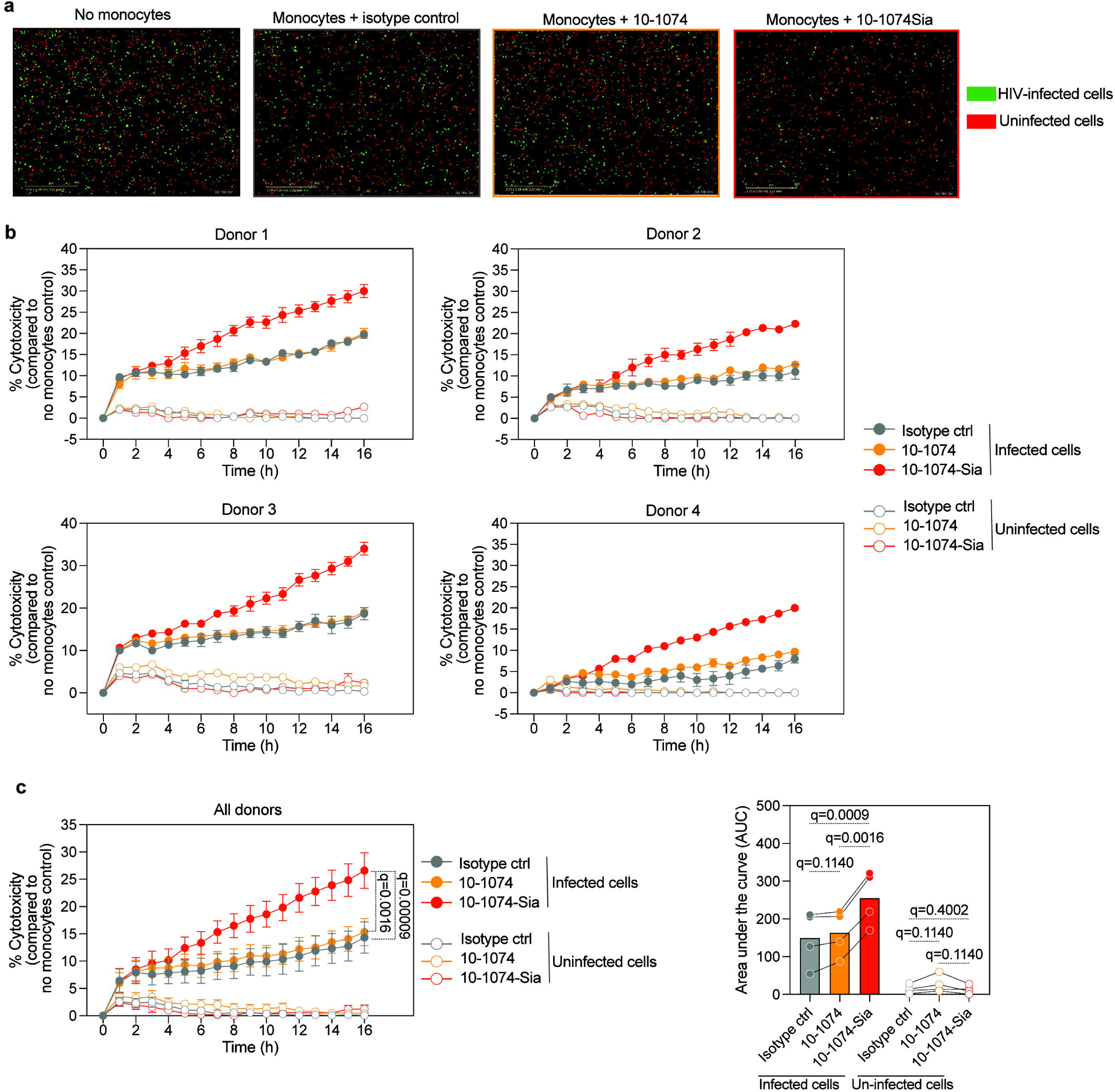
10-1074-Sia exhibits selective cytotoxicity against HIV-infected CD4^+^ T cells. **(a)** Primary monocytes from healthy donors were co-cultured with uninfected CEM.NKR CCR5^+^ Luc^+^ (stained with Incucyte Rapid red dye) and HIV_IIIB_-infected CEM.NKR CCR5^+^ Luc^+^ (stained with Incucyte Rapid green dye) target cells in the presence of isotype control, 10-1074, or 10-1074-Sia, and imaged for 16 hours using the Incucyte SX1. Representative images from each condition across four independent experiments are shown. **(b)** Kinetic measurement of monocyte-mediated cytotoxicity against HIV-infected cells (closed circles) and uninfected cells (open circles) was performed at 1-hour intervals over a 16-hour period. Each panel represents data obtained in triplicates from one donor (E:T ratio = 5:1; n = 4 donors). Percent cytotoxicity was calculated as the change in the green or red cell count under antibody-treated conditions compared to the target cells-only control. **(c)** Kinetic data representing cytotoxicity from four donors (left panel). The AUC was calculated to demonstrate cytotoxic potential over time (right panel). Statistical analysis was performed using paired one-way ANOVA, corrected by the two-stage step-up procedure of Benjamini, Krieger, and Yekutieli. N = 4.

### Disrupting Siglec interactions enhances the capacity of monocytes and neutrophils to target autologous HIV-infected primary CD4^+^ T cells and increases monocyte immune activation and cytotoxic capacity

Our data, thus far indicate that disrupting Siglec interactions significantly enhances monocyte- and neutrophil-mediated cytotoxicity against HIV-infected cells. However, as the cytotoxic assays shown in Figs 3-4 used an HIV-infected cell line as target cells, we next examined whether these effects hold true in autologous assays using primary monocytes and neutrophils from the same donors as primary HIV-infected CD4^+^ T cells. To address this, we isolated monocytes or neutrophils from seven HIV-negative donors and co-cultured them with HIV_TYBE_-infected CD4^+^ T cells from the same donors in the presence or absence of an isotype control, 10-1074, or 10-1074-Sia (**Fig. 5a; Supplementary Figure 4**). As shown in **Fig. 5b-e**, disrupting Siglec interactions using 10-1074-Sia significantly enhanced the ability of autologous monocytes and neutrophils to reduce intracellular HIV p24 levels in the co-cultures.

**Figure 5.**
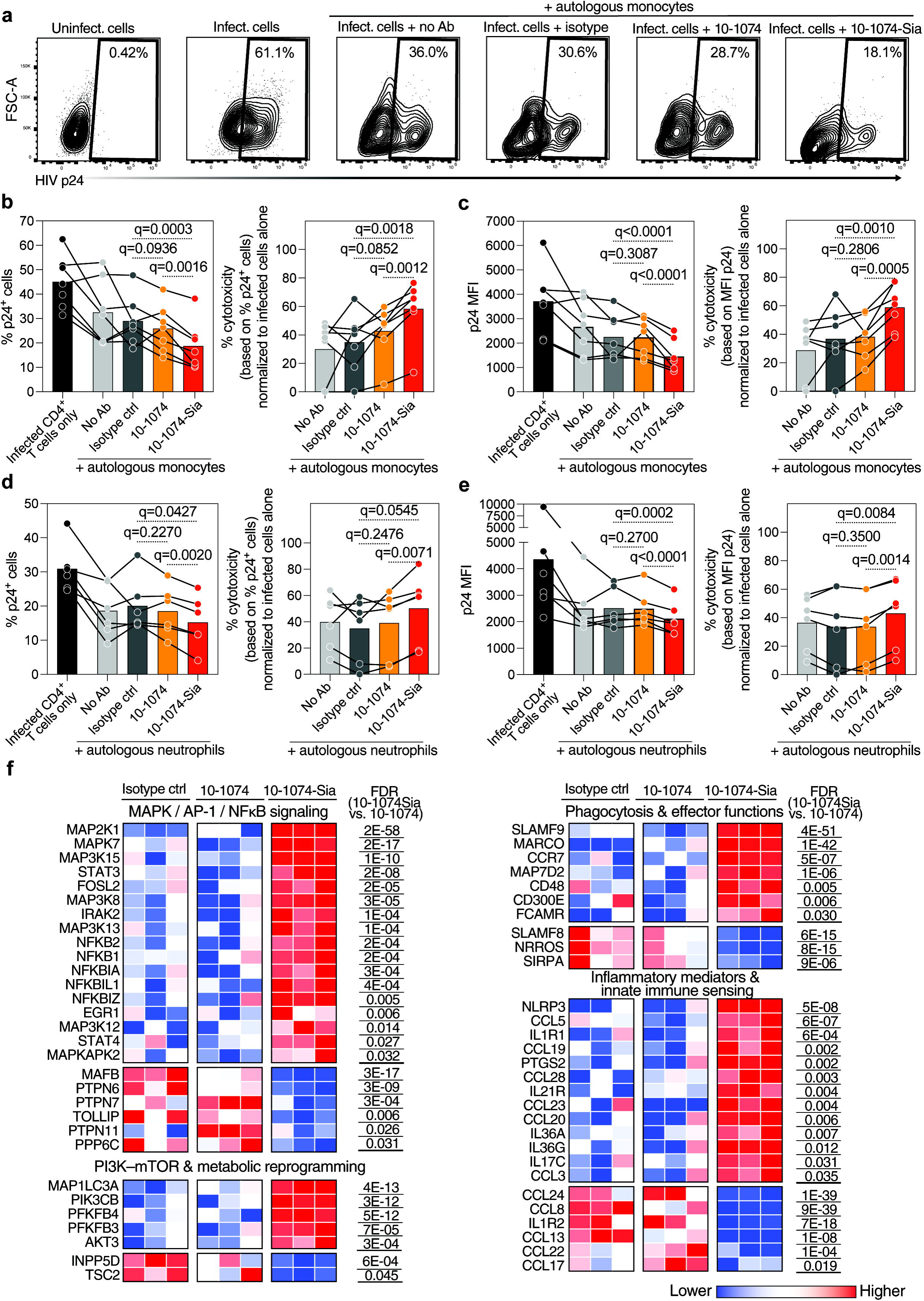
Disrupting Siglec interactions enhances the ability of monocytes and neutrophils to target autologous HIV-infected primary CD4^+^ T cells and increases monocyte immune activation and cytotoxic capacity. Activated primary CD4^+^ T cells were infected with HIV_TYBE_ for 72 hours, followed by co-culture with autologous monocytes in the presence of isotype controls, 10-1074, or 10-1074-Sia for 16 hours (E:T ratio = 10:1). After overnight incubation, the co-cultures were stained with Zombie Aqua (BV510) and antibodies against CD3 and intracellular HIV p24. **(a)** Representative contour plots showing the decline in p24+ HIV-infected CD4^+^ T cells when co-cultured with autologous effector cells in the presence of 10-1074 or 10-1074-Sia. **(b-c)** The left panels show (b) the percentage of p24+ HIV-infected CD4^+^ T cells and (c) p24 median fluorescence intensity (MFI). The right panels depict normalized cytotoxicity based on either the percentage of p24+ cells (b) or reduction in p24 MFI (c). **(d-e)** HIV_TYBE_-infected primary CD4^+^ T cells were used as targets for autologous neutrophils, and cytotoxicity was evaluated by intracellular p24 staining 16 hours after co-culture (E:T ratio = 5:1). The left panels show the percentage of p24+ cells (d) and p24 MFI (e), while the right panels depict cytotoxicity based on either percentage p24+ cells (d) or p24 MFI (e). Statistical analysis was performed using paired one-way ANOVA, corrected by the two-stage step-up procedure of Benjamini, Krieger, and Yekutieli. **(f)** Transcriptomic analysis was performed on monocytes co-cultured with autologous HIV-infected CD4^+^ T cells for 16 hours in the presence of isotype control, 10-1074, or 10-1074-Sia. Differential gene expression analysis was conducted using DESeq2. Heatmaps represent genes differentially regulated in MAPK/AP-1/NF-κB signaling (top left), PI3K-mTOR and metabolic reprogramming (bottom left), phagocytosis and effector functions (top right), and inflammatory mediators and innate immune sensing (bottom right). Red indicates higher gene expression, while blue indicates reduced gene expression. Paired t-tests with p-values corrected using the Benjamini-Hochberg procedure were applied to determine the false discovery rate (FDR) and control for multiple testing. N = 3–7.

To examine the immune signaling pathways involved in this enhanced cytotoxicity, we performed a transcriptomic analysis (using RNA-sequencing) on monocytes co-cultured with autologous CD4^+^ T cells in the presence or absence of an isotype control, 10-1074, or 10-1074-Sia. Transcriptomic analysis of these co-cultures reveals that 10-1074-Sia treatment significantly modulated multiple genes and pathways associated with enhanced monocyte activation and cytotoxicity (FDR < 0.05), compared to both 10-1074 and isotype treated cultures (**Fig 5f**). In particular, and as expected from blocking Siglec interactions, we observed robust upregulation of NF-κB components (NFKB1, NFKB2, NFKBIA, NFKBIZ, NFKBIL1) and several MAPK kinases (e.g., MAP3K8, MAP3K12, MAP3K13, MAP3K15, MAP2K1, MAPK7, MAPKAPK2), as well as AP-1-associated transcription factors (FOSL2, EGR1). Together, these changes indicate heightened transcriptional output of proinflammatory genes, consistent with classic NF-κB- and AP-1-driven cytokine production. Meanwhile, key negative regulators normally engaged by Siglec–SHP pathways, such as PTPN6 (SHP1), PTPN11 (SHP2), TOLLIP, and PPP6C, were significantly downregulated, removing multiple “brakes” on NF-κB/MAPK signaling (**Fig. 5f, top left**). We also observed a significant activation of the PI3K-AKT-mTOR axis in 10-1074-Sia treated cultures than controls **(Fig. 5f, bottom left)**. Specifically, PIK3CB and AKT3 were upregulated, promoting survival and metabolic reprogramming, as were PFKFB3 and PFKFB4, key enzymes driving glycolysis in activated myeloid cells. Simultaneously, downregulation of TSC2 lifts checkpoints on PI3K–AKT–mTOR activity, reinforcing a proinflammatory metabolic state.

In addition, our data show a significant upregulation of surface receptors and mediators that facilitate phagocytosis and effector functions of myeloid cells (**Fig. 5f, top right**). Genes such as MARCO, CD300E, SLAMF9, FCAMR, and CCR7 were significantly elevated in 10-1074-Sia cultures (than controls), indicating improved pathogen recognition and cell–cell interaction. Consistently, downregulation of SIRPA and SLAMF8 removes inhibitory signals on phagocytic activity, while decreased NRROS supports a more potent oxidative burst. Together, these changes suggest a shift toward heightened phagocytosis and cytotoxic effector mechanisms. Finally, we observed a strong reconfiguration of inflammatory mediators and innate immune sensing pathways by 10-1074-Sia treatment, highlighted by the upregulation of several proinflammatory chemokines (CCL19, CCL20, CCL3, CCL5, CCL23, CCL28) and cytokines (IL36A, IL36G, IL17C), along with NLRP3 (inflammasome sensor) and PTGS2 (COX-2). In contrast, inhibitory chemokines (CCL17, CCL22, CCL13, CCL24, CCL8) and the IL-1 decoy receptor (IL1R2) were downregulated, shifting the balance towards a more proinflammatory milieu (**Fig. 5f, bottom right**). Together, these changes support that Siglec blockade drives a reprogramming of monocytes, reducing inhibitory checkpoints while upregulating proinflammatory, metabolic, and phagocytic pathways. This coordinated response ultimately enhances monocyte activation and cytotoxic capacity, consistent with our functional cytotoxicity assays.

### Targeting sialic acid/Siglec interactions enhances the anti-HIV activity of 10-1074 in a humanized mouse model of HIV infection

Our data thus far suggest that HIV infection induces the overexpression of sialoglycan ligands for several Siglecs, primarily expressed on multiple myeloid cells and NK cells. These sialoglycans bind to Siglecs on immune cells, triggering inhibitory signals that contribute to the ability of HIV-infected cells to evade immune surveillance. We also show that disrupting these interactions using a conjugate of 10-1074 and sialidase can selectively target HIV-infected cells, enhancing the cytotoxic capacity of these immune cells against HIV-infected cells with minimal impact on HIV-uninfected cells, as shown by *in vitro* assays using both cell lines and autologous primary cells.

Next, we tested this strategy *in vivo* using a humanized mouse model of HIV infection ^39^. Five-week-old immunodeficient NSG mice (n=15) were humanized by engrafting isolated memory CD4^+^ T cells from healthy donors (**Supplementary Fig. 5**; **Fig. 6a**). Thirty-four days after humanization, mice were randomized into three groups based on the number of human CD4^+^ T cells per µl of blood (**Supplementary Fig. 6;** **Fig. 6b**). Mice were then infected with HIV-1_SUMA_ (a transmitted/founder virus), and one week later, the three groups had similar viral loads (**Fig. 6c**). Mice were subsequently treated weekly for five weeks with one of the following: (1) PBS (control), (2) 10-1074 combined with autologous monocytes and NK cells (**Supplementary Fig. 7**), or (3) 10-1074-Sia combined with autologous monocytes and NK cells.

**Figure 6.**
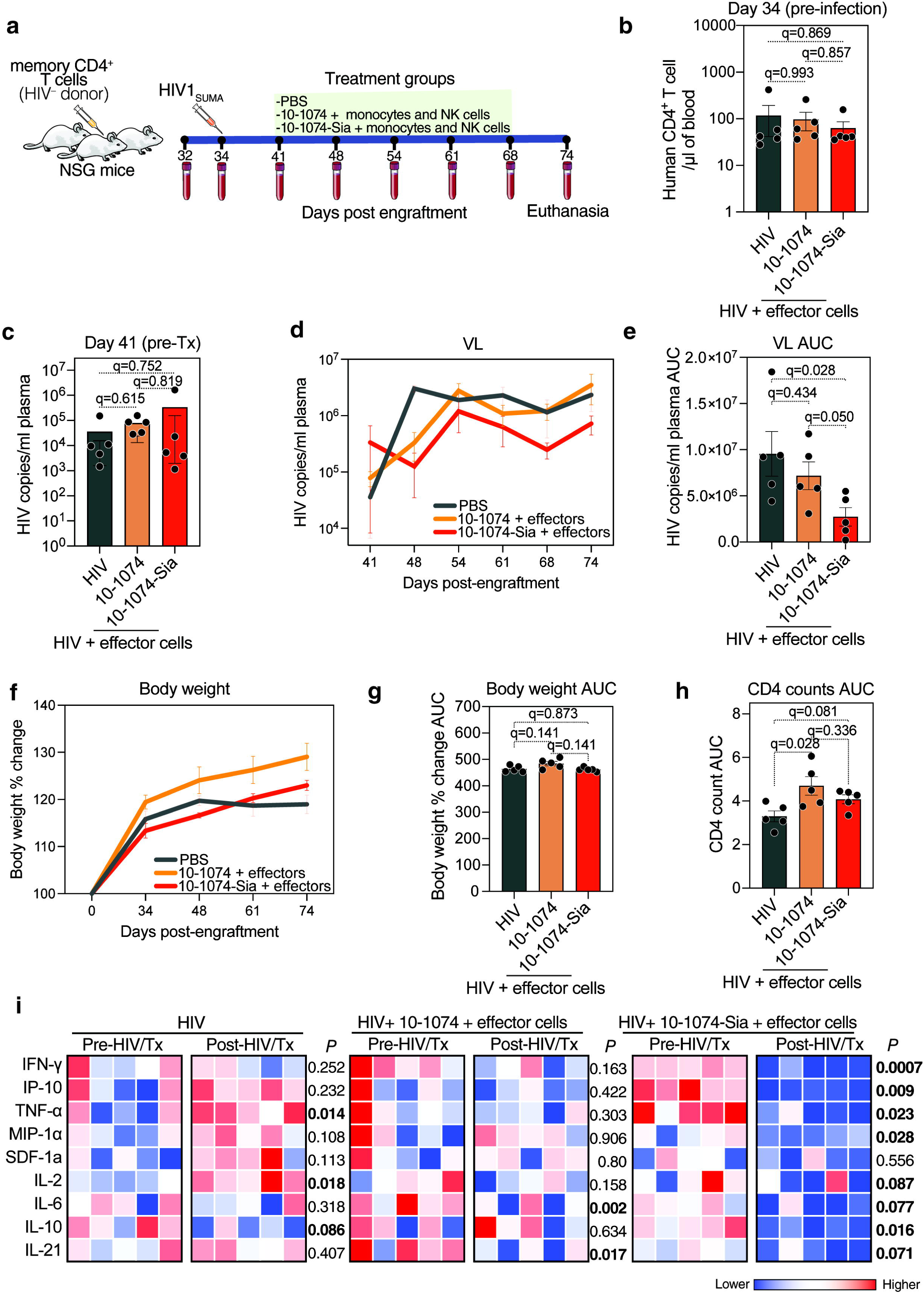
Targeting sialic acid/Siglec interactions enhances the anti-HIV activity of 10-1074 in a humanized mouse model of HIV infection. **(a)** Study design outlining the generation of humanized NSG mice engrafted with human memory CD4^+^ T cells, followed by HIV_SUMA_ infection and weekly treatment with 10-1074 or 10-1074-Sia (0.75 mg/dose). Monocytes and NK cells were injected weekly. Control mice were treated with PBS only (n = 5 per group). **(b)** Peripheral blood human CD4^+^ T cell counts before HIV_SUMA_ infection, measured by flow cytometry. **(c)** Humanized NSG mice were infected intravenously with 1500 TCID_50_ of HIV_SUMA_. Establishment of infection was confirmed by measuring HIV viral load (VL) prior to treatment initiation. **(d-e)** Weekly plasma viral load, measured as HIV copies/mL (y-axis) over the experimental duration (x-axis) (d), and presented as AUC (e) to represent cumulative HIV infection over time across the three groups. **(f-g)** NSG mice were weighed weekly, starting before engraftment and continuing until the end of the study. Data are presented as percentage changes from the initial weight. **(h)** Absolute CD4^+^ T cell counts in peripheral blood, presented as AUC to represent cumulative CD4^+^ T cell levels over time. Statistical analyses for panels (b), (c), (e), and (g) were performed using the Kruskal-Wallis test, corrected by the two-stage step-up procedure of Benjamini, Krieger, and Yekutieli. **(i)** Heatmap representing the protein levels of several pro-inflammatory markers in plasma across the three treatment groups, before (day 34) and after HIV infection and treatment initiation (day 48). Red indicates higher marker abundance, while blue indicates lower abundance. P-values were computed using paired t-tests. N = 15.

While treatment with 10-1074 plus effector cells (autologous monocytes and NK cells) alone did not significantly reduce viral load compared to PBS-treated mice, 10-1074-Sia treatment with autologous effector cells demonstrated superior efficacy in reducing viral load (**Fig. 6d-e**). Importantly, mice treated with 10-1074-Sia did not experience significant weight loss compared to PBS-treated mice (**Fig. 6f-g**), suggesting minimal off-target effects of the sialidase conjugation. CD4^+^ T cell depletion was reduced in mice treated with either 10-1074 or 10-1074-Sia compared to controls (**Fig. 6h**). Lastly, while 10-1074 + effector cells were unable to reduce HIV-associated inflammation, 10-1074-Sia + effector cells significantly reduced HIV-induced inflammatory markers, including TNF-α, IFN-γ, IP10, and MIP-1α (**Fig. 6i**). Together, these data highlight 10-1074-Sia as a promising anti-HIV immunotherapeutic strategy.

## DISCUSSION

This study identifies a novel immune evasion mechanism employed by HIV, wherein the virus reprograms the glycosylation machinery of infected CD4^+^ T cells to upregulate sialoglycans that specifically engage multiple, yet distinct, Siglecs expressed on immune cells, particularly myeloid cells. These interactions suppress the cytotoxic functions of monocytes, neutrophils, and other immune cells, thereby facilitating the immune evasion of HIV-infected cells. By targeting this glyco-immune checkpoint using a sialidase-conjugated bNAb, 10-1074-Sia, we show enhanced immune clearance of HIV-infected cells both *in vitro* and *in vivo*, providing a promising therapeutic strategy.

The cell-surface glycome plays pivotal roles in modulating various cellular processes^40^ and mediating cell-cell^41, 42^ and cell-pathogen^43^ interactions. For instance, the cell-surface glycosylation of T cells has recently been implicated in influencing HIV susceptibility and selection during transmission ^44^. While aberrant glycosylation is a well-established mechanism employed by cancer cells for immune evasion, whether HIV leverages a similar strategy has remained unclear. Our findings reveal that, beyond influencing cell susceptibility and viral selection, HIV-induced glycomic alterations on infected cells actively contribute to immune evasion by promoting binding to inhibitory Siglecs on immune cells. This highlights the multifaceted role of cell-surface glycosylation in the pathogenesis and persistence of HIV. Whether this glyco-immune evasion mechanism contributes to the persistence of HIV-infected cells during suppressive antiretroviral therapy (ART) warrants further exploration. Recent studies indicating that elevated Siglec expression correlates with shorter time to viral rebound after treatment interruption^45^ further highlight the potential of Siglec interactions for modulating anti-HIV immune responses, particularly those mediated by innate immune cells.

Myeloid cells, including monocytes, macrophages, and neutrophils, play critical roles in controlling HIV infection through their immune cytotoxicity and effector functions. These cells can recognize and eliminate infected cells via mechanisms such as antibody-dependent cellular phagocytosis (ADCP), contributing significantly to the control of viral replication, particularly in tissues ^46, 47, 48, 49^. However, the ability of HIV-infected cells to evade these immune-mediated mechanisms by inducing the expression of inhibitory glycans could present a significant barrier to effective immune clearance. Our study focuses on the impact of Siglec interactions on the direct cytotoxicity of innate immune cells. However, Siglec interactions on myeloid cells could have broader implications, such as inhibiting antigen-presenting cell functions and indirectly suppressing downstream T cell responses ^50, 51, 52^. Therefore, the HIV-mediated induction of Siglec interactions, driven by elevated Siglec ligands, may also impact other critical myeloid cell functions, such as their role in shaping adaptive immune responses during HIV infection. Investigating these mechanisms is needed and could be crucial for improving the efficacy of HIV eradication strategies that rely on enhancing anti-HIV immune functions, including antibody-mediated effector functions, such as those enabled by bNAbs.

In this study, we show that disrupting Siglec-ligand interactions using 10-1074-Sia significantly enhances the cytotoxic activity of myeloid cells against HIV-infected cells, with minimal impact on uninfected cells. Blocking the Siglec – sialoglycan interaction, induced multifaceted signaling changes in the monocytic effector cells promoting both phagocytic and pro-inflammatory responses against HIV-infected cells. Similar approaches have been tested safely and effectively to augment innate immune functions against cancer cells ^29, 30, 53^. This specificity and efficacy underscore the therapeutic potential of this approach as a targeted strategy to overcome immune evasion. Examining the effects of this approach on innate immune responses, particularly those mediated by NK cells and myeloid cells, during suppressive ART and after ART interruption, will be crucial. Importantly, this strategy relies on the expression of viral antigens on the surface of HIV-infected cells, which positions it as a complementary approach to existing HIV eradication strategies, such as “shock and kill” ^54^. By reactivating latent HIV reservoirs, “shock and kill” renders infected cells susceptible to immune clearance, which could be further amplified by sialidase-conjugated antibodies that neutralize Siglec-mediated immune suppression. Future studies should investigate the potential of combining multiple bNAbs conjugated with sialidase to target diverse epitopes of viral antigens on the surface of HIV-infected cells. Such combinations could broaden and enhance the efficacy of this strategy, ensuring robust immune clearance. Together, these studies could position this glycan-focused strategy as a novel and integral component of combination therapies aimed at eradicating HIV-infected cells.

While our findings provide novel evidence for the role of sialoglycan-Siglec interactions in HIV immune evasion, there are several limitations to consider. First, future studies should examine the glycosylation profiles of infected cells isolated from the blood and tissues of people living with HIV (PLWH) on ART. However, isolating these cells remains challenging, as there are currently no specific cell-surface markers to distinguish HIV-infected cells from uninfected cells. Despite this, our recent studies have shown that transcriptionally active HIV-infected cells isolated from the blood of PLWH on ART exhibit an aberrant glycosylation profile, including elevated levels of sialic acid ^55^. This suggests that glycomic alterations, potentially including enrichment of Siglec ligands, are evident *in vivo* in PLWH on ART. Second, immune interactions differ between blood and tissues. For instance, while peripheral T cells do not express Siglecs, Siglec expression is tissue-specific and tumor-infiltrating lymphocytes (TILs) express Siglec-9 ^21, 27^. Siglec-9^+^ TILs are tumor-specific, and inhibiting Siglec-9 signaling significantly increases T cell-mediated tumor cell killing ^21, 27^. Whether HIV-induced sialoglycans similarly modulate differential immune responses in tissues warrants further investigation. Third, this study primarily focused on Siglecs with documented roles in immune evasion during cancer and other pathological conditions—specifically, Siglecs-3, -7, and -9. However, the Siglec family comprises 15 members with differential roles in modulating immunity. Examining the roles of other Siglecs could provide further insights into their contribution to immune responses against HIV-infected cells. Lastly, as mentioned, it will be critical to assess the efficacy of the sialidase-conjugated antibody approach in enhancing immune-mediated clearance of HIV-infected cells in animal models of HIV infection treated with ART. Studies in models with intact immune systems will be particularly important, as they would allow for a more comprehensive examination of Siglec-mediated interactions across multiple facets of anti-HIV immunity. Despite these limitations, our study significantly advances the understanding and targeting of glyco-immune checkpoints in HIV pathogenesis.

## MATERIALS AND METHODS

### Ethics Statement

Blood and cryopreserved PBMCs from HIV-negative donors were collected from the Wistar Institute and Human Immunology Core, University of Pennsylvania, respectively. Research protocols were approved by The Wistar Institute committee on Human Research (IRB# 2110176-6a). Written informed consent was obtained, and all data and specimens were coded to protect confidentiality. All human experimentation was conducted in accordance with the guidelines of the US Department of Health and Human Services and those of the authors’ institutions.

### Isolation of primary CD4 T^+^ cells and effector cells

Primary human CD4^+^ T cells, CD8^+^ T cells, γδ T cells, monocytes and NK cells were isolated from fresh or cryopreserved PBMCs from healthy donors by immunomagnetic negative selection using the EasySep™ Human CD4^+^ T Cell Isolation Kit (Catalog #17952), EasySep™ Human CD8+ T Cell Isolation Kit (Catalog #17953), EasySep™ Human Gamma/Delta T Cell Isolation Kit (Catalog #19255), EasySep™ Human Monocyte Isolation Kit (Catalog #19359) and the EasySep™ Human NK Cell Isolation Kit (Catalog #17955; STEMCELL Technologies), respectively, following manufacturer’s protocol. Neutrophils were isolated directly from whole blood using the EasySep™ Direct Human Neutrophil Isolation Kit (STEMCELL Technologies, Catalog #19666) following manufacturer’s protocol.

### Cell Culture

CEM.NKR CCR5^+^ Luc^+^ cells obtained from NIH AIDS Reagent Program (Catalog #ARP-5198, contributed by Dr. John Moore and Dr. Catherine Spenlehauer) were cultured in RPMI 1640 supplemented with 10% heat-inactivated fetal bovine serum (FBS), penicillin (50 U/ml), and streptomycin (50 mg/ml) (complete R10) at 37°C ^56^. Similarly, PBMCs, primary NK cells and monocytes were cultured in complete R10. Primary CD4^+^ T cells were cultured using the same medium supplemented with 30 U/ml recombinant human IL-2 (PeproTech, Catalog #200-02-50UG). Primary monocytes derived macrophages (MDMs) were differentiated for 7-8 days, as described previously ^31, 57^. Briefly, freshly isolated monocytes were cultured in IMDM (Gibco) supplemented with 10% AB human serum (Sigma) and 50 ng/mL M-CSF (Peprotech). After 3-4 days of differentiation, the cells were supplemented with 50 ng/mL TGF-β (Peprotech) and 50 ng/mL human IL-10 (Peprotech), and further cultured till day 8, for flow cytometry staining and cytotoxicity assays.

### Primary CD4 T cell infection with HIV_TYBE_ for siglec ligand staining

CD4^+^ T cells (2 ×10^5^) were concurrently exposed to concentrated HIV-1 TYBE ∼ 4 µg/ml (Penn Center for AIDS Research Virus and Molecular Core, University of Pennsylvania, Philadelphia) and Dynabeads Human T-Activator CD3/CD28 (Thermo Scientific, Catalog #11132D) and incubated at 37°C for 72 h hours in 48-well plates. Activation beads were added at a ratio of 5 beads/cell. *O*-glycan biosynthesis inhibitor Benzyl 2-acetamido-2-deoxy-α-D-galactopyranoside, BADG (Sigma, Catalog #B4894) was dissolved in DMSO and used at a final concentration of 2.5 mM, at time of virus exposure to block *O*-glycosylation. After 72 h, cells were separated from the beads, washed with 1X PBS supplemented with 0.5% BSA and 0.1% sodium azide (FACS buffer) and processed for intracellular p24 staining and cell surface siglec ligand staining.

### Primary CD4 T cell infection with HIV_DH12_ without T cell activation

1 mL of CEMx174-grown HIV-1 DH12 (312,500 TCID_50_) was mixed with 50 μl of HIV Infectivity Enhancement Reagent (Miltenyi Biotec, Catalog # 130-095-093) and incubated at 4°C for 30 min. 250 μl of the virus-CD44 microbead mixture was then added to 300,000 unstimulated CD4^+^ T cells in 250 μl IL-2 supplemented complete R10 media for 16 h. After overnight incubation, cells were washed with 5x volume of the R10 medium and incubated for additional 24 h and 48 h at 37°C for RNAseq and recombinant Siglec-7 Fc staining, respectively. Aliquots of virus exposed cells were taken from culture and washed with FACS buffer. HIV infection and CD4^+^ T cell activation status was evaluated by staining for intracellular p24 using KC57-RD1, anti-p24 antibody (Beckman Coulter) and anti-CD69 FITC (Biolegend) markers respectively.

### Detection of cells surface siglec ligands by flow cytometry

1 × 10^5^ uninfected or HIV-infected CD4^+^ T cells were resuspended in 100 µl FACS buffer. Indicated amounts of recombinant human Siglec-3, -7, -9 and -10 Fc proteins (R&D Systems) resuspended in PBS were added to the cells and incubated for 1 h at room temperature. Following incubation, the cells were washed twice with FACS buffer and incubated with BV421 Fc-specific rat anti-human IgG (BioLegend, Clone - M1310G05); at 1:20 dilution, for 20 min at room temperature. Cells were further washed twice, fixed for 15 min at room temperature (Biolegend Cytofix/Cytoperm), and acquired by flow cytometry on BD LSR18. Flow cytometry data were analyzed using FlowJo software.

### Transcriptomic analysis of glycosylation related genes

We performed transcriptomic analysis to determine the alterations in glycosylation-related genes in HIV-infected primary CD4^+^ T cells. RNA extraction was performed using the Qiagen RNeasy mini kit (Catalog # 74104) according to the manufacturer’s protocol. 6-12 ng of total RNA was used as input to prepare libraries using the Stranded Total RNAseq with Ribo-zero Plus kit (Illumina, San Digo,CA) as per manufacturer’s instructions. Library size was assessed using the 4200 Tapestation and the High-Sensitivity DNA assay (Agilent, Santa Clara, CA). Concentration was determined using the Qubit Fluorometer 2.0 (Thermofisher, Waltham, MA). Next Generation Sequencing with a paired-end 2×150bp run length was done on the NovaSeq X series platform (Illumina, San Diego, CA). A minimum of 30M reads per sample were acquired for each sample. RNA-seq data were aligned to GRCh37 using the RSEM (RNA-Seq by Expectation-Maximization) V1.3.3 software in conjunction with bowtie2 ^58, 59^. Raw counts below 10 were filtered out and differential gene expression analysis was performed using the DESeq2 ^60^. DESeq2 normalized count values were used to determine the log2 fold change, and adjusted p-values. P values were adjusted using Benjamini-Hochberg procedure to determine the False Discovery Rate (FDR) and control for multiple testing.

### cDNA synthesis and quantitative real-time PCR (qPCR)

15 μL of RNA was used to prepare cDNA using the SuperScript VILO cDNA Synthesis Kit (Thermo, Catalog #11754050) according to manufacturer’s protocol. Briefly, the reaction was carried out at 25°C for 10 min, 42°C for 60 min, 85°C for 5 min, and 4°C, hold. qPCR for GALNT6 and NANP was performed on QuantStudio 6 Pro (Thermo Fisher) and the following FAM-MGB probes were obtained from Thermo: GALNT6 (Hs00926629_m1), NANP (Hs00600979_m1). 18S (Thermo, Catalog #4352930E) was used as the endogenous control to calculate relative copy number.

### Detection of cell surface Siglec-3, -7 and -9 receptors

Primary γδ T cells, CD8^+^ T cells, NK cells, monocytes, macrophages and neutrophils were stained with anti-Siglec-3 (Biolegend; Clone P67.6), Siglec-7 (Biolegend; Clone 6-434) and Siglec-9 (Biolegend; Clone K8) antibodies to evaluate their cell surface expression by flow cytometry.

### Generation of 10-1074-Sia conjugate

To construct pET151-FB-E25TAG-GGGS-STSia plasmid, FB-E25TAG gene was generated by PCR using primers CY060/JP001 and template pET-22b-T5-FB-E25TAG, and inserted into pET151-STSia vector amplified with primers CY061/JP002 (**Supplementary Tables 2, 3**). The pET151-FBE25-GGGS-STSia and pUltra-KcrRS were then co-transformed into E. coli BL21(DE3) strain. Cells were grown in LB media, supplemented with ampicillin (50 μg/mL) and spectinomycin (25 μg/mL) at 37°C. When the OD reached to 0.6, 1 mM IPTG and 10 mM FPheK were added to the culture, and the culture was grown over night at 30°C. The cells were harvested by centrifugation at 5000 rpm for 10 min. The cell pellet was collected, resuspended in PBS and lysed by sonication. The supernatant was filtered, and the Sia-FB protein was purified by running through a Ni-NTA column using the FPLC (buffer A: 50 mM Imidazole in PBS, pH 7.8; buffer B: 500 mM Imidazole in PBS, pH 7.8). The eluted Sia-FB protein was concentrated and buffer exchanged into PBS using a 30K concentrator and characterized by ESI-MS analysis. Endotoxin was removed using PierceTM High-Capacity Endotoxin Removal Spin Column, 0.50 ml. 10-1074 antibody was then co-incubated with 10 equivalent amounts of Sia-FB at 37°C for 48 hours. After showing the success of the conjugation by SDS-PAGE analysis, the conjugate was concentrated using a 50K cutoff concentrator and further purified by running through a size-exclusion column using the FPLC. The purified conjugate was collected from the 96-well plate and characterized by SDS-PAGE analysis.

### Immune cell mediated killing of HIV-infected CEM.NKR CCR5^+^ Luc^+^ cells

CEM.NKR CCR5^+^ Luc^+^ cells were infected with HIV_IIIB_ (1 × 10^6^ TCID_50_/ml) on RetroNectin precoated dishes as described previously ^24^. After 72 h, the infected cells were washed, resuspended in complete R10 and plated at 2 × 10^4^ cells/well in a V-bottom 96-well microplate. HIV-infected CEM.NKR CCR5^+^ Luc^+^ were treated with indicated concentrations of 10-1074 or 10-1074-Sia conjugates for 2 h at 37°C. Following incubation, human PBMCs were added to the wells at an effector-to-target (E: T) ratio of 100:1. Similarly, cytotoxicity assays against HIV_IIIB_-infected CEM.NKR CCR5^+^ Luc^+^ cells using NK cells, monocytes or freshly isolated neutrophils were performed at an E: T ratio of 10 :1 in the presence or absence of 10-1074 or 10-1074-Sia. Monocyte-derived macrophages differentiated for 8 days were co-cultured with the HIV-infected target cells at an E: T ratio of 2.5 :1, to evaluate their cytotoxic potential in the presence or absence of 10-1074 and 10-1074-Sia. The cell co-cultures were centrifuged at 200*g* for 2 min and incubated for 16 h at 37°C. Following incubation, 100 µl supernatant was removed from all wells and replaced with 100 µl Bright-Glo luciferase substrate reagent (Promega, Catalog # E2620). After 2 min, the well contents were mixed and transferred to a clear-bottom black 96-well microplate and luminescence (RLU) measurements were integrated over 1 second per well.

### Incucyte imaging

Uninfected or HIV_IIIB_-infected CEM.NKR CCR5^+^ Luc^+^ cells were stained 800 nM of cytolight rapid red dye (Incucyte, Catalog #4706) or 400 nM of cytolight rapid green dye (Incucyte, Catalog #4705) according to manufacturer’s instructions. Briefly, the cells were washed with PBS to remove serum and incubated with the indicated concentrations of dye in PBS at 37°C for 20 min. Following this, the cells were washed with 5x volume of 10% FBS containing RPMI 1640 media to quench and wash off unbound dye. The cells were resuspended in complete R10 and a mix of 20,000 uninfected (red) and HIV-infected (green) target cells, were added to 96-well flat bottom plates (10,000 cells each/well) and co-cultured with 3.75 nM of isotype control, 10-1074 or 10-1074-Sia antibodies for 2 h at 37°C for opsonization. After 2 h, primary monocytes from 4 HIV-negative donors were added as the effector cells (E :T=5 :1), and the cells were imaged on Incucyte® SX1 Live Cell Imager (Incucyte) at 1 h intervals for a time period of 16 h. % cytotoxicity was calculated based on the live red and green cell count at each time point using the Incucyte analysis software.

### Autologous cytotoxicity assays

Primary CD4^+^ T cells from six to seven HIV-negative donors were infected with HIV_TYBE_ as described earlier. Autologous monocytes and HIV_TYBE_-infected CD4^+^ T cells were cocultured at E:T ratio of 10:1 in the presence of 50 nM isotype control, 10-1074 or 10-1074-Sia, for 16 h at 37°C. Similarly, freshly isolated neutrophils were co-cultured with autologous HIV-infected target cells at an E: T ratio of 5: 1 in the presence of 10 nM isotype control, 10-1074 or 10-1074-Sia for 16 h at 37°C. Following incubation, the co-cultures were harvested and evaluated for intracellular p24 by staining with Zombie Aqua fixable viability dye (BioLegend), anti-CD3 BV421 (BioLegend, Clone UCHT1), and anti-p24 KC57-RD1 (Beckman Coulter).

### Transcriptomic analysis from autologous monocyte-HIV-infected CD4^+^ T cell co-cultures

Monocytes co-cultured with autologous HIV-infected CD4^+^ in the presence of 50 nM isotype control, 10-1074 and 10-1074-Sia for 16 h were harvested and RNA isolation was performed using the using RNAeasy micro kit (Qiagen, Catalog #74004) with on-column DNA digestion. Libraries were constructed on Stranded mRNA Library Prep Kit (Illumina) using 25 ng input RNA. The sequencing was performed on a NovaSeq X Plus 10B flowcell as 50bp single-end, and a minimum of 30M reads were captured for each sample. The RNA-seq data analysis was performed by Bencos Research Solutions Pvt. Ltd., Mumbai, India. For the analysis, the raw data obtained after performing the RNA sequencing were aligned to Reference Human Genome assembly GRCh38 using the salmon (v 1.10.1) software in conjunction with STAR aligner (v 2.7.10a) ^61^. Raw read counts were used as the input to perform differential gene expression analysis using the R package DESeq2 (v 1.34.0)^60^. DESeq2 normalized read counts were used to calculate the log2 fold change and adjusted p-values. P values were adjusted using Benjamini-Hochberg procedure to determine the False Discovery Rate (FDR) and control for multiple testing.

### Human memory CD4^+^ T cell isolation and engraftment into immunodeficient mice

Memory CD4^+^ T cells were isolated from PBMCs using EasySep human memory CD4^+^ T cell enrichment kit (StemCell Technologies, Catalog #19157), according to the manufacturer’s recommendation. Six-week-old NSG mice (The Wistar Institute, Philadelphia) were engrafted with 5 × 10^6^ memory CD4^+^ T cells via intravenous injection. Mice were bled weekly to assess human immune cell reconstitution and plasma viremia. Approximately 70 μL of peripheral blood was collected into EDTA-coated tubes via submandibular bleeding and processed immediately. Blood was centrifuged at 2500*g* for 10 min to separate plasma, which was stored at −80°C to further viral RNA extraction. Cell pellets were immediately stained for flow cytometry analysis as described below. One-week post-infection, mice were treated with 30 mg/kg of either 10-1074 or 10-1074-Sia conjugate, with 5 × 10^6^ effector cells, consisting of NK cells and monocytes in a 1:1 ratio, administered intravenously, once weekly for five consecutive weeks. The control group received only PBS using the same route. Mouse body weight was measured three times a week until the end of the study.

### Flow cytometric analysis

Samples were stained using a compensated panel of the following human-specific antibodies: anti-CD3 APC-Cy7 (Biolegend, Clone SK7), anti-CD4 Alexafluor647 (Biolegend, Clone OKT4), anti-CD8 BV421 (Biolegend, Clone SK1), anti-CD45 BUV395 (Invitrogen, Clone HI30), anti-CD56 PerCP-eFluor710 (Invitrogen, Clone TULY56), anti-CD14 FITC (Biolegend, clone M5E2), anti-CD16 BUV661 (Invitrogen, Clone CB16), anti-CD19 BV605 (Biolegend, Clone HIB19), CD45RA BV750 anti-(Biolegend, Clone HI100), anti-CD45RO PE-Cy7 (Biolegend, Clone UCHL1), Live/Dead BV510 (Zombie Aqua, Catalog #423102), and anti-mouse CD45 BUV805 (Invitrogen, Clone 30-F11) for 30 minutes at room temperature. After surface staining, red blood cells were lysed using ACK lysis buffer (Gibco, Catalog #A104920), and washed with PBS twice. Cells were then fixed, and fluorescent counting beads (Invitrogen; Catalog #C36950) were added to each sample to determine absolute cell count. Flow cytometric analysis of human cells was conducted using Symphony A5 and FlowJo version 10 software.

### In vivo HIV infection and analysis of infection

Mice were infected intravenously with 1500 TCID_50_ HIV_SUMA_ (a transmitter/founder virus). HIV infection in mice was monitored weekly by measuring HIV RNA copies in the plasma. Plasma samples from each time point was spiked in with a known quantity of RCAS virus to use as internal control, and RNA was isolated using QIAmp Viral RNA mini kit (Qiagen, Catalog #52906). cDNA was generated using SuperScript VILO Master Mix (Invitrogen, Catalog #11755500) on a thermal cycle at 25°C for 10 min, 42°C for 60 min, 85°C for 5 min, and 4°C. HIV quantitation was done from cDNA using Real-Time PCR. The reaction was carried out using Universal Master Mix II with UNG (Applied Biosystems, catalog #4440038) and following HIV-1 primers and probe: F522-43 Kumar-5’ GCCTCAATAAAGCTTGCCTTGA 3’, R626-43 Kumar-5’ GGGCGCCACTGCTAGAGA 3’, Probe Kumar (FAM/ZEN/BHQ-1)-5’ CCAGAGTCACACAACAGACGGGCACA 3’. The run was done on a Quant Studio 6 Real-Time PCR system using the following cycling parameters: 50°C for 2 min, 95°C for 10 min, followed by 60 cycles of 95°C for 15 s and 59°C for 1 min. Cycle threshold values were compared with validated HIV RNA standard run on each plate to determine HIV RNA copies.

### Measurement of plasma inflammatory markers

Plasma levels of human INF-γ, IP-10, TNF-α, MIP-1α, SDF-1a, IL-2, IL-6, IL-10, and IL-21 were determined using U-PLEX kits from Meso Scale Diagnostics (Biomarker Group 1 (hu) Assay; Catalog #K151AEM-2, Custom Immuno-Oncology Grp (hu) Assay; Catalog #K15067L-2, and Human MIG Antibody set; Catalog #F210I-3) according to the manufacturer’s instruction.

### Statistical Analysis

Statistical analysis was performed on GraphPad Prism, v.10 and R. The analysis used for each figure has been described in the corresponding figure legends.

## Abbreviations

10-1074-Sia: Sialidase conjugated 10-1074
ACK: Ammonium-Chloride-Potassium (lysis buffer)
ADCP: Antibody-dependent cellular phagocytosis
ART: Antiretroviral therapy
AUC: Area under the curve
BADG: Benzyl 2-acetamido-2-deoxy-α-D-galactopyranoside
bNAb: Broadly neutralizing antibodies
E:T: ratio Effector-to-target ratio
FACS: Fluorescence-activated cell sorting
Fc: Fragment crystallizable region (of an antibody)
FDR: False discovery rate
FITC: Fluorescein isothiocyanate
FPheK: 4-fluorophenyl carbamate lysine
IP10: Interferon gamma-induced protein 10 (CXCL10)
IPTG: Isopropyl β-d-1-thiogalactopyranoside
ITIMs: Immunoreceptor tyrosine-based inhibitory motifs
M-CSF: Macrophage colony-stimulating factor
MDMs: Monocyte-derived macrophages
NK: cells Natural Killer cells
PBS: Phosphate-buffered saline
PLWH: People Living with HIV
PI3K: Phosphoinositide 3-kinase
qPCR: Quantitative polymerase chain reaction
RCAS: Replication-competent avian sarcoma-leukosis virus
RLU: Relative luminescence units
RSEM: RNA-Seq by Expectation-Maximization
Siglecs: Sialic acid-binding immunoglobulin-like lectins
TGF-β: Transforming growth factor-beta
TILs: Tumor infiltrating lymphocytes
TNF-α: Tumor necrosis factor-alpha
U-PLEX: Multiplex assay platform from Meso Scale Diagnostics

## SUPPLEMENTARY FIGURE LEGENDS

**Supplementary Figure 1. Cytotoxicity of PBMCs against HIV-infected CEM.NKR CCR5^+^ Luc^+^ target cells.** HIV-infected CEM.NKR CCR5^+^ Luc^+^ target cells were co-cultured with PBMCs isolated from three HIV-negative donors (E:T ratio = 100:1) in the presence of increasing concentrations of 10-1074 or 10-1074-Sia for 16 hours. Luminescence was measured after 16 hours. **(a)** Total cytotoxicity across a range of antibody concentrations (left panel). **(b)** Antibody-mediated cytotoxicity, calculated as total cytotoxicity minus direct cytotoxicity, across a range of antibody concentrations (left panel). AUCs (right panel) were computed to demonstrate cytotoxicity over multiple antibody concentrations. Statistical analysis was performed using paired t-tests.

**Supplementary Figure 2. Expression of Siglec receptors on CD8^+^ and** γδ **T cells.** Expression of Siglec-3, Siglec-7 and Siglec-9 receptors on purified CD8^+^ T cells **(a)** and γδ T cells **(b)** was measured by flow cytometry.

Supplementary Figure 3. Total cytotoxicity of primary NK cells and distinct myeloid cell types against HIV-infected CEM.NKR CCR5^+^ Luc^+^. Total cytotoxicity by primary NK cells (a), monocytes (b), MDMs (c) and neutrophils (d) against HIV_IIIB_-infected CEM.NKR CCCR5^+^ Luc^+^ cells were measured in the presence of 10-1074 or 10-1074-Sia. Left panels show the cytotoxicity across multiple dosage of the antibody, while right panel depicts the impact as calculated by AUC. Statistical analyses were performed by paired t-test.

**Supplementary Figure 4. Gating strategy for autologous cytotoxicity assays.** Representative contour plots show the gating strategy used for flow cytometric analysis of intracellular p24 for autologous cytotoxicity assays using monocyte and neutrophil effector cells.

**Supplementary Figure 5. Gating strategy for evaluating the purity of memory CD4^+^ T cells prior to engraftment in NSG mice.** Representative flow cytometry contour plots show the gating strategy used for evaluating the purity of memory CD4^+^ T cells before engraftment in NSG mice.

**Supplementary Figure 6. Gating strategy to examine the engraftment of memory CD4^+^ T cell in humanized mice.** Representative contour plots to show the successful reconstitution of human memory CD4^+^ T cells in NSG at day 34 post engraftment, as assessed by flow cytometry. hCD45^+^ CD4^+^ population was measured at day 34 post engraftment.

**Supplementary Figure 7. Purity check of monocytes and NK cells before treatment in humanized mice. (a-b)** Purity of monocytes **(a)** and NK cells **(b)** measured by flow cytometry post isolation by surface staining with either anti-CD14 and anti-CD16 antibodies or anti-CD56 and anti-CD16 antibodies.

## SUPPLEMENTARY TABLES

**Supplementary Table 1. List of glycosylation related genes.**

**Supplementary Table 2: DNA sequence of oligonucleotides**

**Supplementary Table 3: DNA sequence of FB-sia protein**

## AUTHOR CONTRIBUTIONS

M.A-M and H.X conceived and designed the study. S.S, S.M.S, R.L., L.B.G, P.S. O.S.A carried out the experiments. A.D, P.W.D, B.J interpreted data. S.S, S.M.S, and M.A-M wrote the manuscript, and all authors edited it.

## ACKNOWLEDGMENTS

This study is supported by NIH R01AI165079 to M.A-M and H.X. M.A-M is also supported by NIH grants (R01AA029859, R01DK123733, R01AG062383, and R01NS117458). M.A-M is also funded by the NIH-funded BEAT-HIV Martin Delaney Collaboratory to cure HIV-1 infection (1UM1Al126620). P.W.D. has research time supported by P20GM103427 and R15AI178516. We Would like to thank Drs. Michel Nussenzweig, Costin Tomescu, and Luis J. Montaner for providing the wild-type 10-1074.

