## Supplementary figures and images for "HIV-Induced Sialoglycans on Infected Cells Promote Immune Evasion from Myeloid Cell-Mediated Killing"

### Supplementary Figure 1

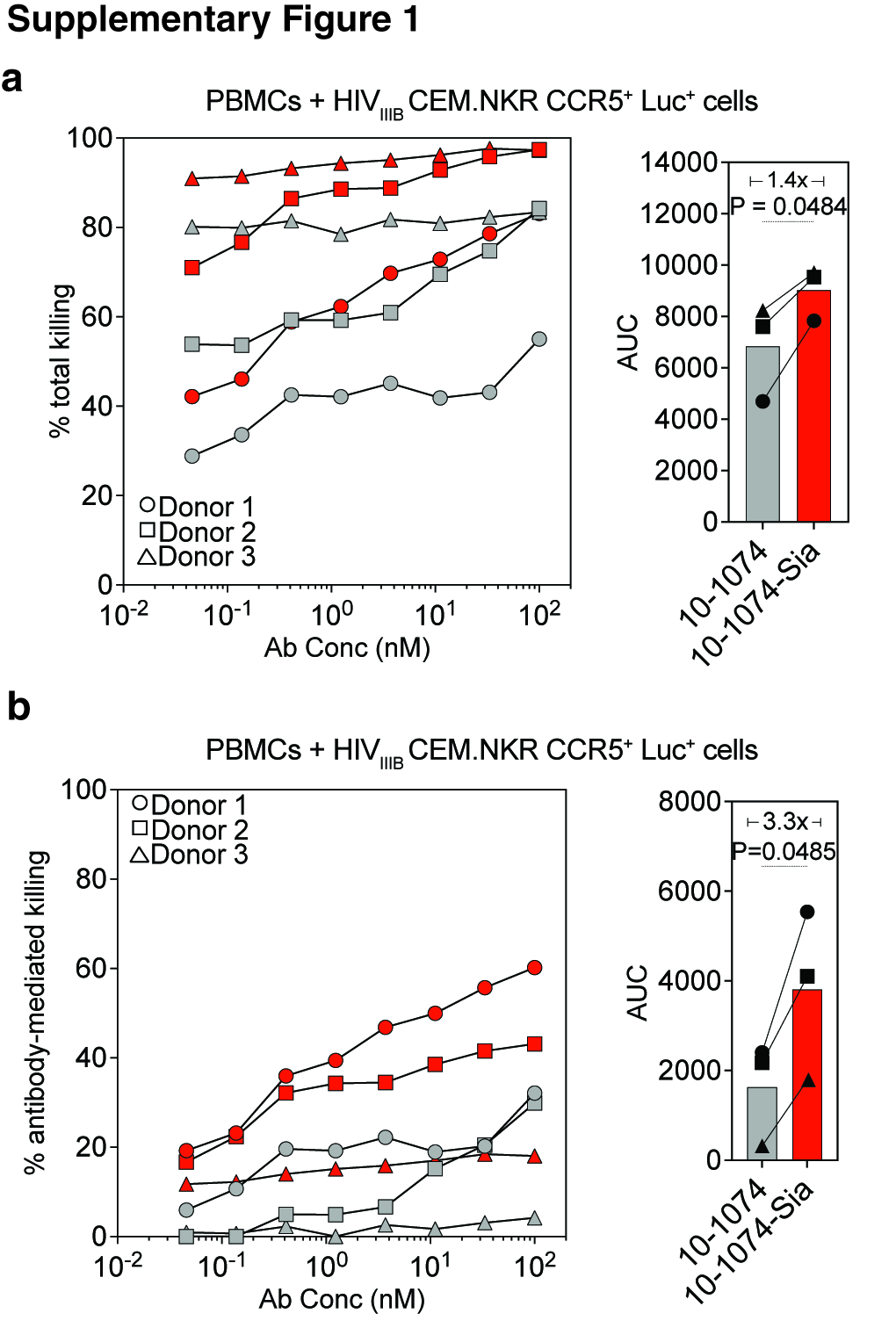

### Supplementary Figure 2

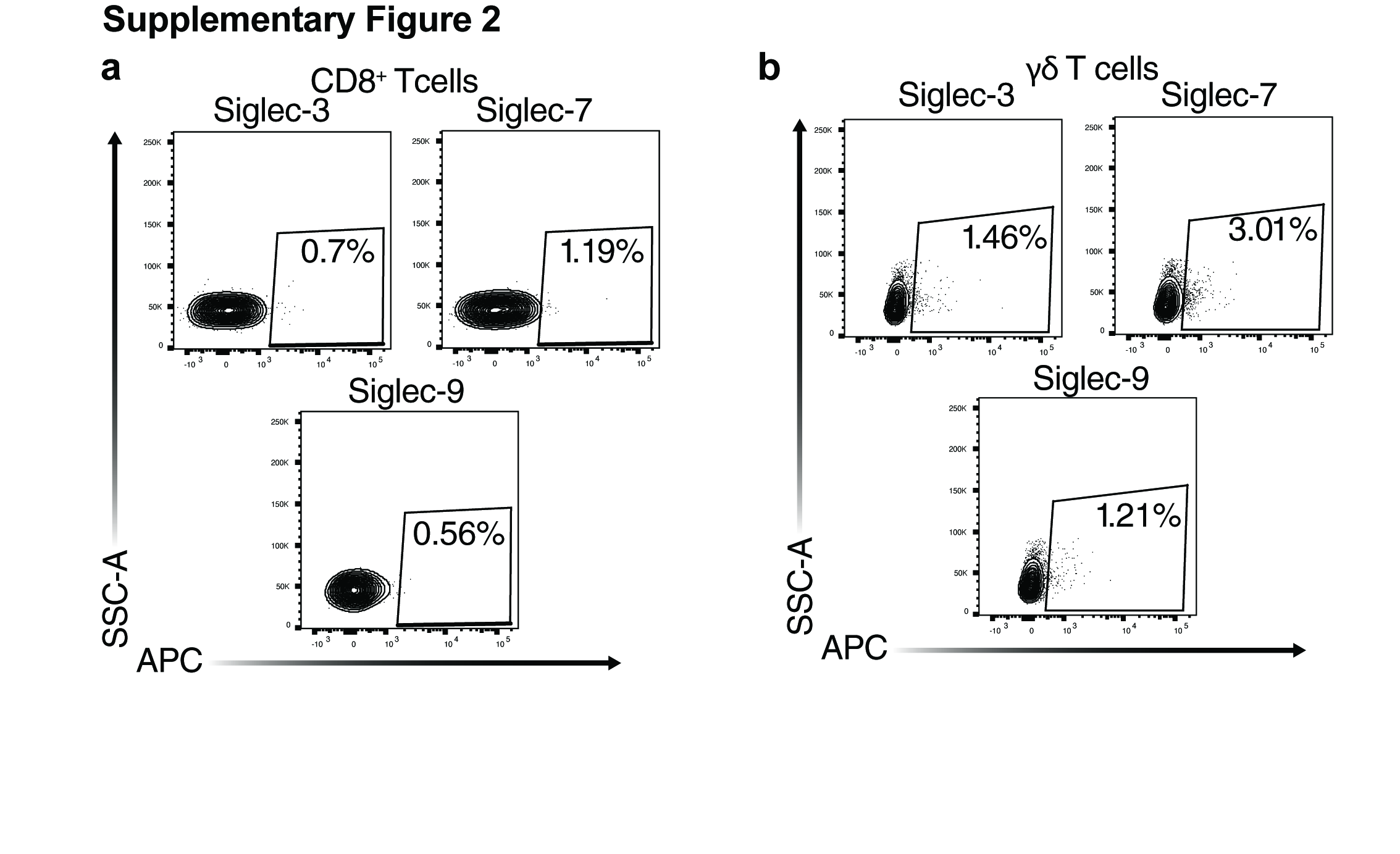

### Supplementary Figure 3

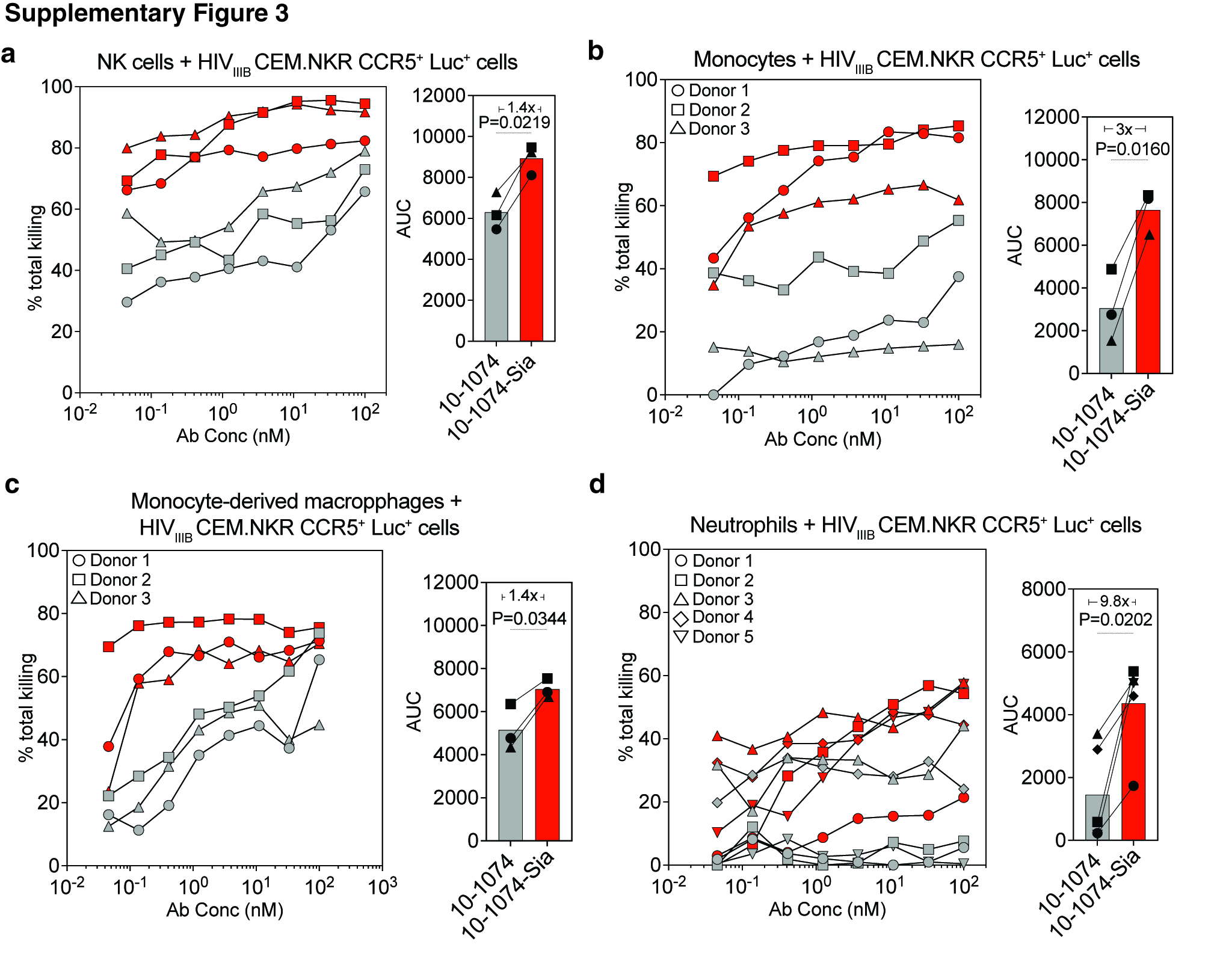

### Supplementary Figure 4

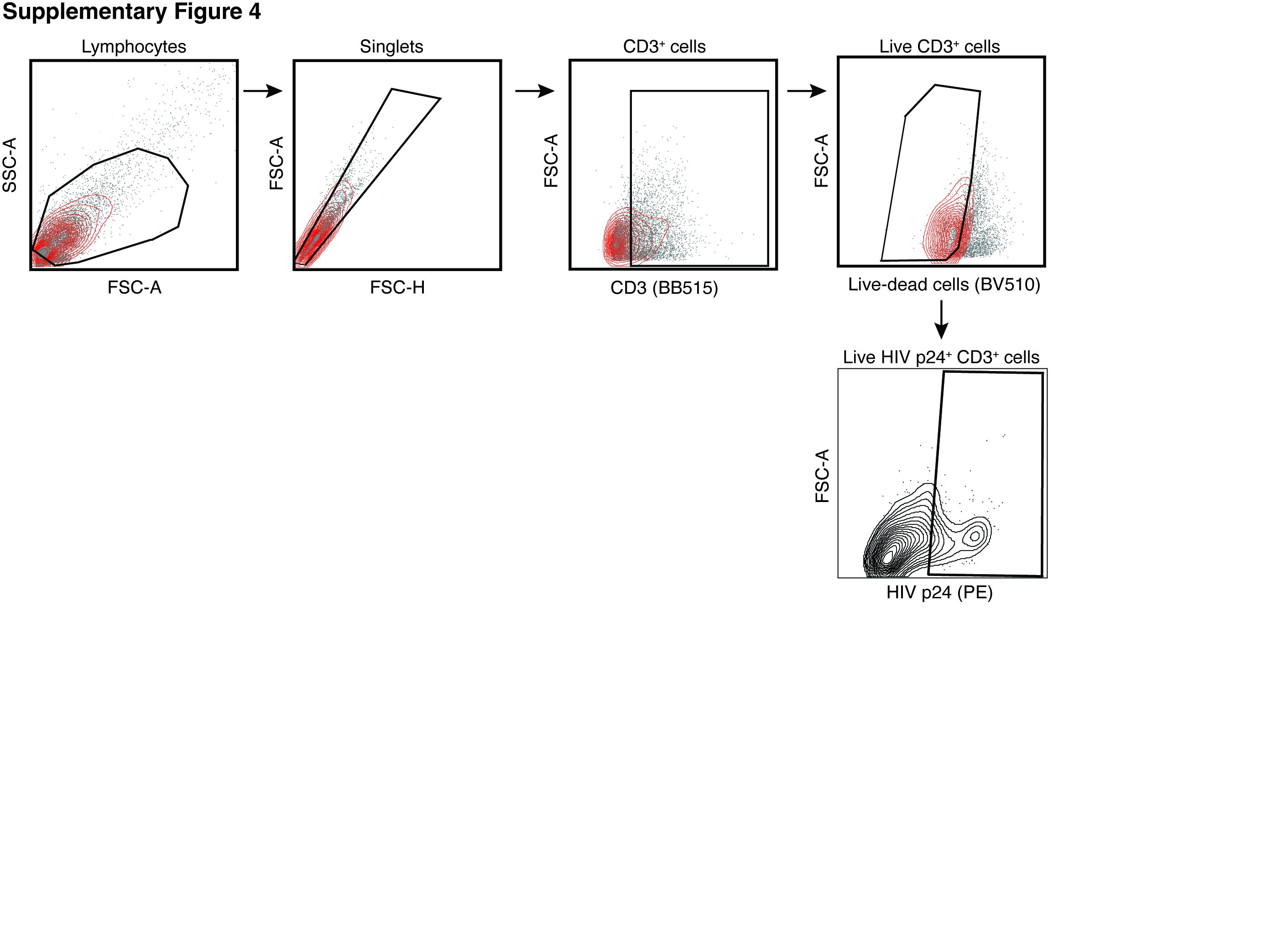

### Supplementary Figure 5

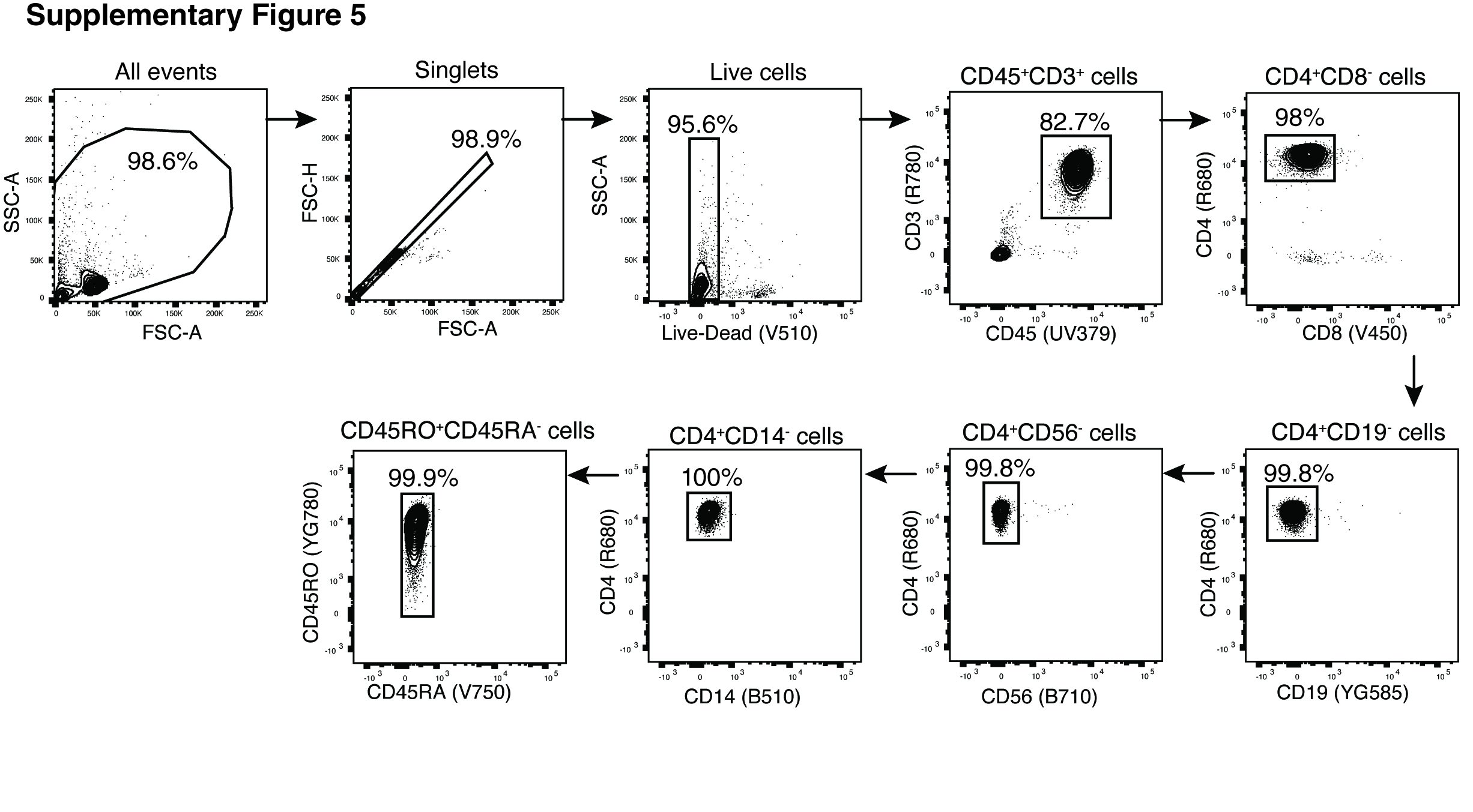

### Supplementary Figure 6

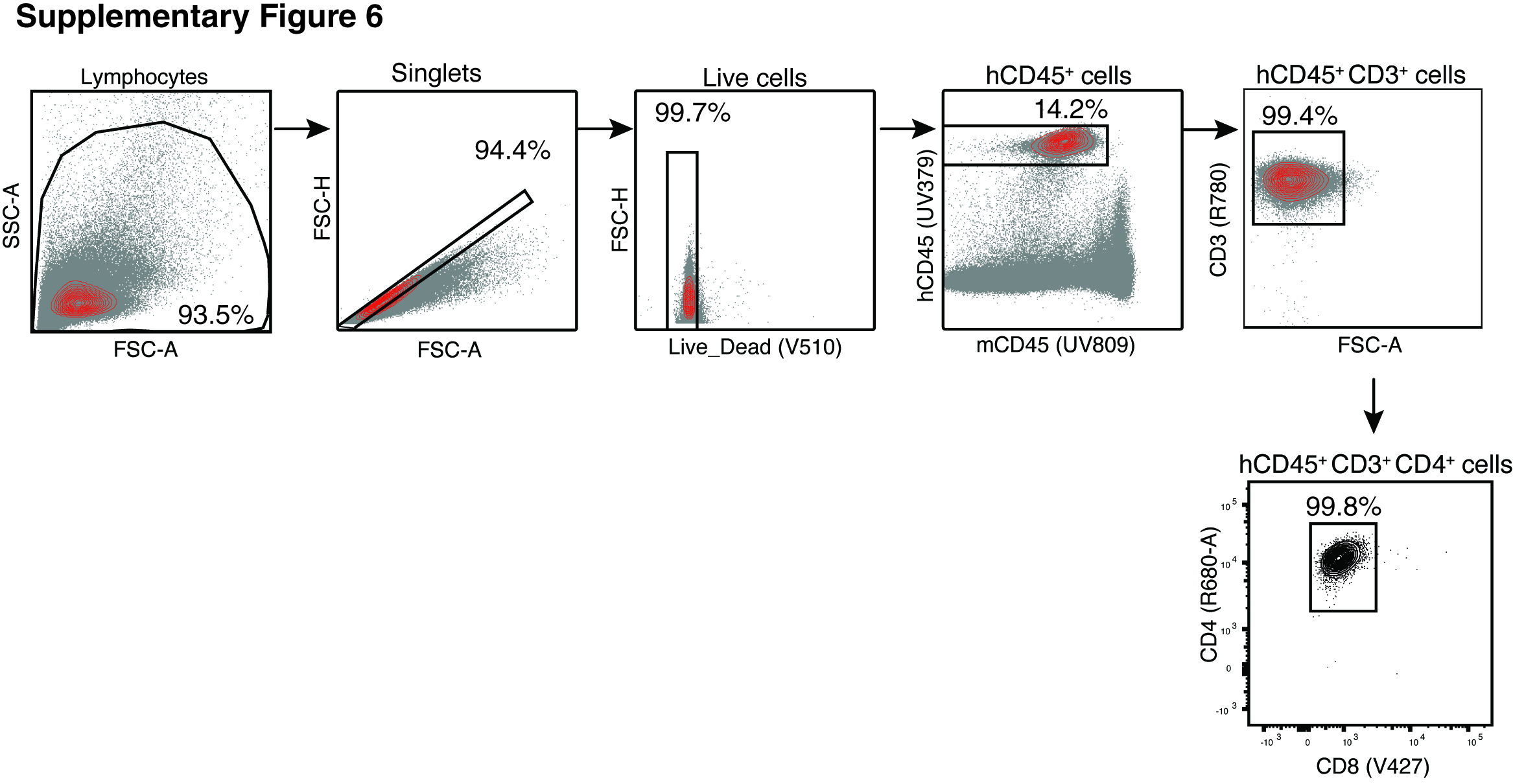

### Supplementary Figure 7

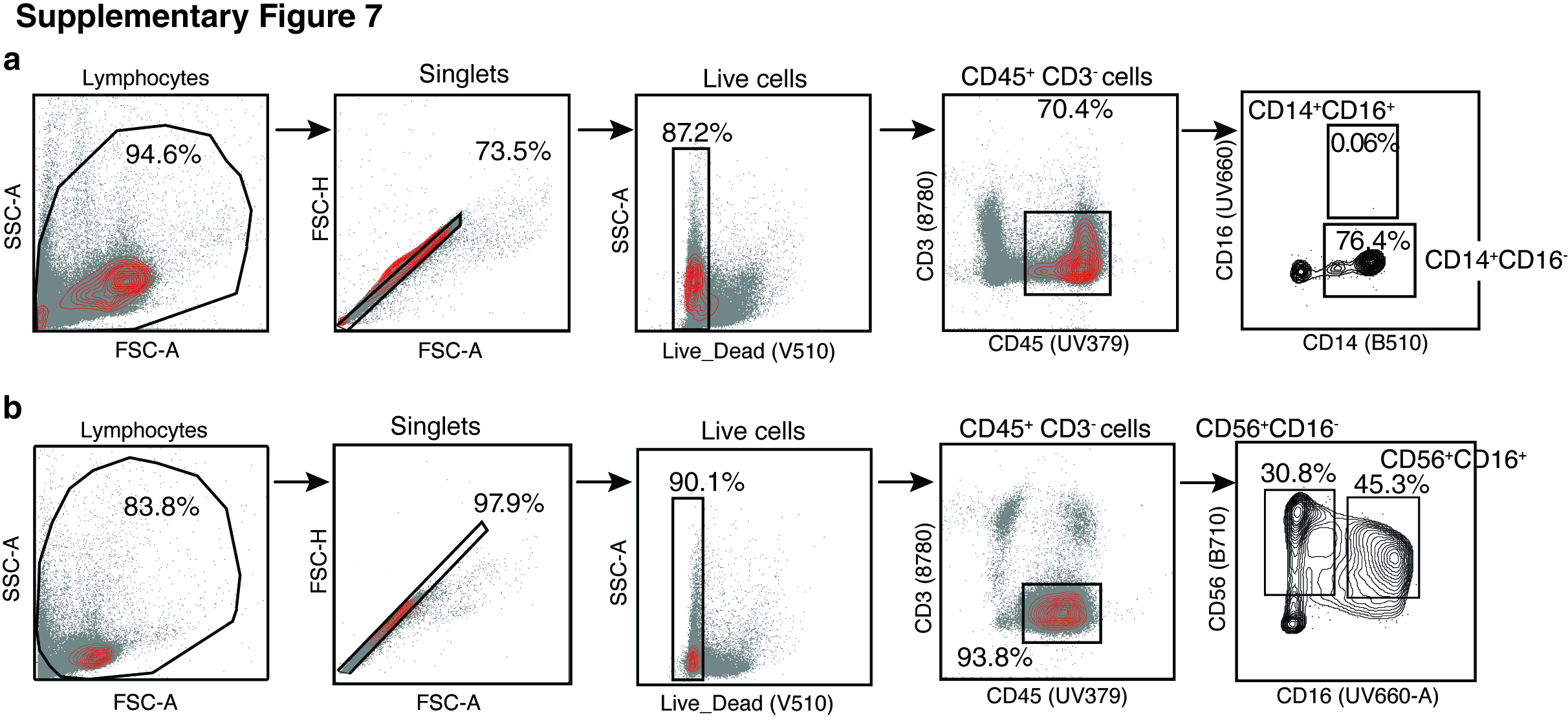
