## Supplementary Table 1 for "HIV-Induced Sialoglycans on Infected Cells Promote Immune Evasion from Myeloid Cell-Mediated Killing"

**Supplementary Table 1.** List of glycosylation related genes

| # | Name | UniProt | Ensembl ID | Full name | Activity/Function | Primary Monosaccharide/ Sugar | Pathway |
|---|---|---|---|---|---|---|---|
| 1 | A3GALT2 | U3KPV4 | ENSG00000184389 | Alpha 1,3-galactosyltransferase 2 | Inactive/Putative | Galactose | Inactive/Putative |
| 2 | A4GALT | Q9NPC4 | ENSG00000128274 | Alpha 1,4-galactosyltransferase | Transferase | Galactose | Glycolipid |
| 3 | A4GNT | Q9UNA3 | ENSG00000118017 | Alpha-1,4-N-acetylglucosaminyltransferase | Transferase | N-acetylglucosamine (GlcNAc) | N-acetylglucosamine (GlcNAc) |
| 4 | ABO | P16442 | ENSG00000175164 | ABO blood group (transferase A, alpha 1-3-N-acetylgalactosaminyltransferase; transferase B, alpha 1-3-galactosyltransferase) | Transferase | More than one monosacchrides | Galactose/N-acetylgalactosamine (GalNAc) |
| 5 | ACAN | P16112 | ENSG00000157766 | Aggrecan | Glycan Binding/lectin | Hyaluronic acid | Proteoglycan |
| 6 | AGA | P20933 | ENSG00000038002 | Aspartylglucosaminidase | Glycosidase | N-acetylglucosamine (GlcNAc) | Cleaves the GlcNAc-Asn bond |
| 7 | ALG1 | Q9BT22 | ENSG00000033011 | Asparagine-linked glycosylation 1, beta-1,4-mannosyltransferase homolog (S. cerevisiae) | Transferase | Mannose | N-glycan |
| 8 | ALG10 | Q5BKT4 | ENSG00000139133 | Asparagine-linked glycosylation 10, alpha-1,2-glucosyltransferase homolog (S. pombe) | Transferase | Glucose | N-glycan |
| 9 | ALG10B | Q5I7T1 | ENSG00000175548 | Asparagine-linked glycosylation 10, alpha-1,2-glucosyltransferase homolog B (yeast) | Transferase | Glucose | N-glycan |
| 10 | ALG11 | Q2TAA5 | ENSG00000253710 | Asparagine-linked glycosylation 11, alpha-1,2-mannosyltransferase homolog (yeast) | Transferase | Mannose | N-glycan |
| 11 | ALG12 | Q9BV10 | ENSG00000182858 | Asparagine-linked glycosylation 12, alpha-1,6-mannosyltransferase homolog (S. cerevisiae) | Transferase | Mannose | N-glycan |
| 12 | ALG13 | Q9NP73 | ENSG00000101901 | Asparagine-linked glycosylation 13 homolog (S. cerevisiae) | Inactive/Putative | Inactive/Putative | N-glycan |
| 13 | ALG14 | Q96F25 | ENSG00000172339 | Asparagine-linked glycosylation 14 homolog (S. cerevisiae) | Transferase | N-acetylglucosamine (GlcNAc) | N-glycan |
| 14 | ALG1L | Q6GMV1 | - | Putative glycosyltransferase ALG1-like | Inactive/Putative | Inactive/Putative | N-glycan |
| 15 | ALG1L2 | C9J202 | - | Putative glycosyltransferase ALG1L2 | Inactive/Putative | Inactive/Putative | N-glycan |
| 16 | ALG2 | O75340 | ENSG00000119523 | Asparagine-linked glycosylation 2, alpha-1,3-mannosyltransferase homolog (S. cerevisiae) | Transferase | Mannose | N-glycan |
| 17 | ALG3 | Q92685 | ENSG00000214160 | Asparagine-linked glycosylation 3, alpha-1,3- mannosyltransferase homolog (S. cerevisiae) | Transferase | Mannose | N-glycan |
| 18 | ALG5 | Q9Y673 | ENSG00000120697 | Asparagine-linked glycosylation 5, dolichyl-phosphate beta-glucosyltransferase homolog (S. cerevisiae) | Transferase | Glucose | N-glycan |
| 19 | ALG6 | Q9Y672 | ENSG00000088035 | Asparagine-linked glycosylation 6, alpha-1,3-glucosyltransferase homolog (S. cerevisiae) | Transferase | Glucose | N-glycan |
| 20 | ALG8 | Q9BVK2 | ENSG00000159063 | Asparagine-linked glycosylation 8, alpha-1,3-glucosyltransferase homolog (S. cerevisiae) | Transferase | Glucose | N-glycan |
| 21 | ALG9 | Q9H6U8 | ENSG00000086848 | Asparagine-linked glycosylation 9, alpha-1,2-mannosyltransferase homolog (S. cerevisiae) | Transferase | Mannose | N-glycan |
| 22 | AMDHD2 | Q9Y303 | ENSG00000162066 | N-acetylglucosamine-6-phosphate deacetylase | Glycosidase | Acetate | Nucleotide_Sugar_Synthesis,UDP-GlcNAc_Synthesis |
| 23 | ARSB | P15848 | ENSG00000113273 | Arylsulfatase B | Sulfatase | Sulfate | CS_Lysosome_Degradation,DS_Lysosome_Degradation |
| 28 | ARSG | Q96EG1 | ENSG00000141337 | Arylsulfatase G | Sulfatase | Sulfate | HS_Lysosome_Degradation |
| 29 | ARSK | Q6UWY0 | ENSG00000164291 | Arylsulfatase K | Sulfatase | Sulfate | HS_Lysosome_Degradation,DS_Lysosome_Degradation,CS_Lysosome_Degradation |
| 24 | ASGR1 | P07306 | ENSG00000141505 | Asialoglycoprotein receptor 1 | Glycan Binding/lectin | Sialic Acid | Sialic acid removed glycoproteins |
| 25 | ASGR2 | P07307 | ENSG00000161944 | Asialoglycoprotein receptor 2 | Glycan Binding/lectin | Sialic Acid | Sialic acid removed glycoproteins |
| 26 | ATRN | O75882 | ENSG00000088812 | Attractin | Glycan Binding/lectin | Not Specific | Specific glycan ligand not known |
| 27 | ATRNL1 | Q5VV63 | ENSG00000107518 | Attractin-like 1 | Glycan Binding/lectin | Not Specific | Specific glycan ligand not known |
| 37 | B3GALTL | Q6Y288 | ENSG00000187676 | Beta 1,3-galactosyltransferase-like | Transferase | Galactose | O-glycan |
| 30 | B3GALNT1 | O75752 | ENSG00000169255 | Beta-1,3-N-acetylgalactosaminyltransferase 1 (globoside blood group) | Transferase | N-acetylgalactosamine (GalNAc) | Glycolipid |
| 31 | B3GALNT2 | Q8NCR0 | ENSG00000162885 | Beta-1,3-N-acetylgalactosaminyltransferase 2 | Transferase | N-acetylgalactosamine (GalNAc) | O-glycan/N-glycan |
| 32 | B3GALT1 | Q9Y5Z6 | ENSG00000172318 | UDP-Gal:betaGlcNAc beta 1,3-galactosyltransferase, polypeptide 1 | Transferase | Galactose | Type I LacNAc, N-linked, O-GalNAc |
| 33 | B3GALT2 | O43825 | ENSG00000162630 | UDP-Gal:betaGlcNAc beta 1,3-galactosyltransferase, polypeptide 2 | Transferase | Galactose | Type I LacNAc, N-linked, O-GalNAc |
| 34 | B3GALT4 | O96024 | ENSG00000235863 | UDP-Gal:betaGlcNAc beta 1,3-galactosyltransferase, polypeptide 4 | Transferase | Galactose | Glycolipids,Ganglioside,GPI_Linked |
| 35 | B3GALT5 | Q9Y2C3 | ENSG00000183778 | UDP-Gal:betaGlcNAc beta 1,3-galactosyltransferase, polypeptide 5 | Transferase | Galactose | Glycolipids,Lacto,Core3 |
| 36 | B3GALT6 | Q96L58 | ENSG00000176022 | UDP-Gal:betaGal beta 1,3-galactosyltransferase polypeptide 6 | Transferase | Galactose | GAG,O-Xyl |
| 38 | B3GAT1 | Q9P2W7 | ENSG00000109956 | Beta-1,3-glucuronyltransferase 1 (glucuronosyltransferase P) | Transferase | Glucuronic acid (GlcA) | Capping, HNK-1 |
| 39 | B3GAT2 | Q9NPZ5 | ENSG00000112309 | Beta-1,3-glucuronyltransferase 2 (glucuronosyltransferase S) | Transferase | Glucuronic acid (GlcA) | Capping, HNK-1 |
| 40 | B3GAT3 | O94766 | ENSG00000149541 | Beta-1,3-glucuronyltransferase 3 (glucuronosyltransferase I) | Transferase | Glucuronic acid (GlcA) | GAG,O-Xyl |
| 41 | B3GLCT | Q6Y288 | ENSG00000187676 | Beta-1,3-glucosyltransferase | Transferase | Glucose | O-linked fucosylglycan |
| 42 | B3GNT1/ B4GAT1 | O43505 | ENSG00000174684 | UDP-GlcNAc:betaGal beta-1,3-N-acetylglucosaminyltransferase 1/ | Transferase | Glucuronic acid (GlcA) | O-mannosylation |
| 43 | B3GNT2 | Q9NY97 | ENSG00000170340 | UDP-GlcNAc:betaGal beta-1,3-N-acetylglucosaminyltransferase 2 | Transferase | N-acetylglucosamine (GlcNAc) | poly-N-acetyllactosamine |
| 44 | B3GNT3 | Q9Y2A9 | ENSG00000179913 | UDP-GlcNAc:betaGal beta-1,3-N-acetylglucosaminyltransferase 3 | Transferase | N-acetylglucosamine (GlcNAc) | O-GalNAc, Core1 |
| 45 | B3GNT4 | Q9C0J1 | ENSG00000176383 | UDP-GlcNAc:betaGal beta-1,3-N-acetylglucosaminyltransferase 4 | Transferase | N-acetylglucosamine (GlcNAc) | Elongation_and_Branching, Type_II_LacNAc |
| 46 | B3GNT5 | Q9BYG0 | ENSG00000176597 | UDP-GlcNAc:betaGal beta-1,3-N-acetylglucosaminyltransferase 5 | Transferase | N-acetylglucosamine (GlcNAc) | GlcCer,Lacto,Neolacto |
| 47 | B3GNT6 | Q6ZMB0 | ENSG00000198488 | UDP-GlcNAc:betaGal beta-1,3-N-acetylglucosaminyltransferase 6 (core 3 synthase) | Transferase | N-acetylglucosamine (GlcNAc) | O-GalNAc, Core3 |
| 48 | B3GNT7 | Q8NFL0 | ENSG00000156966 | UDP-GlcNAc:betaGal beta-1,3-N-acetylglucosaminyltransferase 7 | Transferase | N-acetylglucosamine (GlcNAc) | Elongation_and_Branching, Type_II_LacNAc,KS_Polymerization |
| 49 | B3GNT8 | Q7Z7M8 | ENSG00000177191 | UDP-GlcNAc:betaGal beta-1,3-N-acetylglucosaminyltransferase 8 | Transferase | N-acetylglucosamine (GlcNAc) | Elongation_and_Branching, Type_II_LacNAc |
| 50 | B3GNTL1 | Q67FW5 | ENSG00000175711 | UDP-GlcNAc:betaGal beta-1,3-N-acetylglucosaminyltransferase-like 1 | Inactive/Putative | Inactive/Putative | Mucin Type O-glycosylation |
| 51 | B4GALNT1 | Q00973 | ENSG00000135454 | Beta-1,4-N-acetyl-galactosaminyl transferase 1 | Transferase | N-acetylgalactosamine (GalNAc) | gangliosides |
| 52 | B4GALNT2 | Q8NHY0 | ENSG00000167080 | Beta-1,4-N-acetyl-galactosaminyl transferase 2 | Transferase | N-acetylgalactosamine (GalNAc) | Capping,SdaAntigen |
| 53 | B4GALNT3 | Q6L9W6 | ENSG00000139044 | Beta-1,4-N-acetyl-galactosaminyl transferase 3 | Transferase | N-acetylgalactosamine (GalNAc) | Capping,LacDiNAc |
| 54 | B4GALNT4 | Q76KP1 | ENSG00000182272 | Beta-1,4-N-acetyl-galactosaminyl transferase 4 | Transferase | N-acetylgalactosamine (GalNAc) | Capping,LacDiNAc |
| 55 | B4GALT1 | P15291 | ENSG00000086062 | UDP-Gal:betaGlcNAc beta 1,4- galactosyltransferase, polypeptide 1 | Transferase | Galactose | Type II LacNAc,N-linked, O-GalNAc,POFUT1_Type,KS_Polymerization |
| 56 | B4GALT2 | O60909 | ENSG00000117411 | UDP-Gal:betaGlcNAc beta 1,4- galactosyltransferase, polypeptide 2 | Transferase | Galactose | Type II LacNAc, N-linked, O-GalNAc |
| 57 | B4GALT3 | O60512 | ENSG00000158850 | UDP-Gal:betaGlcNAc beta 1,4- galactosyltransferase, polypeptide 3 | Transferase | Galactose | Type II LacNAc, N-linked, O-GalNAc, KS_Polymerization |
| 58 | B4GALT4 | O60513 | ENSG00000121578 | UDP-Gal:betaGlcNAc beta 1,4- galactosyltransferase, polypeptide 4 | Transferase | Galactose | Type II LacNAc, N-linked, O-GalNAc, KS_Polymerization |
| 59 | B4GALT5 | O43286 | ENSG00000158470 | UDP-Gal:betaGlcNAc beta 1,4- galactosyltransferase, polypeptide 5 | Transferase | Galactose | Type II LacNAc, N-linked, O-GalNAc |
| 60 | B4GALT6 | Q9UBX8 | ENSG00000118276 | UDP-Gal:betaGlcNAc beta 1,4- galactosyltransferase, polypeptide 6 | Transferase | Galactose | GlcCer, Lacto,Neolacto,Ganglioside,Globo |
| 61 | B4GALT7 | Q9UBV7 | ENSG00000027847 | Xylosylprotein beta 1,4-galactosyltransferase, polypeptide 7 (galactosyltransferase I) | Transferase | Galactose | GAG,O-Xyl |
| 62 | BCAN | Q96GW7 | ENSG00000132692 | Brevican | Glycan Binding/lectin | Hyaluronic acid | |
| 63 | B4GAT1 | O43505 | ENSG00000174684 | Beta-1,4-glucuronyltransferase 1 | Transferase | Glucuronic acid (GlcA) | O-mannosylation |
| 64 | C1GALT1 | Q9NS00 | ENSG00000106392 | Core 1 synthase, glycoprotein-N-acetylgalactosamine 3-beta-galactosyltransferase, 1 | Transferase | Galactose | core 1 O-glycan |
| 65 | C1GALT1C1 | Q96EU7 | ENSG00000171155 | C1GALT1-specific chaperone 1 | chaperone | Not Specific | Synthesis of T antigen |
| 66 | CALR | P27797 | ENSG00000179218 | Calreticulin | chaperone | Not Specific | Glycosylation_Regulation |
| 67 | CALR3 | Q96L12 | ENSG00000269058 | Calreticulin 3 | chaperone | Not Specific | Spermatogenesis, Lectin-independent chaperone for specific client proteins such as ADAM3. Required for sperm fertility |
| 68 | CANX | P27824 | ENSG00000127022 | Calnexin | chaperone | Not Specific | Glycosylation_Regulation |
| 69 | CASD1 | Q96PB1 | ENSG00000127995 | N-acetylneuraminate 9-O-acetyltransferase | Transferase | Sialic Acid | 9-O-acetylation of sialic acids |
| 70 | CD207 | Q9UJ71 | ENSG00000116031 | CD207 molecule, langerin | Glycan Binding/lectin | Mannose | sulfated, mannosylated glycans |
| 71 | CD209 | Q9NNX6 | ENSG00000090659 | CD209 molecule | Glycan Binding/lectin | Mannose | mannose-like carbohydrates |
| 72 | CD33L3 | Q6ZMC9 | ENSG00000197046 | Sialic acid binding Ig-like lectin 15 | Glycan Binding/lectin | Sialic Acid | |
| 73 | CD69 | Q07108 | ENSG00000110848 | Early activation antigen CD69 | Glycan Binding/lectin | Not Specific | Gal-1 |
| 74 | CD72 | P21854 | ENSG00000137101 | B-cell differentiation antigen CD72 | Glycan Binding/lectin | Not Specific | RNA-related lupus self-antigen Sm/RNP, Glycan Ligand not clearly known |
| 75 | CEMIP2 | Q9UHN6 | ENSG00000135048 | Cell surface hyaluronidase | Inactive/Putative | Inactive/Putative | Inactive/Putative |
| 76 | ChGn | Q8TDX6 | ENSG00000147408 | Chondroitin sulfate N-acetylgalactosaminyltransferase 1 | Transferase | N-acetylgalactosamine (GalNAc) | chondroitin chain biosynthesis |
| 77 | CHODL | Q9H9P2 | ENSG00000154645 | Chondrolectin | Glycan Binding/lectin | Not Specific | ? |
| 78 | CHPF | Q8IZ52 | ENSG00000123989 | Chondroitin polymerizing factor | Transferase | N-acetylgalactosamine (GalNAc) | chondroitin polymerization |
| 79 | CHPF2 | Q9P2E5 | ENSG00000033100 | Chondroitin sulfate glucuronyltransferase | Transferase | Glucuronic acid (GlcA) | chondroitin polymerization |
| 80 | CHST1 | O43916 | ENSG00000175264 | Carbohydrate (keratan sulfate Gal-6) sulfotransferase 1 | Transferase | Sulfate | KS_Sulfation,O-GalNAc |
| 81 | CHST10 | O43529 | ENSG00000115526 | Carbohydrate sulfotransferase 10 | Transferase | Sulfate | Capping,HNK-1 |
| 82 | CHST11 | Q9NPF2 | ENSG00000171310 | Carbohydrate (chondroitin 4) sulfotransferase 11 | Transferase | Sulfate | CS&DS_Sulfation |
| 83 | CHST12 | Q9NRB3 | ENSG00000136213 | Carbohydrate (chondroitin 4) sulfotransferase 12 | Transferase | Sulfate | CS&DS_Sulfation |
| 84 | CHST13 | Q8NET6 | ENSG00000180767 | Carbohydrate (chondroitin 4) sulfotransferase 13 | Transferase | Sulfate | CS&DS_Sulfation |
| 85 | CHST14 | Q8NCH0 | ENSG00000169105 | Carbohydrate (N-acetylgalactosamine 4-0) sulfotransferase 14 | Transferase | Sulfate | CS&DS_Sulfation |
| 86 | CHST15 | Q7LFX5 | ENSG00000182022 | Carbohydrate sulfotransferase 15 | Transferase | Sulfate | CS&DS_Sulfation |
| 87 | CHST2 | Q9Y4C5 | ENSG00000175040 | Carbohydrate (N-acetylglucosamine-6-O) sulfotransferase 2 | Transferase | Sulfate | KS_Sulfation,O-GalNAc,Core2, 6-Sulfo_sLeX |
| 88 | CHST3 | Q7LGC8 | ENSG00000122863 | Carbohydrate (chondroitin 6) sulfotransferase 3 | Transferase | Sulfate | CS&DS_Sulfation,KS_Sulfation,Type_II_LacNAc |
| 89 | CHST4 | Q8NCG5 | ENSG00000140835 | Carbohydrate (N-acetylglucosamine 6-O) sulfotransferase 4 | Transferase | Sulfate | O-GalNAc,Core2 |
| 90 | CHST5 | Q9GZS9 | ENSG00000135702 | Carbohydrate (N-acetylglucosamine 6-O) sulfotransferase 5 | Transferase | Sulfate | KS_Sulfation,O-GalNAc,Core2, 6-Sulfo_sLeX |
| 91 | CHST6 | Q9GZX3 | ENSG00000183196 | Carbohydrate (N-acetylglucosamine 6-O) sulfotransferase 6 | Transferase | Sulfate | KS_Sulfation,O-GalNAc,Core2, 6-Sulfo_sLeX |
| 92 | CHST7 | Q9NS84 | ENSG00000147119 | Carbohydrate (N-acetylglucosamine 6-O) sulfotransferase 7 | Transferase | Sulfate | CS&DS_Sulfation |
| 93 | CHST8 | Q9H2A9 | ENSG00000124302 | Carbohydrate (N-acetylgalactosamine 4-0) sulfotransferase 8 | Transferase | Sulfate | Capping,LacDiNAc |
| 94 | CHST9 | Q7L1S5 | ENSG00000154080 | Carbohydrate (N-acetylgalactosamine 4-0) sulfotransferase 9 | Transferase | Sulfate | Capping,LacDiNAc |
| 95 | CHSY1 | Q86X52 | ENSG00000131873 | Chondroitin sulfate synthase 1 | Transferase | More than one monosacchrides | Glucuronic acid, N-acetylgalactosamine |
| 96 | CHSY3 | Q70JA7 | ENSG00000198108 | Chondroitin sulfate synthase 3 | Transferase | More than one monosacchrides | Glucuronic acid, N-acetylgalactosamine |
| 98 | CLEC10A | Q8IUN9 | ENSG00000132514 | C-type lectin domain family 10, member A | Glycan Binding/lectin | More than one monosacchrides | Galctose and N-acetylgalactosamine |
| 99 | CLEC11A | Q9Y240 | ENSG00000105472 | C-type lectin domain family 11, member A | Glycan Binding/lectin | Not Specific | - |
| 100 | CLEC12A | Q5QGZ9 | ENSG00000172322 | C-type lectin domain family 12, member A | Glycan Binding/lectin | Not Specific | uric acid crystals (cell death danger signals) |
| 101 | CLEC12B | Q2HXU8 | ENSG00000256660 | C-type lectin domain family 12, member B | Glycan Binding/lectin | Not Specific | - |
| 102 | CLEC14A | Q86T13 | ENSG00000176435 | C-type lectin domain family 14, member A | Glycan Binding/lectin | Not Specific | - |
| 103 | CLEC1A | Q8NC01 | ENSG00000150048 | C-type lectin domain family 1, member A | Glycan Binding/lectin | Not Specific | - |
| 104 | CLEC1B | Q9P126 | ENSG00000165682 | C-type lectin domain family 1, member B | Glycan Binding/lectin | Not Specific | attachment factor for Human immunodeficiency virus type 1 (HIV-1) |
| 105 | CLEC2A | Q6UVW9 | ENSG00000188393 | C-type lectin domain family 2, member A | Glycan Binding/lectin | Not Specific | - |
| 106 | CLEC2B | Q92478 | ENSG00000110852 | C-type lectin domain family 2, member B | Glycan Binding/lectin | Sialic Acid ? | rhodocytin (aggretin), podoplanin (disialyl core ( NeuAcα2-3Galβ1-3(NeuAcα2-6)GalNAc) and the stereostructure of the protein) |
| 107 | CLEC2D | Q9UHP7 | ENSG00000069493 | C-type lectin domain family 2, member D | Glycan Binding/lectin | Not Specific | high molecular weight sulfated glycosaminoglycans |
| 108 | CLEC2L | P0C7M8 | ENSG00000236279 | C-type lectin domain family 2, member L | Glycan Binding/lectin | Not Specific | - |
| 109 | CLEC3A | O75596 | ENSG00000166509 | C-type lectin domain family 3, member A | Glycan Binding/lectin | Not Specific | Laminin-332 and fibronectin. |
| 110 | CLEC3B | P05452 | ENSG00000163815 | C-type lectin domain family 3, member B | Glycan Binding/lectin | Not Specific | plasminogen |
| 111 | CLEC4A | Q9UMR7 | ENSG00000111729 | C-type lectin domain family 4, member A | Glycan Binding/lectin | More than one monosacchrides | Mannose, Fucose, N-acetylglucosamine (GlcNAc) |
| 112 | CLEC4C | Q8WTT0 | ENSG00000198178 | C-type lectin domain family 4, member C | Glycan Binding/lectin | Galactose | Non-sialylated galactose-terminated biantennary glycans containing the trisaccharide epitope Gal(beta1-3/4)GlcNAc(beta1-2)Man , serum IgG |
| 113 | CLEC4D | Q8WXI8 | ENSG00000166527 | C-type lectin domain family 4, member D | Glycan Binding/lectin | Mannose | Alpha-mannans on C.albicans hypheas and mycobacterial trehalose 6,6'-dimycolate (TDM) |
| 114 | CLEC4E | Q9ULY5 | ENSG00000166523 | C-type lectin domain family 4, member E | Glycan Binding/lectin | Non-Human glycans | Mycobacterial trehalose 6,6'-dimycolate (TDM), damage-associated molecular patterns (DAMPs) |
| 115 | CLEC4F | Q8N1N0 | ENSG00000152672 | C-type lectin domain family 4, member F | Glycan Binding/lectin | More than one monosacchrides | Galactose and fucose |
| 116 | CLEC4G | Q6UXB4 | ENSG00000182566 | C-type lectin domain family 4, member G | Glycan Binding/lectin | More than one monosacchrides | Mannose, N-acetylglucosamine (GlcNAc) and fucose |
| 117 | CLEC4M | Q9H2X3 | ENSG00000104938 | C-type lectin domain family 4, member M | Glycan Binding/lectin | Mannose | Mannose-like carbohydrates, Man9GlcNAc2 |
| 118 | CLEC5A | Q9NY25 | ENSG00000258227 | C-type lectin domain family 5, member A | Glycan Binding/lectin | Not Specific | Self Ligand , Glycan ligand not clear, Galectin-9? |
| 119 | CLEC6A | Q6EIG7 | ENSG00000205846 | C-type lectin domain family 6, member A | Glycan Binding/lectin | Mannose | Alpha-mannans, mannose-dependent manner, high-mannose structures such as Man9GlcNAc2 |
| 120 | CLEC7A | Q9BXN2 | ENSG00000172243 | C-type lectin domain family 7, member A | Glycan Binding/lectin | Not Specific | Beta-1,3-linked and beta-1,6-linked glucans |
| 121 | CLEC9A | Q6UXN8 | ENSG00000197992 | C-type lectin domain family 9, member A | Glycan Binding/lectin | Not Specific | Filamentous actin (F-actin) |
| 122 | CLGN | O14967 | ENSG00000153132 | Calmegin | chaperone | Not Specific | chaperone |
| 123 | CMAHP | Q9Y471 | ENSG00000168405 | Inactive cytidine monophosphate-N-acetylneuraminic acid hydroxylase | Inactive/Putative | Inactive/Putative | Hydroxylase |
| 124 | CMAS | Q8NFW8 | ENSG00000111726 | N-acylneuraminate cytidylyltransferase | Transferase | Sialic Acid | Cytidine, Sialic acid synthesis |
| 125 | COG1 | Q8WTW3 | ENSG00000166685 | Component of oligomeric golgi complex 1 | Transport | Protein transport | |
| 126 | COG2 | Q14746 | ENSG00000135775 | Component of oligomeric golgi complex 2 | Transport | Protein transport | |
| 127 | COG3 | Q96JB2 | ENSG00000136152 | Component of oligomeric golgi complex 3 | Transport | Protein transport | |
| 128 | COG4 | Q9H9E3 | ENSG00000103051 | Component of oligomeric golgi complex 4 | Transport | Protein transport | |
| 129 | COG5 | Q9UP83 | ENSG00000164597 | Component of oligomeric golgi complex 5 | Transport | Protein transport | |
| 130 | COG6 | Q9Y2V7 | ENSG00000133103 | Component of oligomeric golgi complex 6 | Transport | Protein transport | |
| 131 | COG7 | P83436 | ENSG00000168434 | Component of oligomeric golgi complex 7 | Transport | Protein transport | |
| 132 | COG8 | Q96MW5 | ENSG00000213380 | Component of oligomeric golgi complex 8 | Transport | Protein transport | |
| 133 | COLEC10 | Q9Y6Z7 | ENSG00000184374 | Collectin sub-family member 10 (C-type lectin) | Glycan Binding/lectin | More than one monosacchrides | galactose > mannose = fucose > N-acetylglucosamine > N-acetylgalactosamine |
| 134 | COLEC11 | Q9BWP8 | ENSG00000118004 | Collectin sub-family member 11 | Glycan Binding/lectin | More than one monosacchrides | galactose > mannose = fucose > N-acetylglucosamine > N-acetylgalactosamine |
| 135 | COLEC12 | Q5KU26 | ENSG00000158270 | Collectin sub-family member 12 | Glycan Binding/lectin | More than one monosacchrides | Gal-type ligands, D-galactose, L- and D-fucose, GalNAc, T and Tn antigens |
| 136 | COLGALT1 | Q8NBJ5 | ENSG00000130309 | Procollagen galactosyltransferase 1 | Transferase | Inactive/Putative | |
| 137 | COLGALT2 | Q8IYK4 | ENSG00000198756 | Procollagen galactosyltransferase 2 | Transferase | Inactive/Putative | |
| 138 | CSGlcA-T | Q9P2E5 | ENSG00000033100 | Chondroitin sulfate glucuronyltransferase | Transferase | More than one monosacchrides | glucuronic acid (GlcUA)/Inactive |
| 139 | CRPPA | A4D126 | ENSG00000214960 | D-ribitol-5-phosphate cytidylyltransferase | Transferase | Mannose | CDP on L-ribitol |

| | | | | | | | |
|---|---|---|---|---|---|---|---|
| 140 | CSGALNACT1 | Q8TDX6 | ENSG00000147408 | Chondroitin sulfate N-acetylgalactosaminyltransferase 1 | Transferase | N-acetylgalactosamine (GalNAc) | chondroitin polymerization |
| 141 | CSGALNACT2 | Q8N6G5 | ENSG00000169826 | Chondroitin sulfate N-acetylgalactosaminyltransferase 2 | Transferase | N-acetylgalactosamine (GalNAc) | chondroitin polymerization |
| 142 | CTBS | Q01459 | ENSG00000117151 | Di-N-acetylchitobiase | Glycosidase | N-acetyl-beta-D-glucosamine | N-acetyl-beta-D-glucosamine (1-4)N-acetylglucosamine chitobiose core |
| 143 | DAD1 | P61803 | ENSG00000129562 | Defender against cell death 1 | Transferase | Mannose | Glc3Man9GlcNAc2 |
| 144 | DCAL1/CLECL1P | Q8IZS7 | ENSG00000184293 | Putative C-type lectin-like domain family 1 | Inactive/Putative | Inactive/Putative | |
| 145 | DDOST | P39656 | ENSG00000244038 | Dolichyl-diphosphooligosaccharide-protein glycosyltransferase | Transferase | More than one monosacchrides | Glc3Man9GlcNAc2 |
| 146 | DHDDS | Q86SQ9 | ENSG00000117682 | Dehydrodolichyl diphosphate synthase | Transferase | Phosphate | dolichol phosphate synthesis |
| 147 | DOLPP1 | Q86YN1 | ENSG00000167130 | Dolichyl pyrophosphate phosphatase 1 | Miscellaneous | Phosphate | dolichyl pyrophosphate |
| 148 | DNAJB11 | Q9UBS4 | ENSG00000090520 | DnaJ homolog subfamily B member 11 | chaperone | Not Specific | Glycosylation Regulation |
| 149 | DNAJC10 | Q8IXB1 | ENSG00000077232 | DnaJ homolog subfamily C member 10 | chaperone | Not Specific | Glycosylation Regulation |
| 150 | DPAGT1 | Q9H3H5 | ENSG00000172269 | Dolichyl-phosphate (UDP-N-acetylglucosamine) N-acetylglucosaminephosphotransferase 1 (GlcNAc-1-P transferase) | Transferase | N-acetylglucosamine (GlcNAc) | N-acetyl-alpha-D-glucosamine |
| 151 | DPM1 | O60762 | ENSG00000000419 | Dolichyl-phosphate mannosyltransferase polypeptide 1, catalytic subunit | Transferase | Mannose | mannose |
| 152 | DPM2 | O94777 | ENSG00000136908 | Dolichyl-phosphate mannosyltransferase polypeptide 2, regulatory subunit | Regulator | Not Specific | Regulatory subunit of the dolichol-phosphate mannose (DPM) synthase complex |
| 153 | DPM3 | Q9P2X0 | ENSG00000179085 | Dolichyl-phosphate mannosyltransferase polypeptide 3 | Regulator | Not Specific | Stabilizer subunit |
| 154 | DPY19L1 | Q2PZI1 | ENSG00000173852 | Probable C-mannosyltransferase DPY19L1 | Transferase | Mannose | C-Mannose |
| 155 | DPY19L2 | Q6NUT2 | ENSG00000177990 | Probable C-mannosyltransferase DPY19L2 | Transferase | Mannose | C-Mannose |
| 156 | DPY19L3 | Q6ZPD9 | ENSG00000178904 | Probable C-mannosyltransferase DPY19L3 | Transferase | Mannose | C-Mannose |
| 157 | DPY19L4 | Q7Z388 | ENSG00000156162 | Probable C-mannosyltransferase DPY19L4 | Transferase | Mannose | C-Mannose |
| 158 | DSE | Q9UL01 | ENSG00000111817 | Dermatan sulfate epimerase | Epimerase | Glucuronic acid (GlcA) | Converts D-glucuronic acid to L-iduronic acid (IdoUA) |
| 159 | DSEL | Q8IZU8 | ENSG00000171451 | Dermatan-sulfate epimerase-like protein | Epimerase | Glucuronic acid (GlcA) | D-glucuronic acid to L-iduronic acid (IdoUA) |
| 160 | EDEM1 | Q92611 | ENSG00000134109 | ER degradation enhancer, mannosidase alpha-like 1 | Glycosidase | Mannose | misfolded glycoproteins/Mannose |
| 161 | EDEM2 | Q9BV94 | ENSG00000088298 | ER degradation enhancer, mannosidase alpha-like 2 | Glycosidase | Mannose | misfolded glycoproteins/Mannose |
| 162 | EDEM3 | Q9BZQ6 | ENSG00000116406 | ER degradation enhancer, mannosidase alpha-like 3 | Glycosidase | Mannose | misfolded glycoproteins/Mannose |
| 163 | ENGASE | Q8NFI3 | ENSG00000167280 | Cytosolic endo-beta-N-acetylglucosaminidase | Glycosidase | N-acetylglucosamine (GlcNAc) | N-glycans |
| 164 | EOGT | Q5NDL2 | ENSG00000163378 | EGF domain-specific O-linked N-acetylglucosamine transferase | Transferase | N-acetylglucosamine (GlcNAc) | N-acetylglucosamine |
| 165 | ERLEC1 | Q96DZ1 | ENSG00000068912 | Endoplasmic reticulum lectin 1 | Regulator | Not Specific | improperly folded lumenal proteins |
| 166 | EXT1 | Q16394 | ENSG00000182197 | Exostoses (multiple) 1 | Transferase | N-acetylglucosamine (GlcNAc) | heparan-sulfate |
| 167 | EXT2 | Q93063 | ENSG00000151348 | Exostoses (multiple) 2 | Transferase | N-acetylglucosamine (GlcNAc) | heparan-sulfate |
| 168 | EXTL1 | Q92935 | ENSG00000158008 | Exostoses (multiple)-like 1 | Transferase | N-acetylglucosamine (GlcNAc) | heparan-sulfate |
| 169 | EXTL2 | Q9UBQ6 | ENSG00000162694 | Exostoses (multiple)-like 2 | Transferase | N-acetylglucosamine (GlcNAc) | heparan-sulfate |
| 170 | EXTL3 | O43909 | ENSG00000012232 | Exostoses (multiple)-like 3 | Transferase | N-acetylglucosamine (GlcNAc) | heparan-sulfate |
| 171 | FAM20B | O75063 | ENSG00000116199 | Glycosaminoglycan xylosylkinase | Kinase | Phosphate | 2-O-phosphorylation of xylose |
| 172 | FBXO27 | Q8NI29 | ENSG00000161243 | F-box protein 27 | Ubiquitin ligase | Not Specific | denatured glycoproteins |
| 173 | FBXO44 | Q9H4M3 | ENSG00000132879 | F-box protein 44 | Ubiquitin ligase | Not Specific | no significant binding |
| 174 | FBXO6 | Q9NRD1 | ENSG00000116663 | F-box protein 6 | Ubiquitin ligase | Not Specific | unfolded glycoproteins |
| 175 | FCMD | O75072 | ENSG00000106692 | Fukutin/ Ribitol-5-phosphate transferase | Transferase | Ribitol | ribitol-phosphate |
| 176 | FCSK | Q8N0W3 | ENSG00000157353 | L-fucose kinase | Kinase | Phosphate | Phosphate on fucose |
| 177 | FCN1 | O00602 | ENSG00000085265 | Ficolin (collagen/fibrinogen domain containing) 1 | Glycan Binding/lectin | Not Specific | pathogen-associated molecular patterns (PAMPs) |
| 178 | FCN2 | Q15485 | ENSG00000160339 | Ficolin (collagen/fibrinogen domain containing lectin) 2 (hucolin) | Glycan Binding/lectin | N-acetylglucosamine (GlcNAc) | |
| 179 | FCN3 | O75636 | ENSG00000142748 | Ficolin (collagen/fibrinogen domain containing) 3 (Hakata antigen) | Glycan Binding/lectin | More than one monosacchrides | GlcNAc, also have affinity for GalNAc, GlcNAc, D-fucose, as mono/oligosaccharide |
| 180 | FKRP | Q9H9S5 | ENSG00000181027 | Fukutin related protein | Transferase | Mannose | ribitol 5-phosphate |
| 181 | FKTN | O75072 | ENSG00000106692 | Fukutin | Transferase | Mannose | ribitol-phosphate |
| 182 | FLJ21865 | Q8NFI3 | ENSG00000167280 | Endo-beta-N-acetylglucosaminidase | Glycosidase | Not Specific | N-glycans |
| 183 | FPGT | O14772 | ENSG00000254685 | Fucose-1-phosphate guanylyltransferase | Transferase | Fucose | GDP on L-fucose |
| 184 | FUCA1 | P04066 | ENSG00000179163 | Fucosidase, alpha-L- 1, tissue | Glycosidase | Fucose | N-linked_Lysosome_Degradation,O-linked_Lysosome_Degradation,GSL_Lysosome_Degradation |
| 185 | FUCA2 | Q9BTY2 | ENSG00000001036 | Fucosidase, alpha-L- 2, plasma | Glycosidase | Fucose | N-linked_Lysosome_Degradation,O-linked_Lysosome_Degradation,GSL_Lysosome_Degradation |
| 186 | FUT1 | P19526 | ENSG00000174951 | Fucosyltransferase 1 (galactoside 2-alpha-L-fucosyltransferase, H blood group) | Transferase | Fucose | Capping,ABO_BloodGroups,Fucosylation |
| 187 | FUT10 | Q6P4F1 | ENSG00000172728 | Fucosyltransferase 10 (alpha (1,3) fucosyltransferase) | Transferase | Fucose | Capping,Fucosylation |
| 188 | FUT11 | Q495W5 | ENSG00000196968 | Fucosyltransferase 11 (alpha (1,3) fucosyltransferase) | Transferase | Fucose | Capping,Fucosylation |
| 189 | FUT2 | Q10981 | ENSG00000176920 | Fucosyltransferase 2 (secretor status included) | Transferase | Fucose | Capping,ABO_BloodGroups,Fucosylation |
| 190 | FUT3 | P21217 | ENSG00000171124 | Fucosyltransferase 3 (galactoside 3(4)-L-fucosyltransferase, Lewis blood group) | Transferase | Fucose | Capping,Fucosylation |
| 191 | FUT4 | P22083 | ENSG00000196371 | Fucosyltransferase 4 (alpha (1,3) fucosyltransferase, myeloid-specific) | Transferase | Fucose | Capping,Fucosylation |
| 192 | FUT5 | Q11128 | ENSG00000130383 | Fucosyltransferase 5 (alpha (1,3) fucosyltransferase) | Transferase | Fucose | Capping,Fucosylation |
| 193 | FUT6 | P51993 | ENSG00000156413 | Fucosyltransferase 6 (alpha (1,3) fucosyltransferase) | Transferase | Fucose | Capping,Fucosylation |
| 194 | FUT7 | Q11130 | ENSG00000180549 | Fucosyltransferase 7 (alpha (1,3) fucosyltransferase) | Transferase | Fucose | Capping,Fucosylation |
| 195 | FUT8 | Q9BYC5 | ENSG00000033170 | Fucosyltransferase 8 (alpha (1,6) fucosyltransferase) | Transferase | Fucose | N-linked,Nlinked_Branching |
| 196 | FUT9 | Q9Y231 | ENSG00000172461 | Fucosyltransferase 9 (alpha (1,3) fucosyltransferase) | Transferase | Fucose | Capping,Fucosylation |
| 197 | G6PC1 | P35575 | ENSG00000131482 | Glucose-6-phosphatase catalytic subunit 1 | Phosphatase | Phosphate | remove phosphate glucose-6-phosphate |
| 198 | GAA | P10253 | ENSG00000171298 | Glucosidase, alpha; acid | Glycosidase | Glucose | Glucosidase/Glycogen breakdown |
| 199 | GAL3ST1 | Q99999 | ENSG00000128242 | Galactose-3-O-sulfotransferase 1 | Transferase | Sulfate | GalCer,GlcCer,Terminal_Gal_Sulfation |
| 200 | GAL3ST2 | Q9H3Q3 | ENSG00000154252 | Galactose-3-O-sulfotransferase 2 | Transferase | Sulfate | Glycolipids,GlcCer,O-GalNAc,Terminal_Gal_Sulfation |
| 201 | GAL3ST3 | Q96A11 | ENSG00000175229 | Galactose-3-O-sulfotransferase 3 | Transferase | Sulfate | Glycolipids,GlcCer,O-GalNAc,Terminal_Gal_Sulfation |
| 202 | GAL3ST4 | Q96RP7 | ENSG00000197093 | Galactose-3-O-sulfotransferase 4 | Transferase | Sulfate | Glycolipids,GlcCer,O-GalNAc,Terminal_Gal_Sulfation |
| 203 | GALC | P54803 | ENSG00000054983 | Galactosylceramidase | Glycosidase | Galactose | GSL_Lysosome_Degradation |
| 204 | GALE | Q14376 | ENSG00000117308 | UDP-glucose 4-epimerase | Epimerase | Galactose | epimerization of UDP-glucose to UDP-galactose and UDP-N-acetylglucosamine to UDP-N-acetylgalactosamine |
| 205 | GALK1 | P51570 | ENSG00000108479 | Galactokinase | Kinase | Phosphate | Phosphate on alpha-D-galactose |
| 206 | GALK2 | Q01415 | ENSG00000156958 | N-acetylgalactosamine kinase | Kinase | Phosphate | Phosphate on N-acetyl-alpha-D-galactosamine |
| 207 | GALNAC4S-6ST | Q7LFX5 | ENSG00000182022 | Carbohydrate (N-acetylgalactosamine 4-sulfate 6-O) sulfotransferase 15 | Transferase | Sulfate | |
| 208 | GALNS | P34059 | ENSG00000141012 | Galactosamine (N-acetyl)-6-sulfate sulfatase | Sulfatase | Sulfate | CS_Lysosome_Degradation,DS_Lysosome_Degradation,KS_Lysosome_Degradation |
| 209 | GALNT1 | Q10472 | ENSG00000141429 | UDP-N-acetyl-alpha-D-galactosamine:polypeptide N-acetylgalactosaminyltransferase 1 (GalNAc-T1) | Transferase | N-acetylgalactosamine (GalNAc) | O-GalNAc |
| 210 | GALNT10 | Q86SR1 | ENSG00000164574 | UDP-N-acetyl-alpha-D-galactosamine:polypeptide N-acetylgalactosaminyltransferase 10 (GalNAc-T10) | Transferase | N-acetylgalactosamine (GalNAc) | O-GalNAc |
| 211 | GALNT11 | Q8NCW6 | ENSG00000178234 | UDP-N-acetyl-alpha-D-galactosamine:polypeptide N-acetylgalactosaminyltransferase 11 (GalNAc-T11) | Transferase | N-acetylgalactosamine (GalNAc) | O-GalNAc |
| 212 | GALNT12 | Q8IXK2 | ENSG00000119514 | UDP-N-acetyl-alpha-D-galactosamine:polypeptide N-acetylgalactosaminyltransferase 12 (GalNAc-T12) | Transferase | N-acetylgalactosamine (GalNAc) | O-GalNAc |
| 213 | GALNT13 | Q8IUC8 | ENSG00000144278 | UDP-N-acetyl-alpha-D-galactosamine:polypeptide N-acetylgalactosaminyltransferase 13 (GalNAc-T13) | Transferase | N-acetylgalactosamine (GalNAc) | O-GalNAc |
| 214 | GALNT14 | Q96FL9 | ENSG00000158089 | UDP-N-acetyl-alpha-D-galactosamine:polypeptide N-acetylgalactosaminyltransferase 14 (GalNAc-T14) | Transferase | N-acetylgalactosamine (GalNAc) | O-GalNAc |
| 215 | GALNT15 | Q7Z4T8 | ENSG00000106648 | Inactive polypeptide N-acetylgalactosaminyltransferase-like protein 5 | Inactive/Putative | Inactive/Putative | O-GalNAc |
| 216 | GALNT16 | Q8N428 | ENSG00000100626 | Polypeptide N-acetylgalactosaminyltransferase 16 | Transferase | N-acetylgalactosamine (GalNAc) | O-GalNAc |
| 217 | GALNT17 | Q49A17 | ENSG00000174473 | UDP-N-acetyl-alpha-D-galactosamine:polypeptide N-acetylgalactosaminyltransferase-like 6 | Transferase | N-acetylgalactosamine (GalNAc) | O-GalNAc |
| 218 | GALNT18 | Q6P9A2 | ENSG00000110328 | Polypeptide N-acetylgalactosaminyltransferase 18 | Transferase | N-acetylgalactosamine (GalNAc) | O-GalNAc |
| 219 | GALNT19 | Q2L4S5 | ENSG00000185274 | Polypeptide N-acetylgalactosaminyltransferase 18 | Transferase | N-acetylgalactosamine (GalNAc) | O-GalNAc |
| 220 | GALNT2 | Q10471 | ENSG00000143641 | UDP-N-acetyl-alpha-D-galactosamine:polypeptide N-acetylgalactosaminyltransferase 2 (GalNAc-T2) | Transferase | N-acetylgalactosamine (GalNAc) | O-GalNAc |
| 221 | GALNT3 | Q14435 | ENSG00000115339 | UDP-N-acetyl-alpha-D-galactosamine:polypeptide N-acetylgalactosaminyltransferase 3 (GalNAc-T3) | Transferase | N-acetylgalactosamine (GalNAc) | O-GalNAc |
| 222 | GALNT4 | Q8N4A0 | ENSG00000257594 | UDP-N-acetyl-alpha-D-galactosamine:polypeptide N-acetylgalactosaminyltransferase 4 (GalNAc-T4) | Transferase | N-acetylgalactosamine (GalNAc) | O-GalNAc |
| 223 | GALNT5 | Q7Z7M9 | ENSG00000136542 | UDP-N-acetyl-alpha-D-galactosamine:polypeptide N-acetylgalactosaminyltransferase 5 (GalNAc-T5) | Transferase | N-acetylgalactosamine (GalNAc) | O-GalNAc |
| 224 | GALNT6 | Q8NCL4 | ENSG00000139629 | UDP-N-acetyl-alpha-D-galactosamine:polypeptide N-acetylgalactosaminyltransferase 6 (GalNAc-T6) | Transferase | N-acetylgalactosamine (GalNAc) | O-GalNAc |
| 225 | GALNT7 | Q86SF2 | ENSG00000109586 | UDP-N-acetyl-alpha-D-galactosamine:polypeptide N-acetylgalactosaminyltransferase 7 (GalNAc-T7) | Transferase | N-acetylgalactosamine (GalNAc) | O-GalNAc |
| 226 | GALNT8 | Q9NY28 | ENSG00000130035 | UDP-N-acetyl-alpha-D-galactosamine:polypeptide N-acetylgalactosaminyltransferase 8 (GalNAc-T8) | Transferase | N-acetylgalactosamine (GalNAc) | O-GalNAc |
| 227 | GALNT9 | Q9HCQ5 | ENSG00000182870 | UDP-N-acetyl-alpha-D-galactosamine:polypeptide N-acetylgalactosaminyltransferase 9 (GalNAc-T9) | Transferase | N-acetylgalactosamine (GalNAc) | O-GalNAc |
| 228 | GALNTL1/GALNT16 | Q8N428 | ENSG00000100626 | UDP-N-acetyl-alpha-D-galactosamine:polypeptide N-acetylgalactosaminyltransferase-like 1 | Transferase | N-acetylgalactosamine (GalNAc) | Initiation of O-linked glycan |
| 229 | GALNTL2/GALNT15 | Q8N3T1 | ENSG00000131386 | UDP-N-acetyl-alpha-D-galactosamine:polypeptide N-acetylgalactosaminyltransferase-like 2 | Transferase | N-acetylgalactosamine (GalNAc) | Initiation of O-linked glycan |
| 230 | GALNTL4/GALNT18 | Q6P9A2 | ENSG00000110328 | UDP-N-acetyl-alpha-D-galactosamine:polypeptide N-acetylgalactosaminyltransferase-like 4 | Transferase | N-acetylgalactosamine (GalNAc) | Initiation of O-linked glycan |
| 231 | GALNTL5 | Q7Z4T8 | ENSG00000106648 | Inactive polypeptide N-acetylgalactosaminyltransferase-like protein 5 | Inactive/Putative | Inactive/Putative | Probable inactive glycosyltransferase required during spermatid development |
| 232 | GALT | P07902 | ENSG00000213930 | Galactose-1-phosphate uridylyltransferase | Transferase | Galactose | UDP |
| 233 | GANAB | Q14697 | ENSG00000089597 | Glucosidase, alpha; neutral AB | Glycosidase | Glucose | N-linked |
| 234 | GANC | Q8TET4 | ENSG00000214013 | glucosidase, alpha; neutral C | Glycosidase | Glucose | Glycogen_Breakdown |
| 235 | GBA | P04062 | ENSG00000177628 | Glucosidase, beta; acid (includes glucosylceramidase) | Glycosidase | Glucose | GSL_Glycan_Degradation |
| 236 | GBA2 | Q9HCG7 | ENSG00000070610 | Glucosidase, beta (bile acid) 2 | Glycosidase | Glucose | GSL_Glycan_Degradation |
| 237 | GBA3 | Q9H227 | ENSG00000249948 | Glucosidase, beta, acid 3 (cytosolic) | Glycosidase | Glucose | GSL_Glycan_Degradation |
| 238 | GBAP | - | ENSG00000160766 | Glucosidase, beta/ glucosylceramidase beta pseudogene 1 | Inactive/Putative | Inactive/Putative | |
| 239 | GBGT1 | Q8N5D6 | ENSG00000148288 | Globoside alpha-1,3-N-acetylgalactosaminyltransferase 1 | Inactive/Putative | Inactive/Putative | |
| 240 | GCK | P35557 | ENSG00000106633 | Glucokinase | Kinase | Phosphate | Nucleotide_Sugar_Synthesis,UDP-Glc_Synthesis,GDP-Man_Synthesis |
| 241 | GCNT1 | Q02742 | ENSG00000187210 | Glucosaminyl (N-acetyl) transferase 1, core 2 (beta-1,6-N-acetylglucosaminyltransferase) | Transferase | N-acetylglucosamine (GlcNAc) | O-GalNAc, Core2 |
| 242 | GCNT2 | Q8N0V5 | ENSG00000111846 | Glucosaminyl (N-acetyl) transferase 2, I-branching enzyme (I blood group) | Transferase | N-acetylglucosamine (GlcNAc) | I-Branching |
| 243 | GCNT3 | O95395 | ENSG00000140297 | Glucosaminyl (N-acetyl) transferase 3, mucin type | Transferase | N-acetylglucosamine (GlcNAc) | O-GalNAc, Core4,O-GalNAc,Core2,Elongation_and_Branching, I-Branching |
| 244 | GCNT4 | Q9P109 | ENSG00000176928 | Glucosaminyl (N-acetyl) transferase 4, core 2 (beta-1,6-N-acetylglucosaminyltransferase) | Transferase | N-acetylglucosamine (GlcNAc) | O-GalNAc, Core2 |
| 245 | GCNT6/GCNT2P1 | - | ENSG00000205318 | Glucosaminyl (N-acetyl) transferase 6/GCNT2 Pseudogene 1 | Inactive/Putative | Inactive/Putative | |
| 246 | GCNT7 | Q6ZNI0 | ENSG00000124091 | Glucosaminyl (N-acetyl) transferase family member 7 | Transferase | N-acetylglucosamine (GlcNAc) | Predicted |
| 247 | GCS1 | Q13724 | ENSG00000115275 | Mannosyl-oligosaccharide glucosidase | Glycosidase | Glucose | Glycan metabolism; N-glycan degradation |
| 248 | GFPT1 | Q06210 | ENSG00000198380 | Glutamine-fructose-6-phosphate aminotransferase [isomerizing] 1 | Transferase | N-acetylglucosamine (GlcNAc) | Amine on fructose 6-phosphate to synthesize D-glucosamine 6-phosphate |
| 249 | GFPT2 | O94808 | ENSG00000131459 | Glutamine-fructose-6-phosphate aminotransferase [isomerizing] 2 | Transferase | N-acetylglucosamine (GlcNAc) | Amine on fructose 6-phosphate to synthesize D-glucosamine 6-phosphate |
| 250 | GFUS | Q13630 | ENSG00000104522 | GDP-L-fucose synthase | Transferase | Fucose | Conversion of GDP-4-dehydro-6-deoxy-D-mannose to GDP-fucose |
| 251 | GGTA1 | Q4G0N0 | ENSG00000204136 | Glycoprotein, alpha-galactosyltransferase 1 | Inactive/Putative | Inactive/Putative | |
| 252 | GLA | P06280 | ENSG00000102393 | Galactosidase, alpha | Glycosidase | Galactose | GSL_Lysosome_Degradation |
| 253 | GLB1 | P16278 | ENSG00000170266 | Galactosidase, beta 1 | Glycosidase | Galactose | N-linked_Lysosome_Degradation,O-linked_Lysosome_Degradation,GSL_Lysosome_Degradation,KS_Lysosome_Degradation |
| 254 | GLB1L | Q6UWU2 | ENSG00000163521 | Galactosidase, beta 1-like | Inactive/Putative | Inactive/Putative | |
| 255 | GLCE | O94923 | ENSG00000138604 | Glucuronic acid epimerase | Epimerase | Glucuronic acid (GlcA) | D-glucuronic acid to L-iduronic acid |
| 256 | GM2A | P17900 | ENSG00000196743 | GM2 ganglioside activator | Miscellaneous | Glycolipid | breakdown of ganglioside GM2 and glycolipid GA2 |
| 257 | GMDS | O60547 | ENSG00000112699 | GDP-mannose 4,6 dehydratase | Miscellaneous | Mannose | GDP-D-mannose to GDP-4-dehydro-6-deoxy-D-mannose |
| 258 | GMPPA | Q96IJ6 | ENSG00000144591 | Mannose-1-phosphate guanyltransferase alpha | Transferase | Mannose | GDP to mannose |
| 259 | GMPPB | Q9Y5P6 | ENSG00000111670 | Mannose-1-phosphate guanyltransferase beta | Transferase | Mannose | GDP to mannose |
| 260 | GNE | Q9Y223 | ENSG00000159921 | Bifunctional UDP-N-acetylglucosamine 2-epimerase/N-acetylmannosamine kinase | Epimerase | Sialic Acid | Initiates biosynthesis of sialic acid (convert glucosamine to mannosamine) |
| 261 | GNPDA1 | P46926 | ENSG00000113552 | Glucosamine-6-phosphate isomerase 1 | Isomerase | N-acetylglucosamine (GlcNAc) | alpha-D-glucosamine 6-phosphate (GlcN-6P) into beta-D-fructose 6-phosphate (Fru-6P) |
| 262 | GNPDA2 | Q8TDQ7 | ENSG00000163281 | Glucosamine-6-phosphate isomerase 2 | Isomerase | N-acetylglucosamine (GlcNAc) | alpha-D-glucosamine 6-phosphate (GlcN-6P) into beta-D-fructose 6-phosphate (Fru-6P) |
| 263 | GNPNAT1 | Q96EK6 | ENSG00000100522 | Glucosamine 6-phosphate N-acetyltransferase | Transferase | Acetate | |
| 264 | GNPTAB | Q3T906 | ENSG00000111670 | N-acetylglucosamine-1-phosphate transferase, alpha and beta subunits | Transferase | Mannose | formation of mannose 6-phosphate |
| 265 | GNPTG | Q9UJJ9 | ENSG00000090581 | N-acetylglucosamine-1-phosphate transferase, gamma subunit | Regulator | Mannose | formation of mannose 6-phosphate |
| 266 | GNS | P15586 | ENSG00000135677 | Glucosamine (N-acetyl)-6-sulfatase | Sulphatase | Sulfate | Sulphate from N-acetyl-D-glucosamine 6-sulfate |
| 267 | GPAA1 | O43292 | ENSG00000197858 | Glycosylphosphatidylinositol anchor attachment protein 1 homolog (yeast) | Transamidase | Glycolipid | Transfer of GPI to proteins |
| 268 | GPI | P06744 | ENSG00000105220 | Glucose-6-phosphate isomerase | Isomerase | Glucose | Conversion of glucose-6-phosphate to fructose-6-phosphate |
| 269 | GPLD1 | P80108 | ENSG00000112293 | Glycosylphosphatidylinositol specific phospholipase D1 | Miscellaneous | Phosphate | remove phosphate from GPI-anchored protein |
| 270 | GUSB | P08236 | ENSG00000169919 | Glucuronidase, beta | Glycosidase | Glucuronic acid (GlcA) | Beta-D-glucuronoside |
| 271 | GXYLT1 | Q4G148 | ENSG00000151233 | Glucoside xylosyltransferase 1 | Transferase | Xylose | O-Glc in EGF |
| 272 | GXYLT2 | A0PJZ3 | ENSG00000172986 | Glucoside xylosyltransferase 2 | Transferase | Xylose | O-Glc in EGF |
| 273 | GYLTL1B | Q8N3Y3 | ENSG00000165905 | Glycosyltransferase-like 1B/LARGE Xylosyl- And Glucuronyltransferase 2 | Transferase | More than one monosacchrides | Xylose, glucuronate |
| 274 | HAS1 | Q92839 | ENSG00000105509 | Hyaluronan synthase 1 | Transferase | More than one monosacchrides | glucosamine, glucuronate |
| 275 | HAS2 | Q92819 | ENSG00000170961 | Hyaluronan synthase 2 | Transferase | More than one monosacchrides | glucosamine, glucuronate |
| 276 | HAS3 | O00219 | ENSG00000103044 | Hyaluronan synthase 3 | Transferase | More than one monosacchrides | glucosamine, glucuronate |
| 277 | HEXA | P06865 | ENSG00000213614 | Hexosaminidase A (alpha polypeptide) | Glycosidase | More than one monosacchrides | Hydrolyzes the non-reducing end N-acetyl-D-hexosamine and/or sulfated N-acetyl-D-hexosamine |
| 278 | HEXB | P07686 | ENSG00000049860 | Hexosaminidase B (beta polypeptide) | Glycosidase | More than one monosacchrides | Hydrolyzes the non-reducing end N-acetyl-D-hexosamine and/or sulfated N-acetyl-D-hexosamine |
| 279 | HEXDC | Q8WVB3 | ENSG00000169660 | Hexosaminidase (glycosyl hydrolase family 20, catalytic domain) containing | Glycosidase | More than one monosacchrides | cleavage of the monosaccharides N-acetylglucosamine (GlcNAc) and N-acetylgalactosamine (GalNAc) |
| 280 | HGSNAT | Q68CP4 | ENSG00000165102 | Heparan-alpha-glucosaminide N-acetyltransferase | Transferase | Acetate | Acetate on alpha-D-glucosaminyl-[heparan sulfate] |

| | | | | | | | |
|---|---|---|---|---|---|---|---|
| 281 | HK1 | P19367 | ENSG00000156515 | Hexokinase-1 | Kinase | Phosphate | phosphorylation of D-glucose, D-glucosamine, D-fructose, D-mannose and 2-deoxy-D-glucose |
| 282 | HK2 | P52789 | ENSG00000159399 | Hexokinase-2 | Kinase | Phosphate | phosphorylation of D-glucose, D-fructose |
| 283 | HK3 | P52790 | ENSG00000160883 | Hexokinase-3 | Kinase | Phosphate | phosphorylation of D-glucose, D-fructose |
| 284 | HPSE | Q9Y251 | ENSG00000173083 | Heparanase | Glycosidase | Glucuronic acid (GlcA) | Cleaves heparan sulfate proteoglycans |
| 285 | HPSE2 | Q8WWQ2 | ENSG00000172987 | Heparanase 2 | Inactive/Putative | Inactive/Putative | Glucuronidase |
| 286 | HS2ST1 | Q7LGA3 | ENSG00000153936 | Heparan sulfate 2-O-sulfotransferase 1 | Sulfotransferase | Sulfate | Heparan Sulphate Sulfation |
| 287 | HS3ST1 | O14792 | ENSG00000002587 | Heparan sulfate (glucosamine) 3-O-sulfotransferase 1 | Sulfotransferase | Sulfate | Heparan Sulphate Sulfation |
| 288 | HS3ST2 | Q9Y278 | ENSG00000122254 | Heparan sulfate (glucosamine) 3-O-sulfotransferase 2 | Sulfotransferase | Sulfate | Heparan Sulphate Sulfation |
| 289 | HS3ST3A1 | Q9Y663 | ENSG00000153976 | Heparan sulfate (glucosamine) 3-O-sulfotransferase 3A1 | Sulfotransferase | Sulfate | Heparan Sulphate Sulfation |
| 290 | HS3ST3B1 | Q9Y662 | ENSG00000125430 | Heparan sulfate (glucosamine) 3-O-sulfotransferase 3B1 | Sulfotransferase | Sulfate | Heparan Sulphate Sulfation |
| 291 | HS3ST4 | Q9Y661 | ENSG00000182601 | Heparan sulfate (glucosamine) 3-O-sulfotransferase 4 | Sulfotransferase | Sulfate | Heparan Sulphate Sulfation |
| 292 | HS3ST5 | Q8IZT8 | ENSG00000249853 | Heparan sulfate (glucosamine) 3-O-sulfotransferase 5 | Sulfotransferase | Sulfate | Heparan Sulphate Sulfation |
| 293 | HS3ST6 | Q96QI5 | ENSG00000162040 | Heparan sulfate glucosamine 3-O-sulfotransferase 6 | Sulfotransferase | Sulfate | Heparan Sulphate Sulfation |
| 294 | HS6ST1 | O60243 | ENSG00000136720 | Heparan sulfate 6-O-sulfotransferase 1 | Sulfotransferase | Sulfate | Heparan Sulphate Sulfation |
| 295 | HS6ST2 | Q96MM7 | ENSG00000171004 | Heparan sulfate 6-O-sulfotransferase 2 | Sulfotransferase | Sulfate | Heparan Sulphate Sulfation |
| 296 | HS6ST3 | Q8IZP7 | ENSG00000185352 | Heparan sulfate 6-O-sulfotransferase 3 | Sulfotransferase | Sulfate | Heparan Sulphate Sulfation |
| 297 | HSP90B1 | P14625 | ENSG00000166598 | Heat Shock Protein 90 Beta Family Member 1 | Regulator | Not Specific | chaperone |
| 298 | HSPA5 | P11021 | ENSG00000044574 | Heat Shock Protein Family A (Hsp70) Member 5 | Regulator | Not Specific | Endoplasmic reticulum chaperone |
| 299 | HSPC159 | Q3ZCW2 | ENSG00000119862 | Galectin-related protein | Glycan Binding/lectin | Not Specific | may not bind carbohydrates |
| 300 | HYAL1 | Q12794 | ENSG00000114378 | Hyaluronoglucosaminidase 1 | Glycosidase | Glucuronic acid (GlcA) | Hydrolysis of (1->4)-linkages between N-acetyl-beta-D-glucosamine and D-glucuronate residues in hyaluronate |
| 301 | HYAL2 | Q12891 | ENSG00000068001 | Hyaluronoglucosaminidase 2 | Glycosidase | Hyaluronic acid | Hydrolyzes high molecular weight hyaluronic acid |
| 302 | HYAL3 | O43820 | ENSG00000186792 | Hyaluronidase-3 | Glycosidase | Hyaluronic acid | Hyaluronic acid in sperm penetration |
| 303 | HYAL4 | Q2M3T9 | ENSG00000106302 | Hyaluronidase-4 | Glycosidase | Hyaluronic acid | Hyaluronan to smaller oligosaccharide fragments |
| 304 | HYOU1 | Q9Y4L1 | ENSG00000149428 | Hypoxia up-regulated protein 1 | Regulator | Not Specific | Chaperone |
| 305 | ICAM1 | P05362 | ENSG00000090339 | Intercellular adhesion molecule 1 | Glycan Binding/lectin | Not Specific | LFA-1 (integrin alpha-L/beta-2) |
| 306 | ICAM2 | P13598 | ENSG00000108622 | Intercellular adhesion molecule 2 | Glycan Binding/lectin | Not Specific | LFA-1 (integrin alpha-L/beta-2) |
| 307 | ICAM3 | P32942 | ENSG00000076662 | Intercellular adhesion molecule 3 | Glycan Binding/lectin | Not Specific | LFA-1 (integrin alpha-L/beta-2) (alpha-D/beta-2) |
| 308 | IDS | P22304 | ENSG00000010404 | Iduronate 2-sulfatase | Sulphatase | Sulfate | HS_Lysosome_Degradation,DS_Lysosome_Degradation |
| 309 | IDUA | P35475 | ENSG00000127415 | Iduronidase, alpha-L- | Glycosidase | Iduronic Acid | Hydrolysis of unsulfated alpha-L-iduronosidic linkages |
| 310 | IGF2R | P11717 | ENSG00000197081 | Cation-independent mannose-6-phosphate receptor | Glycan Binding/lectin | Mannose | mannose-6-phosphate |
| 311 | ITLN1 | Q8WWA0 | ENSG00000179914 | Intelectin 1 (galactofuranose binding) | Glycan Binding/lectin | Non-Human glycans | Microbial glycans that contain a terminal acyclic 1,2-diol, beta-linked D-galactofuranose (beta-Galf), D-phosphoglycerol-modified glycans, D-glycero-D-talo-oct-2-ulosonic acid (KO) and 3-deoxy-D-manno-oct-2-ulosonic acid (KDO) |
| 312 | ITLN2 | Q8WWU7 | ENSG00000158764 | Intelectin 2 | Glycan Binding/lectin | Not Specific | Not Specifically known |
| 313 | KIAA0350 | Q2KHT3 | ENSG00000038532 | C-type lectin domain family 16, member A | Glycan Binding/lectin | Not Specific | does not contain a C-type lectin domain/ proteasomal degradation |
| 314 | KLRA1 | O75889 | ENSG00000256667 | Killer cell lectin-like receptor subfamily A, member 1 | Glycan Binding/lectin | Not Specific | MHC class I chain-related A and B proteins |
| 315 | KLRB1 | Q12918 | ENSG00000111796 | Killer cell lectin-like receptor subfamily B, member 1 | Glycan Binding/lectin | More than one monosacchrides | terminal carbohydrate Gal-alpha(1,3)Gal epitope, N-acetyllactosamine epitope |
| 316 | KLRC1 | P26715 | ENSG00000134545 | Killer cell lectin-like receptor subfamily C, member 1 | Glycan Binding/lectin | Not Specific | MHC class I HLA-E molecules |
| 317 | KLRC2 | P26717 | ENSG00000205809 | Killer cell lectin-like receptor subfamily C, member 2 | Glycan Binding/lectin | Not Specific | Glycan ligand not clear |
| 318 | KLRC3 | Q07444 | ENSG00000205810 | Killer cell lectin-like receptor subfamily C, member 3 | Glycan Binding/lectin | Not Specific | MHC class I HLA-E molecules/Glycan ligand not clear |
| 319 | KLRC4 | O43908 | ENSG00000183542 | Killer cell lectin-like receptor subfamily C, member 4 | Glycan Binding/lectin | Not Specific | MHC class I HLA-E molecules/Glycan ligand not clear |
| 320 | KLRD1 | Q13241 | ENSG00000134539 | Killer cell lectin-like receptor subfamily D, member 1 | Glycan Binding/lectin | Not Specific | MHC class I HLA-E molecules/Glycan ligand not clear |
| 321 | KLRF1 | Q9NZS2 | ENSG00000150045 | Killer cell lectin-like receptor subfamily F, member 1 | Glycan Binding/lectin | Not Specific | Glycan ligand not clear |
| 322 | KLRG1 | Q96E93 | ENSG00000139187 | Killer cell lectin-like receptor subfamily G, member 1 | Glycan Binding/lectin | Not Specific | E-cadherin/CDH1, N-cadherin/CDH2 and R-cadherin/CDH4 |
| 323 | KLRG2 | A4D1S0 | ENSG00000188883 | Killer cell lectin-like receptor subfamily G, member 2 | Glycan Binding/lectin | Not Specific | Glycan ligand not clear |
| 324 | KLRK1 | P26718 | ENSG00000213809 | Killer cell lectin-like receptor subfamily K, member 1 | Glycan Binding/lectin | Not Specific | MHC class I-related glycoproteins including MICA, MICB, RAET1E, RAET1G, RAET1L/ULBP6, ULBP1, ULBP2, ULBP3 (ULBP2>ULBP1>ULBP3) and ULBP4 |
| 325 | L1CAM | P32004 | ENSG00000198910 | L1 cell adhesion molecule | Glycan Binding/lectin | Not Specific | Glycan ligand not clear |
| 326 | LARGE1 | O95461 | ENSG00000133424 | Like-glycosyltransferase | Transferase | More than one monosacchrides | Xylose, glucuronate |
| 327 | LARGE2 | Q8N3Y3 | ENSG00000165905 | LARGE xylosyl- and glucuronyltransferase 2 | Transferase | More than one monosacchrides | Xylose, glucuronate |
| 328 | LAYN | Q6UX15 | ENSG00000204381 | Layilin | Glycan Binding/lectin | Hyaluronic acid | Receptor for hyaluronate |
| 329 | LCT | P09848 | ENSG00000115850 | Lactase | Glycosidase | More than one monosacchrides | Lactose to produce D-glucose and D-galactose |
| 330 | LFNG | Q8NES3 | ENSG00000106003 | LFNG O-fucosylpeptide 3-beta-N-acetylglucosaminyltransferase | Transferase | N-acetylglucosamine (GlcNAc) | O-Fuc |
| 331 | LMAN2 | Q12907 | ENSG00000169223 | Vesicular integral-membrane protein VIP36 | Glycan Binding/lectin | More than one monosacchrides | N-acetyl-D-galactosamine and high-mannose type glycans and may also bind to O-linked glycans |
| 332 | LGALS1 | P09382 | ENSG00000100097 | Lectin, galactoside-binding, soluble, 1 | Glycan Binding/lectin | More than one monosacchrides | beta-galactoside and a wide array of complex carbohydrates |
| 333 | LGALS12 | Q96DT0 | ENSG00000133317 | Lectin, galactoside-binding, soluble, 12 | Glycan Binding/lectin | Lactose | Lactose |
| 334 | LGALS13 | Q9UHV8 | ENSG00000105198 | Lectin, galactoside-binding, soluble, 13 | Glycan Binding/lectin | More than one monosacchrides | Beta-galactoside and lactose |
| 335 | LGALS14 | Q8TCE9 | ENSG00000006659 | Lectin, galactoside-binding, soluble, 14 | Glycan Binding/lectin | More than one monosacchrides | Beta-galactoside and lactose |
| 336 | LGALS2 | P05162 | ENSG00000100079 | Lectin, galactoside-binding, soluble, 2 | Glycan Binding/lectin | Beta-galactoside | |
| 337 | LGALS3 | P17931 | ENSG00000131981 | Lectin, galactoside-binding, soluble, 3 | Glycan Binding/lectin | Galactose | Galactose-specific lectin which binds IgE |
| 338 | LGALS4 | P56470 | ENSG00000171747 | Lectin, galactoside-binding, soluble, 4 | Glycan Binding/lectin | Lactose | Lactose and a related range of sugars |
| 339 | LGALS7 | P47929 | ENSG00000205076 | Lectin, galactoside-binding, soluble, 7 | Glycan Binding/lectin | Beta-galactoside | |
| 340 | LGALS8 | O00214 | ENSG00000116977 | Lectin, galactoside-binding, soluble, 8 | Glycan Binding/lectin | Beta-galactoside | |
| 341 | LGALS9 | O00182 | ENSG00000168961 | Lectin, galactoside-binding, soluble, 9 | Glycan Binding/lectin | Beta-galactoside | |
| 342 | LMAN1 | P49257 | ENSG00000074695 | Lectin, mannose-binding, 1/Protein ERGIC-53 | Glycan Binding/lectin | Mannose | Mannose on glycoproteins, glycolipids, or glycosylphosphatidyl inositol anchors |
| 343 | LMAN1L | Q9HAT1 | ENSG00000140506 | Lectin, mannose-binding, 1 like | Glycan Binding/lectin | Mannose | Mannose |
| 344 | LMAN2 | Q12907 | ENSG00000169223 | Lectin, mannose-binding 2 | Glycan Binding/lectin | Mannose | intracellular lectin, N-acetyl-D-galactosamine and high-mannose type glycans and may also bind to O-linked glycans |
| 345 | LMAN2L | Q9H0V9 | ENSG00000114988 | Lectin, mannose-binding 2-like | Glycan Binding/lectin | Mannose | regulator of LMAN1L |
| 346 | LY75 | O60449 | ENSG00000054219 | Lymphocyte antigen 75 | Glycan Binding/lectin | Not Specific | Glycan ligand not clear |
| 347 | M6PR | P20645 | ENSG00000003056 | Mannose-6-phosphate receptor (cation dependent) | Glycan Binding/lectin | Mannose | Receptor |
| 348 | MAG | P20916 | ENSG00000105695 | Myelin associated glycoprotein | Glycan Binding/lectin | Sialic Acid | Preferentially binds to alpha-2,3-linked sialic acid, Neuronal sialic acid-containing gangliosides and to the glycoproteins RTN4R and RTN4RL2 |
| 349 | MAN1A1 | P33908 | ENSG00000111885 | Mannosidase, alpha, class 1A, member 1 | Glycosidase | Mannose | N-linked, Nlinked_Processing |
| 350 | MAN1A2 | O60476 | ENSG00000198162 | Mannosidase, alpha, class 1A, member 2 | Glycosidase | Mannose | N-linked, Nlinked_Processing |
| 351 | MAN1B1 | Q9UKM7 | ENSG00000177239 | Mannosidase, alpha, class 1B, member 1 | Glycosidase | Mannose | N-linked, Nlinked_Processing |
| 352 | MAN1C1 | Q9NR34 | ENSG00000117643 | Mannosidase, alpha, class 1C, member 1 | Glycosidase | Mannose | N-linked, Nlinked_Processing |
| 353 | MAN2A1 | Q16706 | ENSG00000112893 | Mannosidase, alpha, class 2A, member 1 | Glycosidase | Mannose | N-linked, Nlinked_Processing |
| 354 | MAN2A2 | P49641 | ENSG00000196547 | Mannosidase, alpha, class 2A, member 2 | Glycosidase | Mannose | N-linked, Nlinked_Processing |
| 355 | MAN2B1 | O00754 | ENSG00000104774 | Mannosidase, alpha, class 2B, member 1 | Glycosidase | Mannose | N-linked_Lysosome_Degradation |
| 356 | MAN2B2 | Q9Y2E5 | ENSG00000013288 | Mannosidase, alpha, class 2B, member 2 | Glycosidase | Mannose | N-linked_Lysosome_Degradation |
| 357 | MAN2C1 | Q9NTJ4 | ENSG00000140400 | Mannosidase, alpha, class 2C, member 1 | Glycosidase | Mannose | N-linked_Cytosol_Degradation |
| 358 | MANBA | O00462 | ENSG00000109323 | Mannosidase, beta A, lysosomal | Glycosidase | Mannose | N-linked_Lysosome_Degradation |
| 359 | MANBAL | Q9NQG1 | ENSG00000101363 | Mannosidase, beta A, lysosomal-like | Glycosidase | Mannose | N-glycan |
| 360 | MANEA | Q5SRI9 | ENSG00000172469 | Mannosidase, endo-alpha | Glycosidase | Mannose | N-linked, Nlinked_Processing |
| 361 | MANEAL | Q5VSG8 | ENSG00000185090 | Mannosidase, endo-alpha-like | Glycosidase | Mannose | N-glycan |
| 362 | MBL2 | P11226 | ENSG00000165471 | Mannose-binding lectin (protein C) 2, soluble (opsonic defect) | Glycan Binding/lectin | Mannose | |
| 363 | MFNG | O00587 | ENSG00000100060 | MFNG O-fucosylpeptide 3-beta-N-acetylglucosaminyltransferase | Transferase | N-acetylglucosamine (GlcNAc) | O-Fuc |
| 364 | MGAT1 | P26572 | ENSG00000131446 | Mannosyl (alpha-1,3-)-glycoprotein beta-1,2-N-acetylglucosaminyltransferase | Transferase | N-acetylglucosamine (GlcNAc) | N-linked,Nlinked_Branching |
| 365 | MGAT2 | Q10469 | ENSG00000168282 | Mannosyl (alpha-1,6-)-glycoprotein beta-1,2-N-acetylglucosaminyltransferase | Transferase | N-acetylglucosamine (GlcNAc) | N-linked,Nlinked_Branching |
| 366 | MGAT3 | Q09327 | ENSG00000128268 | Mannosyl (beta-1,4-)-glycoprotein beta-1,4-N-acetylglucosaminyltransferase | Transferase | N-acetylglucosamine (GlcNAc) | N-linked,Nlinked_Branching |
| 367 | MGAT4A | Q9UM21 | ENSG00000071073 | Mannosyl (alpha-1,3-)-glycoprotein beta-1,4-N-acetylglucosaminyltransferase, isozyme A | Transferase | N-acetylglucosamine (GlcNAc) | N-linked,Nlinked_Branching |
| 368 | MGAT4B | Q9UQ53 | ENSG00000161013 | Mannosyl (alpha-1,3-)-glycoprotein beta-1,4-N-acetylglucosaminyltransferase, isozyme B | Transferase | N-acetylglucosamine (GlcNAc) | N-linked,Nlinked_Branching |
| 369 | MGAT4C | Q9UBM8 | ENSG00000182050 | Mannosyl (alpha-1,3-)-glycoprotein beta-1,4-N-acetylglucosaminyltransferase, isozyme C (putative) | Transferase | N-acetylglucosamine (GlcNAc) | N-linked,Nlinked_Branching |
| 370 | MGAT5 | Q09328 | ENSG00000152127 | Mannosyl (alpha-1,6-)-glycoprotein beta-1,6-N-acetyl-glucosaminyltransferase | Transferase | N-acetylglucosamine (GlcNAc) | N-linked,Nlinked_Branching |
| 371 | MGAT5B | Q3V5L5 | ENSG00000167889 | Mannosyl (alpha-1,6-)-glycoprotein beta-1,6-N-acetyl-glucosaminyltransferase, isozyme B | Transferase | N-acetylglucosamine (GlcNAc) | O-Man,POMGNT1_Type\|N-linked,Nlinked_Branching |
| 372 | MOGS | Q13724 | ENSG00000115275 | Mannosyl-oligosaccharide glucosidase | Glycosidase | Glucose | distal alpha 1,2-linked glucose residue from the Glc3Man9GlcNAc2 oligosaccharide |
| 373 | MPDU1 | O75352 | ENSG00000129255 | Mannose-P-dolichol utilization defect 1 | Transport | Mannose | Utilization of mannose-dolichol phosphate (Dol-P-Man) |
| 374 | MPI | P34949 | ENSG00000178802 | Mannose phosphate isomerase | Isomerase | Mannose | D-mannose 6-phosphate to D-fructose 6-phosphate |
| 375 | MPPE1 | Q53F39 | ENSG00000154889 | Metallophosphoesterase 1 | Miscellaneous | Not Specific | Transport of GPI-anchor proteins from the endoplasmic reticulum to the Golgi |
| 376 | MRC1 | P22897 | ENSG00000260314 | Mannose receptor, C type 1 | Glycan Binding/lectin | Mannose | Receptor |
| 377 | MRC1L1 | P22897 | ENSG00000260314 | Mannose receptor, C type 1-like 1 | Glycan Binding/lectin | Mannose | Receptor |
| 378 | MRC2 | Q9UBG0 | ENSG00000011028 | Mannose receptor, C type 2 | Glycan Binding/lectin | Mannose | Receptor |
| 379 | NAGA | P17050 | ENSG00000198951 | N-acetylgalactosaminidase, alpha- | Glycosidase | N-acetylgalactosamine (GalNAc) | GSL_Lysosome_Degradation,O-linked_Lysosome_Degradation |
| 380 | NAGK | Q9UJ70 | ENSG00000124357 | N-Acetylglucosamine Kinase | Transferase | Phosphate | Phosphate to N-acetylglucosamine |
| 381 | NAGLU | P54802 | ENSG00000108784 | N-acetylglucosaminidase, alpha- | Glycosidase | N-acetylgalactosamine (GalNAc) | HS_Lysosome_Degradation |
| 382 | NAGPA | Q9UK23 | ENSG00000103174 | N-acetylglucosamine-1-phosphodiester alpha-N-acetylglucosaminidase | Glycosidase | N-acetylgalactosamine (GalNAc) | N-linked, Lysosomal_Targeting |
| 383 | NANS | Q9NR45 | ENSG00000095380 | N-Acetylneuraminate Synthase | Transferase | Sialic Acid | Produces N-acetylneuraminic acid (Neu5Ac) |
| 384 | NCAN | O14594 | ENSG00000130287 | Neurocan | Glycan Binding/lectin | More than one monosacchrides | Neural cell adhesion molecule (N-CAM) and the neuron-glia cell adhesion molecule (Ng-CAM) Chondroitin sulfate proteoglycan; binds to hyaluronic acid |
| 385 | NDST1 | P52848 | ENSG00000070614 | N-deacetylase/N-sulfotransferase (heparan glucosaminyl) 1 | Transferase | Acetate/Sulfate | Remove acetate/ Sulfate group |
| 386 | NDST2 | P52849 | ENSG00000166507 | N-deacetylase/N-sulfotransferase (heparan glucosaminyl) 2 | Transferase | Acetate/Sulfate | Remove acetate/ Sulfate group |
| 387 | NDST3 | O95803 | ENSG00000164100 | N-deacetylase/N-sulfotransferase (heparan glucosaminyl) 3 | Transferase | Acetate/Sulfate | Remove acetate/ Sulfate group |
| 388 | NDST4 | Q9H3R1 | ENSG00000138653 | N-deacetylase/N-sulfotransferase (heparan glucosaminyl) 4 | Transferase | Acetate/Sulfate | Remove acetate/ Sulfate group |
| 389 | NEU1 | Q99519 | ENSG00000204386 | Sialidase 1 (lysosomal sialidase) | Glycosidase | Sialic Acid | N-linked_Lysosome_Degradation,O-linked_Lysosome_Degradation,GSL_Lysosome_Degradation,KS_Lysosome_Degradation |
| 390 | NEU2 | Q9Y3R4 | ENSG00000115488 | Sialidase 2 (cytosolic sialidase) | Glycosidase | Sialic Acid | N-linked_Cytosol_Degradation,O-linked_Cytosol_Degradation,GSL_Cytosol_Degradation,KS_Cytosol_Degradation |
| 391 | NEU3 | Q9UQ49 | ENSG00000162139 | Sialidase 3 (membrane sialidase) | Glycosidase | Sialic Acid | N-linked_Lysosome_Degradation,O-linked_Lysosome_Degradation,GSL_Lysosome_Degradation,KS_Lysosome_Degradation |
| 392 | NEU4 | Q8WWR8 | ENSG00000204099 | Sialidase 4 | Glycosidase | Sialic Acid | N-linked_Lysosome_Degradation,O-linked_Lysosome_Degradation,GSL_Lysosome_Degradation,KS_Lysosome_Degradation |
| 393 | OGA | O60502 | ENSG00000198408 | O-GlcNAcase | Glycosidase | N-acetylglucosamine (GlcNAc) | O-GlcNAc |
| 394 | OGT | O15294 | ENSG00000147162 | O-linked N-acetylglucosamine (GlcNAc) transferase (UDP-N-acetylglucosamine:polypeptide-N-acetylglucosaminyl transferase) | Transferase | N-acetylglucosamine (GlcNAc) | N-acetylglucosamine (GlcNAc) on serine |
| 395 | OS9 | Q13438 | ENSG00000135506 | OS9 Endoplasmic Reticulum Lectin | Glycan Binding/lectin | Not Specific | Terminally misfolded non-glycosylated proteins as well as improperly folded glycoproteins |
| 396 | OLR1 | P78380 | ENSG00000173391 | Oxidized low density lipoprotein (lectin-like) receptor 1 | Glycan Binding/lectin | More than one monosacchrides | Advanced glycation end (AGE) products, Oxidatively modified low density lipoprotein (oxLDL) |
| 397 | PAPSS1 | O43252 | ENSG00000138801 | Bifunctional 3'-phosphoadenosine 5'-phosphosulfate synthase 1 | Transferase | Sulfate | 1. Sulfate group to ATP to yield adenosine 5'-phosphosulfate (APS). 2. transfer of a phosphate group from ATP to APS yielding 3'-phosphoadenylylsulfate (PAPS |
| 398 | PAPSS2 | O95340 | ENSG00000198682 | 3'-Phosphoadenosine 5'-Phosphosulfate Synthase 2 | Transferase | Sulfate | 1. Sulfate group to ATP to yield adenosine 5'-phosphosulfate (APS). 2. transfer of a phosphate group from ATP to APS yielding 3'-phosphoadenylylsulfate (PAPS |
| 399 | PECAM1 | P16284 | ENSG00000261371 | Platelet/endothelial cell adhesion molecule | Glycan Binding/lectin | Not Specific | Specific glycan ligand not known |
| 400 | PGAP1 | Q75T13 | ENSG00000197121 | Post-GPI attachment to proteins 1 | Transport | Inositol | Inositol deacylation of GPI-anchored proteins |
| 401 | PGAP4 | Q9BRR3 | ENSG00000165152 | Post-GPI attachment to proteins factor 4 | Transferase | N-acetylgalactosamine (GalNAc) | GalNAc to GPI |
| 402 | PGGHG | Q32M88 | ENSG00000142102 | Protein-glucosylhydrolysine glucosidase | Glycosidase | Glucose | Hydrolysis of glucose from the disaccharide |
| 403 | PGLYRP1 | O75594 | ENSG00000008438 | Peptidoglycan recognition protein 1 | Glycan Binding/lectin | Non-Human glycans | Murein peptidoglycans (PGN) of Gram-positive bacteria |
| 404 | PGLYRP2 | Q96PD5 | ENSG00000161031 | Peptidoglycan recognition protein 2 | Glycan Binding/lectin | Non-Human glycans | Digesting biologically active peptidoglycan (PGN) |
| 405 | PGLYRP3 | Q96LB9 | ENSG00000159527 | Peptidoglycan recognition protein 3 | Glycan Binding/lectin | Non-Human glycans | Murein peptidoglycans (PGN) of Gram-positive bacteria, also to Gram-negative bacteria |
| 406 | PGLYRP4 | Q96LB8 | ENSG00000163218 | Peptidoglycan recognition protein 4 | Glycan Binding/lectin | Non-Human glycans | Murein peptidoglycans (PGN) of Gram-positive bacteria, also to Gram-negative bacteria |
| 407 | PGM1 | P36871 | ENSG00000079739 | Phosphoglucomutase-1 | Isomerase | Glucose | Breakdown and synthesis of glucose |
| 408 | PGM2 | Q96G03 | ENSG00000169299 | Phosphoglucomutase-2 | Isomerase | More than one monosacchrides | Nucleoside breakdown of ribose-1-phosphate and deoxyribose-1-phosphate to the corresponding 5-phosphopentose, interconversion of glucose-1-phosphate into glucose-6-phosphate |
| 409 | PGM3 | O95394 | ENSG00000013375 | Phosphoacetylglucosamine mutase | Isomerase | N-acetylglucosamine (GlcNAc) | Conversion of GlcNAc-6-P into GlcNAc-1-P |
| 410 | PIGA | P37287 | ENSG00000165195 | Phosphatidylinositol glycan anchor biosynthesis, class A | Transferase | N-acetylglucosamine (GlcNAc) | Subunit of enzyme which transfer of N-acetylglucosamine from UDP-N-acetylglucosamine to phosphatidylinositol |
| 411 | PIGB | Q92521 | ENSG00000069943 | Phosphatidylinositol glycan anchor biosynthesis, class B | Transferase | Mannose | Transfers the third alpha-1,2-mannose to Man2-GlcN-acyl-PI during GPI precursor assembly |
| 412 | PIGC | Q92535 | ENSG00000135845 | Phosphatidylinositol glycan anchor biosynthesis, class C | Transferase | N-acetylglucosamine (GlcNAc) | Subunit of enzyme which transfer of N-acetylglucosamine from UDP-N-acetylglucosamine to phosphatidylinositol |
| 413 | PIGF | Q07326 | ENSG00000151665 | Phosphatidylinositol glycan anchor biosynthesis, class F | Transferase | Mannose | Ethanolamine phosphate to the third mannose of GPI |
| 414 | PIGG | Q5H8A4 | ENSG00000174227 | Phosphatidylinositol glycan anchor biosynthesis, class G | Transferase | Mannose | Ethanolamine phosphate to the GPI second mannose |
| 415 | PIGH | Q14442 | ENSG00000100564 | Phosphatidylinositol glycan anchor biosynthesis, class H | Transferase | N-acetylglucosamine (GlcNAc) | N-acetylglucosamine from UDP-N-acetylglucosamine to phosphatidylinositol |
| 416 | PIGK | Q92643 | ENSG00000142892 | Phosphatidylinositol glycan anchor biosynthesis, class K | Transamidase | Not Specific | GPI anchoring in the endoplasmic reticulum, transfer of GPI to proteins |
| 417 | PIGL | Q9Y2B2 | ENSG00000108474 | Phosphatidylinositol glycan anchor biosynthesis, class L | Miscellaneous | N-acetylglucosamine (GlcNAc) | De-N-acetylation of N-acetylglucosaminyl-phosphatidylinositol |
| 418 | PIGM | Q9H3S5 | ENSG00000143315 | Phosphatidylinositol glycan anchor biosynthesis, class M | Transferase | Mannose | First alpha-1,4-mannose to GlcN-acyl-PI during GPI precursor assembly |
| 419 | PIGN | O95427 | ENSG00000197563 | Phosphatidylinositol glycan anchor biosynthesis, class N | Transferase | Mannose | Ethanolamine phosphate to the first alpha-1,4-linked mannose of the glycosylphosphatidylinositol precursor of GPI-anchor |
| 420 | PIGO | Q8TEQ8 | ENSG00000165282 | Phosphatidylinositol glycan anchor biosynthesis, class O | Transferase | Mannose | Ethanolamine phosphate to the GPI third mannose which links the GPI-anchor to the C-terminus of the proteins by an amide bond |
| 421 | PIGP | P57054 | ENSG00000185808 | Phosphatidylinositol glycan anchor biosynthesis, class P | Transferase | N-acetylglucosamine (GlcNAc) | N-acetylglucosamine from UDP-N-acetylglucosamine to phosphatidylinositol and participates in the first step of GPI biosynthesis |

| | | | | | | | |
|---|---|---|---|---|---|---|---|
| 422 | PIGQ | Q9BRB3 | ENSG00000007541 | Phosphatidylinositol glycan anchor biosynthesis, class Q | Transferase | N-acetylglucosamine (GlcNAc) | N-acetylglucosamine from UDP-N-acetylglucosamine to phosphatidylinositol and participates in the first step of GPI biosynthesis |
| 423 | PIGR | P01833 | ENSG00000162896 | Polymeric immunoglobulin receptor | Transport | More than one monosacchrides | Binds polymeric IgA and IgM, transcytosis |
| 424 | PIGS | Q96S52 | ENSG00000087111 | Phosphatidylinositol glycan anchor biosynthesis, class S | Regulator | Not Specific | Transfer of GPI to proteins, One component |
| 425 | PIGT | Q969N2 | ENSG00000124155 | Phosphatidylinositol glycan anchor biosynthesis, class T | Regulator | Not Specific | Transfer of GPI to proteins, One component |
| 426 | PIGU | Q9H490 | ENSG00000101464 | Phosphatidylinositol glycan anchor biosynthesis, class U | Regulator | Not Specific | Transfer of GPI to proteins, One component |
| 427 | PIGV | Q9NUD9 | ENSG00000060642 | Phosphatidylinositol glycan anchor biosynthesis, class V | Transferase | Mannose | Second mannose to the glycosylphosphatidylinositol during GPI precursor assembly |
| 428 | PIGW | Q7Z7B1 | ENSG00000277161 | Phosphatidylinositol glycan anchor biosynthesis, class W | Transport | More than one monosacchrides | GPI-anchored proteins to the plasma membrane, acetylates the inositol ring of phosphatidylinositol during biosynthesis of GPI-anchor |
| 429 | PIGX | Q8TBF5 | ENSG00000163964 | Phosphatidylinositol glycan anchor biosynthesis, class X | Regulator | Mannose | Stabilizing the mannosyltransferase PIGM |
| 430 | PIGY | Q3MUY2 | ENSG00000255072 | Phosphatidylinositol glycan anchor biosynthesis, class Y | Transferase | N-acetylglucosamine (GlcNAc) | component of enzyme,N-acetylglucosamine from UDP-N-acetylglucosamine to phosphatidylinositol and participates in the first step of GPI biosynthesis |
| 431 | PIGZ | Q86VD9 | ENSG00000119227 | Phosphatidylinositol glycan anchor biosynthesis, class Z | Transferase | Mannose | Fourth mannose to some trimannosyl-GPIs during GPI precursor assembly |
| 432 | PLOD1 | Q02809 | ENSG00000083444 | Procollagen-lysine,2-oxoglutarate 5-dioxygenase 1 | Miscellaneous | Collagen | Hydroxylation of lysine residues in collagen alpha chains |
| 433 | PLOD2 | O00469 | ENSG00000152952 | Procollagen-lysine,2-oxoglutarate 5-dioxygenase 2 | Miscellaneous | Collagen | Forms hydroxylysine residues in -Xaa-Lys-Gly- sequences in collagens |
| 434 | PLOD3 | O60568 | ENSG00000106397 | Multifunctional procollagen lysine hydroxylase and glycosyltransferase LH3 | Transferase | Collagen | Post-translational modifications on Lys residues in procollagen, Forms hydroxylysine residues in -Xaa-Lys-Gly- sequences in collagens |
| 435 | PMM1 | Q92871 | ENSG00000100417 | Phosphomannomutase 1 | Isomerase | Mannose | alpha-D-mannose 1-phosphate to D-mannose 6-phosphate |
| 436 | PMM2 | O15305 | ENSG00000140650 | Phosphomannomutase 2 | Isomerase | Mannose | alpha-D-mannose 1-phosphate to D-mannose 6-phosphate |
| 437 | POFUT1 | Q9H488 | ENSG00000101346 | GDP-fucose protein O-fucosyltransferase 1 | Transferase | Fucose | Fucose to Serine |
| 438 | POFUT2 | Q9Y2G5 | ENSG00000186866 | GDP-fucose protein O-fucosyltransferase 2 | Transferase | Fucose | Fucose to Serine |
| 439 | POGLUT1 | Q8NBL1 | ENSG00000163389 | Protein O-glucosyltransferase 1 | Transferase | More than one monosacchrides | glucose and xylose |
| 440 | POGLUT2 | Q6UW63 | ENSG00000134901 | Protein O-glucosyltransferase 2 | Transferase | Glucose | O-Glc |
| 441 | POGLUT3 | Q7Z4H8 | ENSG00000178202 | Protein O-glucosyltransferase 3 | Transferase | Glucose | O-Glc |
| 442 | POMGNT1 | Q8WZA1 | ENSG00000085998 | Protein O-linked mannose beta1,2-N-acetylglucosaminyltransferase | Transferase | N-acetylglucosamine (GlcNAc) | N-acetylglucosamine to O-linked mannose on glycoproteins |
| 443 | POMGNT2 | Q8NAT1 | ENSG00000144647 | Protein O-linked-mannose beta-1,4-N-acetylglucosaminyltransferase 2 | Transferase | N-acetylglucosamine (GlcNAc) | UDP-N-acetyl-D-glucosamine to the 4-position of the mannose to generate N-acetyl-D-glucosamine-beta-1,4-O-D-mannosylprotein |
| 444 | POMK | Q9H5K3 | ENSG00000185900 | Protein O-mannose kinase | Kinase | Phosphate | Phosphorylation at the 6-position of an O-mannose of the trisaccharide (N-acetylgalactosamine (GalNAc)-beta-1,3-N-acetylglucosamine (GlcNAc)-beta-1,4-mannose) |
| 445 | POMT1 | Q9Y6A1 | ENSG00000130714 | Protein-O-mannosyltransferase 1 | Transferase | Mannose | Mannose to the hydroxyl group of serine or threonine |
| 446 | POMT2 | Q9UKY4 | ENSG00000009830 | Protein-O-mannosyltransferase 2 | Transferase | Mannose | Mannose to the hydroxyl group of serine or threonine |
| 447 | PRKCSH | P14314 | ENSG00000130175 | Glucosidase 2 subunit beta | Glycosidase | Glucose | Cleaves sequentially the 2 innermost alpha-1,3-linked glucose residues from the Glc2Man9GlcNAc2oligosaccharide precursor of immature glycoproteins |
| 448 | PXYLP1 | Q8TE99 | ENSG00000155893 | 2-phosphoxylose phosphatase 1 | Phosphatase | Phosphate | 2-O-dephosphorylation of xylose |
| 449 | RENBP | P51606 | ENSG00000102032 | N-acylglucosamine 2-epimerase | Epimerase | N-acetylglucosamine (GlcNAc) | Interconversion of N-acetylglucosamine to N-acetylmannosamine |
| 450 | RFNG | Q9Y644 | ENSG00000169733 | RFNG O-fucosylpeptide 3-beta-N-acetylglucosaminyltransferase | Transferase | N-acetylglucosamine (GlcNAc) | |
| 451 | RFT1 | Q96AA3 | ENSG00000163933 | RFT1 homolog (S. cerevisiae) | transport | More than one monosacchrides | Translocation of oligosaccharide from the cytoplasmic side to the lumenal side of the endoplasmic reticulum membrane |
| 452 | RPN1 | P04843 | ENSG00000163902 | Ribophorin I | Transferase | More than one monosacchrides | A subunit, Transfer of a defined glycan (Glc3Man9GlcNAc2 in eukaryotes) from the lipid carrier dolichol-pyrophosphate to an asparagine residue within an Asn-X-Ser/Thr |
| 453 | RPN2 | P04844 | ENSG00000118705 | Ribophorin II | Transferase | More than one monosacchrides | A subunit, Transfer of a defined glycan (Glc3Man9GlcNAc2 in eukaryotes) from the lipid carrier dolichol-pyrophosphate to an asparagine residue within an Asn-X-Ser/Thr |
| 454 | RXYLT1 | Q9Y2B1 | ENSG00000118600 | Ribitol-5-phosphate xylosyltransferase 1 | Transferase | Xylose | UDP-D-xylose to ribitol 5-phosphate (Rbo5P) to form the Xylbeta1-4Rbo5P linkage on O-mannosyl glycan |
| 455 | SEL1L | Q9UBV2 | ENSG00000071537 | Protein sel-1 homolog 1 | Regulator | Not Specific | Endoplasmic reticulum quality control (ERQC) |
| 456 | SELE | P16581 | ENSG00000007908 | Selectin E | Glycan Binding/lectin | Sialic Acid ? | Sialyl-Lewisa (sLea) and sialyl-Lewisx (sLex) |
| 457 | SELL | P14151 | ENSG00000188404 | Selectin L | Glycan Binding/lectin | Sialic Acid ? | sialyl-Lewisx (sLex) |
| 458 | SELP | P16109 | ENSG00000174175 | Selectin P (granule membrane protein 140kDa, antigen CD62) | Glycan Binding/lectin | Sialic Acid ? | sialyl-Lewisx (sLex) |
| 459 | SGSH | P51688 | ENSG00000181523 | N-sulphoglucosamine sulphohydrolase | Miscellaneous | Sulfate | Lysosomal heparan sulfate degradation |
| 460 | SIAE | Q9HAT2 | ENSG00000110013 | Sialate O-acetylesterase | Miscellaneous | Sialic Acid | Removal of O-acetyl ester groups from position 9 of sialic acid |
| 461 | SIGLEC1 | Q9BZZ2 | ENSG00000088827 | Sialic acid binding Ig-like lectin 1, sialoadhesin | Glycan Binding/lectin | Sialic Acid | Preferentially binds to alpha-2,3-linked sialic acid |
| 462 | SIGLEC10 | Q96LC7 | ENSG00000142512 | Sialic acid binding Ig-like lectin 10 | Glycan Binding/lectin | Sialic Acid | alpha-2,3- or alpha-2,6-linked sialic acid |
| 463 | SIGLEC11 | Q96RL6 | ENSG00000161640 | Sialic acid binding Ig-like lectin 11 | Glycan Binding/lectin | Sialic Acid | Preferentially binds to alpha-2,8-linked sialic acid |
| 464 | SIGLEC12 | Q96PQ1 | ENSG00000254521 | Sialic acid binding Ig-like lectin 12 | Glycan Binding/lectin | Sialic Acid | - |
| 465 | SIGLEC14 | Q08ET2 | ENSG00000254415 | Sialic acid binding Ig-like lectin 14 | Glycan Binding/lectin | Sialic Acid | alpha-2,6-linked sialic acid, alpha-2,8-linked sialic acid |
| 466 | SIGLEC16 | A6NMB1 | ENSG00000161643 | Sialic acid binding Ig-like lectin 16 | Glycan Binding/lectin | Sialic Acid | Preferentially binds to alpha-2,8-linked sialic acid |
| 467 | SIGLEC2 | P20273 | ENSG00000012124 | Sialic acid binding Ig-like lectin 2 | Glycan Binding/lectin | Sialic Acid | Preferentially binds to alpha-2,6-linked sialic acid |
| 468 | SIGLEC3 | P20138 | ENSG00000105383 | Sialic acid binding Ig-like lectin 3 | Glycan Binding/lectin | Sialic Acid | Preferentially recognizes and binds alpha-2,3- and more avidly alpha-2,6-linked sialic acid-bearing glycans |
| 469 | SIGLEC5 | O15389 | ENSG00000105501 | Sialic acid binding Ig-like lectin 5 | Glycan Binding/lectin | Sialic Acid | Binds equally to alpha-2,3-linked and alpha-2,6-linked sialic acid |
| 470 | SIGLEC6 | O43699 | ENSG00000105492 | Sialic acid binding Ig-like lectin 6 | Glycan Binding/lectin | Sialic Acid | Binds to alpha-2,6-linked sialic acid |
| 471 | SIGLEC7 | Q9Y286 | ENSG00000168995 | Sialic acid binding Ig-like lectin 7 | Glycan Binding/lectin | Sialic Acid | Preferentially binds to alpha-2,3- and alpha-2,6-linked sialic acid. Also binds disialogangliosides (disialogalactosyl globoside, disialyl lactotetraosylceramide and disialyl GalNAc lactotetraoslylceramide). |
| 472 | SIGLEC8 | Q9NYZ4 | ENSG00000105366 | Sialic acid binding Ig-like lectin 8 | Glycan Binding/lectin | Sialic Acid | Preferentially binds to alpha-2,3-linked sialic acid. Also binds to alpha-2,6-linked sialic acid. |
| 473 | SIGLEC9 | Q9Y336 | ENSG00000129450 | Sialic acid binding Ig-like lectin 9 | Glycan Binding/lectin | Sialic Acid | Preferentially binds to alpha-2,3- or alpha-2,6-linked sialic acid |
| 474 | SLC2A1 | P11166 | ENSG00000117394 | Solute carrier family 2, facilitated glucose transporter member 1 | Transport | Glucose | |
| 475 | SLC2A10 | O95528 | ENSG00000197496 | Solute carrier family 2, facilitated glucose transporter member 10 | Transport | Glucose | |
| 476 | SLC2A12 | Q8TD20 | ENSG00000146411 | Solute carrier family 2, facilitated glucose transporter member 12 | Transport | Glucose | |
| 477 | SLC2A2 | P11168 | ENSG00000163581 | Solute carrier family 2, facilitated glucose transporter member 2 | Transport | Glucose | |
| 478 | SLC2A3 | P11169 | ENSG00000059804 | Solute carrier family 2, facilitated glucose transporter member 3 | Transport | Glucose | |
| 479 | SLC2A4 | P14672 | ENSG00000181856 | Solute carrier family 2, facilitated glucose transporter member 4 | Transport | Glucose | |
| 480 | SLC2A5 | P22732 | ENSG00000142583 | Solute carrier family 2, facilitated glucose transporter member 5 | Transport | Glucose | |
| 481 | SLC2A8 | Q9NY64 | ENSG00000136856 | Solute carrier family 2, facilitated glucose transporter member 8 | Transport | Glucose | |
| 482 | SLC33A1 | O00400 | ENSG00000169359 | Solute carrier family 33 (acetyl-CoA transporter), member 1 | Transport | Acetate | Acetyl-CoA transporter, Necessary for O-acetylation of gangliosides |
| 483 | SLC35A1 | P78382 | ENSG00000164414 | Solute carrier family 35 (CMP-sialic acid transporter), member A1 | Transport | Sialic Acid | CMP-Neu5Ac Transport |
| 484 | SLC35A2 | P78381 | ENSG00000102100 | Solute carrier family 35 (UDP-galactose transporter), member A2 | Transport | Galactose | UDP-Gal Transport |
| 485 | SLC35A3 | Q9Y2D2 | ENSG00000117620 | Solute carrier family 35 (UDP-N-acetylglucosamine (UDP-GlcNAc) transporter), member A3 | Transport | N-acetylglucosamine (GlcNAc) | UDP-GlcNAc Transport |
| 486 | SLC35B1 | P78383 | ENSG00000121073 | Solute carrier family 35, member B1 | Transport | | ATP:ADP antiporter |
| 487 | SLC35B2 | Q8TB61 | ENSG00000157593 | Solute carrier family 35, member B2 | Transport | | PAPS Transport |
| 488 | SLC35B3 | Q9H1N7 | ENSG00000124786 | Solute carrier family 35, member B3 | Transport | | PAPS Transport |
| 489 | SLC35B4 | Q969S0 | ENSG00000205060 | Solute carrier family 35, member B4 | Transport | More than one monosacchrides | UDP-Xyl Transport, UDP-GlcNAc Transport |
| 490 | SLC35C1 | Q96A29 | ENSG00000181830 | Solute carrier family 35, member C1 | Transport | Fucose | GDP-Fuc Transport |
| 491 | SLC35D1 | Q9NTN3 | ENSG00000116704 | Solute carrier family 35 (UDP-glucuronic acid/UDP-N-acetylgalactosamine dual transporter), member D1 | Transport | More than one monosacchrides | Antiporter that transports nucleotide sugars across the endoplasmic reticulum (ER) membrane in exchange for either their cognate nucleoside monophosphate or another nucleotide sugar |
| 492 | SLC35D2 | Q76EJ3 | ENSG00000130958 | Solute carrier family 35, member D2 | Transport | More than one monosacchrides | Nucleotide sugar antiporter transporting UDP-N-acetylglucosamine (UDP-GlcNAc) and UDP-glucose (UDP-Glc) from the cytosol into the lumen of the Golgi in exchange of UMP |
| 493 | SPAM1 | P38567 | ENSG00000106304 | Sperm adhesion molecule 1 (PH-20 hyaluronidase, zona pellucida binding) | Glycosidase | Hyaluronic acid | Random hydrolysis of (1->4)-linkages between N-acetyl-beta-D-glucosamine and D-glucuronate residues in hyaluronate |
| 494 | ST3GAL1 | Q11201 | ENSG00000008513 | ST3 beta-galactoside alpha-2,3-sialyltransferase 1 | Transferase | Sialic Acid | |
| 495 | ST3GAL2 | Q16842 | ENSG00000157350 | ST3 beta-galactoside alpha-2,3-sialyltransferase 2 | Transferase | Sialic Acid | |
| 496 | ST3GAL3 | Q11203 | ENSG00000126091 | ST3 beta-galactoside alpha-2,3-sialyltransferase 3 | Transferase | Sialic Acid | |
| 497 | ST3GAL4 | Q11206 | ENSG00000110080 | ST3 beta-galactoside alpha-2,3-sialyltransferase 4 | Transferase | Sialic Acid | |
| 498 | ST3GAL5 | Q11201 | ENSG00000115525 | ST3 beta-galactoside alpha-2,3-sialyltransferase 5 | Transferase | Sialic Acid | Capping_O-GalNAc |
| 499 | ST3GAL6 | Q16842 | ENSG00000064225 | ST3 beta-galactoside alpha-2,3-sialyltransferase 6 | Transferase | Sialic Acid | Capping_Ganglioside |
| 500 | ST6GAL1 | Q11203 | ENSG00000073849 | ST6 beta-galactosamide alpha-2,6-sialyltranferase 1 | Transferase | Sialic Acid | Capping_N-linked,Capping_O-GalNAc |
| 501 | ST6GAL2 | Q11206 | ENSG00000144057 | ST6 beta-galactosamide alpha-2,6-sialyltranferase 2 | Transferase | Sialic Acid | Capping_N-linked,Capping_O-GalNAc |
| 502 | ST6GALNAC1 | Q9NSC7 | ENSG00000070526 | ST6 (alpha-N-acetyl-neuraminyl-2,3-beta-galactosyl-1,3)-N-acetylgalactosaminide alpha-2,6-sialyltransferase 1 | Transferase | Sialic Acid | Capping_O-GalNAc |
| 503 | ST6GALNAC2 | Q9UJ37 | ENSG00000070731 | ST6 (alpha-N-acetyl-neuraminyl-2,3-beta-galactosyl-1,3)-N-acetylgalactosaminide alpha-2,6-sialyltransferase 2 | Transferase | Sialic Acid | Capping_O-GalNAc |
| 504 | ST6GALNAC3 | Q8NDV1 | ENSG00000184005 | ST6 (alpha-N-acetyl-neuraminyl-2,3-beta-galactosyl-1,3)-N-acetylgalactosaminide alpha-2,6-sialyltransferase 3 | Transferase | Sialic Acid | Capping_O-GalNAc,Capping_Ganglioside |
| 505 | ST6GALNAC4 | Q9H4F1 | ENSG00000136840 | ST6 (alpha-N-acetyl-neuraminyl-2,3-beta-galactosyl-1,3)-N-acetylgalactosaminide alpha-2,6-sialyltransferase 4 | Transferase | Sialic Acid | Capping_O-GalNAc,Capping_Ganglioside |
| 506 | ST6GALNAC5 | Q9BVH7 | ENSG00000117069 | ST6 (alpha-N-acetyl-neuraminyl-2,3-beta-galactosyl-1,3)-N-acetylgalactosaminide alpha-2,6-sialyltransferase 5 | Transferase | Sialic Acid | Capping_Ganglioside |
| 507 | ST6GALNAC6 | Q969X2 | ENSG00000160408 | ST6 (alpha-N-acetyl-neuraminyl-2,3-beta-galactosyl-1,3)-N-acetylgalactosaminide alpha-2,6-sialyltransferase 6 | Transferase | Sialic Acid | Capping_Ganglioside |
| 508 | ST8SIA1 | Q92185 | ENSG00000111728 | ST8 alpha-N-acetyl-neuraminide alpha-2,8-sialyltransferase 1 | Transferase | Sialic Acid | Capping_Ganglioside |
| 509 | ST8SIA2 | Q92186 | ENSG00000140557 | ST8 alpha-N-acetyl-neuraminide alpha-2,8-sialyltransferase 2 | Transferase | Sialic Acid | Capping_N-linked,Capping_O-GalNAc |
| 510 | ST8SIA3 | O43173 | ENSG00000177511 | ST8 alpha-N-acetyl-neuraminide alpha-2,8-sialyltransferase 3 | Transferase | Sialic Acid | Capping_N-linked,Capping_Ganglioside |
| 511 | ST8SIA4 | Q92187 | ENSG00000113532 | ST8 alpha-N-acetyl-neuraminide alpha-2,8-sialyltransferase 4 | Transferase | Sialic Acid | Capping_N-linked,Capping_O-GalNAc |
| 512 | ST8SIA5 | O15466 | ENSG00000101638 | ST8 alpha-N-acetyl-neuraminide alpha-2,8-sialyltransferase 5 | Transferase | Sialic Acid | Capping_Ganglioside |
| 513 | ST8SIA6 | P61647 | ENSG00000148488 | ST8 alpha-N-acetyl-neuraminide alpha-2,8-sialyltransferase 6 | Transferase | Sialic Acid | Capping_O-GalNAc |
| 514 | STT3A | P46977 | ENSG00000134910 | STT3, subunit of the oligosaccharyltransferase complex, homolog A (S. cerevisiae) | Transferase | More than one monosacchrides | Subunit, first step in protein N-glycosylation |
| 515 | STT3B | Q8TCJ2 | ENSG00000163527 | STT3, subunit of the oligosaccharyltransferase complex, homolog B (S. cerevisiae) | Transferase | More than one monosacchrides | Subunit, first step in protein N-glycosylation |
| 516 | SULF1 | Q8IWU6 | ENSG00000137573 | Extracellular sulfatase Sulf-1 | Sulfatase | Sulfate | Extracellular_Degradation |
| 517 | SULF2 | Q8IWU5 | ENSG00000196562 | Extracellular sulfatase Sulf-2 | Sulfatase | Sulfate | Extracellular_Degradation |
| 518 | TMTC1 | Q8IUR5 | ENSG00000133687 | Protein O-mannosyl-transferase TMTC1 | Transferase | Mannose | O-Man on cadherin |
| 519 | TMTC2 | Q8N394 | ENSG00000179104 | Protein O-mannosyl-transferase TMTC2 | Transferase | Mannose | O-Man on cadherin |
| 520 | TMTC3 | Q6ZXV5 | ENSG00000139324 | Protein O-mannosyl-transferase TMTC3 | Transferase | Mannose | O-Man on cadherin |
| 521 | TMTC4 | Q5T4D3 | ENSG00000125247 | Protein O-mannosyl-transferase TMTC4 | Transferase | Mannose | O-Man on cadherin |
| 522 | TPST1 | O60507 | ENSG00000169902 | Protein-tyrosine sulfotransferase 1 | Transferase | Sulfate | Protein_Sulfation |
| 523 | TPST2 | O60704 | ENSG00000128294 | Protein-tyrosine sulfotransferase 2 | Transferase | Sulfate | Protein_Sulfation |
| 524 | UAP1 | Q16222 | ENSG00000117143 | UDP-N-acetylhexosamine pyrophosphorylase | Transferase | MOre than one monosacchrides | UDP- to GlcNAc and GalNac |
| 525 | UGCG | Q16739 | ENSG00000148154 | Ceramide glucosyltransferase | Transferase | Glucose | Glucose to ceramide |
| 526 | UGCGL1 | Q9NYU2 | ENSG00000136731 | UDP-glucose ceramide glucosyltransferase-like 1 | Transferase | Glucose | Reglucosylates single N-glycans on misfolded protein |
| 527 | UGCGL2 | Q9NYU1 | ENSG00000102595 | UDP-glucose ceramide glucosyltransferase-like 2 | Transferase | Glucose | Reglucosylates single N-glycans on misfolded protein |
| 528 | UGDH | O60701 | ENSG00000109814 | UDP-glucose 6-dehydrogenase | Miscellaneous | Glucuronic acid (GlcA) | Formation of UDP-alpha-D-glucuronate |
| 529 | UGGT1 | Q9NYU2 | ENSG00000136731 | UDP-glucose:glycoprotein glucosyltransferase 1 | Transferase | Glucose | Reglucosylates single N-glycans on misfolded protein |
| 530 | UGGT2 | Q9NYU1 | ENSG00000102595 | UDP-glucose:glycoprotein glucosyltransferase 2 | Transferase | Glucose | Reglucosylates single N-glycans on misfolded protein |
| 531 | UGP2 | Q16851 | ENSG00000169764 | UTP-glucose-1-phosphate uridylyltransferase | Transferase | Glucose | glucose-1-phosphate into UDP-glucose |
| 532 | UGT1A1 | P22309 | ENSG00000241635 | UDP-glucuronosyltransferase 1A1 | Transferase | Glucuronic acid (GlcA) | Glucuronidation |
| 533 | UGT1A10 | Q9HAW8 | ENSG00000242515 | UDP-glucuronosyltransferase 1A10 | Transferase | Glucuronic acid (GlcA) | Glucuronidation |
| 534 | UGT1A3 | P35503 | ENSG00000288702 | UDP-glucuronosyltransferase 1A3 | Transferase | Glucuronic acid (GlcA) | Glucuronidation |
| 535 | UGT1A4 | P22310 | ENSG00000244474 | UDP-glucuronosyltransferase 1A4 | Transferase | Glucuronic acid (GlcA) | Glucuronidation |
| 536 | UGT1A5 | P35504 | ENSG00000288705 | UDP-glucuronosyltransferase 1A5 | Transferase | Glucuronic acid (GlcA) | Glucuronidation |
| 537 | UGT1A6 | P19224 | ENSG00000167165 | UDP-glucuronosyltransferase 1-6 | Transferase | Glucuronic acid (GlcA) | Glucuronidation |
| 538 | UGT1A7 | Q9HAW7 | ENSG00000244122 | UDP-glucuronosyltransferase 1A7 | Transferase | Glucuronic acid (GlcA) | Glucuronidation |
| 539 | UGT1A8 | Q9HAW9 | ENSG00000242366 | UDP-glucuronosyltransferase 1A8 | Transferase | Glucuronic acid (GlcA) | Glucuronidation |
| 540 | UGT1A9 | O60656 | ENSG00000241119 | UDP-glucuronosyltransferase 1A9 | Transferase | Glucuronic acid (GlcA) | Glucuronidation |
| 541 | UGT2A1 | P0DTE4 | ENSG00000173610 | UDP-glucuronosyltransferase 2A1 | Transferase | Glucuronic acid (GlcA) | Glucuronidation |
| 542 | UGT2A2 | P0DTE5 | ENSG00000271271 | UDP-glucuronosyltransferase 2A2 | Transferase | Glucuronic acid (GlcA) | Glucuronidation |
| 543 | UGT2A3 | Q6UWM9 | ENSG00000135220 | UDP-glucuronosyltransferase 2A3 | Transferase | Glucuronic acid (GlcA) | Glucuronidation |
| 544 | UGT2B10 | P36537 | ENSG00000109181 | UDP-glucuronosyltransferase 2B10 | Transferase | Glucuronic acid (GlcA) | Glucuronidation |
| 545 | UGT2B11 | O75310 | ENSG00000213759 | UDP-glucuronosyltransferase 2B11 | Transferase | Glucuronic acid (GlcA) | Glucuronidation |
| 546 | UGT2B15 | P54855 | ENSG00000196620 | UDP-glucuronosyltransferase 2B15 | Transferase | Glucuronic acid (GlcA) | Glucuronidation |
| 547 | UGT2B17 | O75795 | ENSG00000197888 | UDP-glucuronosyltransferase 2B17 | Transferase | Glucuronic acid (GlcA) | Glucuronidation |
| 548 | UGT2B28 | Q9BY64 | ENSG00000135226 | UDP-glucuronosyltransferase 2B28 | Transferase | Glucuronic acid (GlcA) | Glucuronidation |
| 549 | UGT2B4 | P06133 | ENSG00000156096 | UDP-glucuronosyltransferase 2B4 | Transferase | Glucuronic acid (GlcA) | Glucuronidation |
| 550 | UGT2B7 | P16662 | ENSG00000171234 | UDP-glucuronosyltransferase 2B7 | Transferase | Glucuronic acid (GlcA) | Glucuronidation |
| 551 | UGT3A1 | Q6NUS8 | ENSG00000145626 | UDP glycosyltransferase 3 family, polypeptide A1 | Transferase | Glucuronic acid (GlcA) | Glucuronidation |
| 552 | UGT3A2 | Q3SY77 | ENSG00000168671 | UDP glycosyltransferase 3 family, polypeptide A2 | Transferase | Glucuronic acid (GlcA) | Glucuronidation |
| 553 | UGT8 | Q16880 | ENSG00000174607 | UDP glycosyltransferase 8 | Transferase | Galactose | Galactose to ceramide |
| 554 | UST | Q9Y2C2 | ENSG00000111962 | Uronyl-2-sulfotransferase | Transferase | Sulfate | Sulfate group |
| 555 | UXS1 | Q8NBZ7 | ENSG00000115652 | UDP-glucuronic acid decarboxylase 1 | Miscellaneous | Glucuronic acid (GlcA) | Decarboxylation of UDP-glucuronic acid to UDP-xylose |
| 556 | VCAM1 | P19320 | ENSG00000162692 | Vascular cell adhesion molecule 1 | Glycan Binding/lectin | More than one monosacchrides | α4β1 (VLA-4) and α4β7integrins |
| 557 | VCAN | P13611 | ENSG00000038427 | Versican | Glycan Binding/lectin | Hyaluronic acid | Hyaluronic acid |
| 558 | WBSCR17/ GALNT17 | Q6IS24 | ENSG00000185274 | Williams-Beuren syndrome chromosome region 17 | Transferase | N-acetylgalactosamine (GalNAc) | N-acetyl-D-galactosamine residue to a serine or threonine residue on the protein receptor |
| 559 | XXYLT1 | Q8NBI6 | ENSG00000173950 | Xyloside xylosyltransferase 1 | Transferase | Xylose | Addition of the second xylose, elongates the O-linked xylose-glucose disaccharide attached to EGF-like repeats |
| 560 | XYLT1 | Q86Y38 | ENSG00000103489 | Xylosyltransferase I | Transferase | Xylose | D-xylose to serine residues of the core protein |
| 561 | XYLT2 | Q9H1B5 | ENSG00000015532 | Xylosyltransferase II | Transferase | Xylose | D-xylose to serine residues of the core protein |
